# Distinct genetic determinants and mechanisms of SARS-CoV-2 resistance to remdesivir

**DOI:** 10.1101/2022.01.25.477724

**Authors:** Laura J. Stevens, Andrea J. Pruijssers, Hery W. Lee, Calvin J. Gordon, Egor P. Tchesnokov, Jennifer Gribble, Amelia S. George, Tia M. Hughes, Xiaotao Lu, Jiani Li, Jason K. Perry, Danielle P. Porter, Tomas Cihlar, Timothy P. Sheahan, Ralph S. Baric, Matthias Götte, Mark R. Denison

**Affiliations:** Department of Pediatrics, Vanderbilt University Medical Center, Nashville, TN, 37232, USA; Vanderbilt Institute for Infection, Immunology, and Inflammation, Nashville, TN, 37232, USA; Department of Medical Microbiology and Immunology, University of Alberta, Edmonton, AB, T6G 2R3, CA; Department of Pathology, Microbiology, and Immunology, Vanderbilt University Medical Center, Nashville, TN, 37232, USA; Gilead Sciences, Inc, Foster City, CA, 94404, USA; Department of Epidemiology, University of North Carolina at Chapel Hill, Chapel Hill, NC, 27599, USA

**Author notes:** Corresponding author: Mark R. Denison. Authors contributed equally (Laura J. Stevens, Andrea J. Pruijssers).

## Abstract

The nucleoside analog remdesivir (RDV) is an FDA-approved antiviral for the treatment of SARS-CoV-2 infections, and as such it is critical to understand potential genetic determinants and barriers to RDV resistance. In this study, SARS-CoV-2 was subjected to 13 passages in cell culture with increasing concentrations of GS-441524, the parent nucleoside of RDV. At passage 13 the RDV resistance of the lineages ranged from 2.7-to 10.4-fold increase in EC_50_. Sequence analysis of the three lineage populations identified non-synonymous mutations in the nonstructural protein 12 RNA-dependent RNA polymerase (nsp12-RdRp): V166A, N198S, S759A, V792I and C799F/R. Two of the three lineages encoded the S759A substitution at the RdRp Ser_759_-Asp-Asp active motif. In one lineage, the V792I substitution emerged first then combined with S759A. Introduction of the S759A and V792I substitutions at homologous nsp12 positions in viable isogenic clones of the *betacoronavirus* murine hepatitis virus (MHV) demonstrated their transferability across CoVs, up to 38-fold RDV resistance in combination, and a significant replication defect associated with their introduction. Biochemical analysis of SARS-CoV-2 RdRp encoding S759A demonstrated a ∼10- fold decreased preference for RDV-triphosphate (RDV-TP) as a substrate, while nsp12-V792I diminished the UTP concentration needed to overcome the template-dependent inhibition associated with RDV. The *in vitro* selected substitutions here identified were rare or not detected in the >6 million publicly available nsp12-RdRp consensus sequences in the absence of RDV selection. The results define genetic and biochemical pathways to RDV resistance and emphasize the need for additional studies to define the potential for emergence of these or other RDV resistance mutations in various clinical settings.

**One Sentence Summary:** SARS-CoV-2 develops in vitro resistance to remdesivir by distinct and complementary mutations and mechanisms in the viral polymerase

## Introduction

SARS-CoV-2 infections have caused more than 850,000 deaths in the United States and over 5 million deaths worldwide(*1, 2*). Remdesivir (RDV, GS-5734) is the first FDA approved direct-acting antiviral for the treatment of SARS-CoV-2. RDV is a monophosphoramidate prodrug of the C-adenosine analog GS-441524 that acts by inhibiting RNA synthesis by the viral RNA-dependent RNA polymerase (nsp12-RdRp), and has been shown to be broadly active against multiple RNA viruses(*3–7*). Preferential incorporation of the triphosphate form of RDV (RDV-TP) over its natural ATP nucleotide counterpart results in inhibition of RNA synthesis via several mechanisms. Delayed chain termination may occur three nucleotides following RDV-TP incorporation due to a clash of the RDV-monophosphate (MP) 1’-cyano with S861 in the RdRp RNA exit channel, thereby preventing further enzyme translocation(*8–11*). However, increases in NTP concentrations can overcome this obstacle and RNA synthesis can continue, resulting in RNA strands with incorporated RDV-MP residues. In this setting, template-dependent inhibition of RNA synthesis occurs because of compromised incorporation of the complementary UTP opposite RDV-MP(*12, 13*).

The use of therapeutic RDV has been shown to improve disease outcomes and reduce viral loads in SARS-CoV-infected mice, in mice infected with chimeric SARS-CoV encoding the SARS-CoV-2 RdRp, in SARS-CoV-2 infected mice co-treated with therapeutic antibodies, and in infected rhesus macaques(*4, 7, 14–16*). Furthermore, RDV potently inhibits viral replication of both human endemic CoVs and bat CoVs in primary human lung cell cultures(*4, 6*). A large double-blinded, randomized, placebo-controlled trial of intravenous RDV in adult patients hospitalized with COVID-19 demonstrated that RDV was superior to placebo in shortening time to recovery(*17, 18*) and recent data suggests 50% increased survival rates if given early in infection(*19*). In addition, a 3-day early treatment course of RDV reduced the hospitalization of high-risk COVID-19 patients by 87% compared to placebo(*20*). However, a recent case report described the emergence of possible RDV resistance in an immunocompromised patient, underscoring the importance of further understanding pathways to RDV resistance(*21*). Little is known about the evolution, viral determinants, and specific mechanisms of SARS-CoV-2 resistance to RDV, limiting active surveillance for resistance-associated substitutions.

Here we report multiple pathways by which SARS-CoV-2 achieved varying degrees of resistance to RDV during serial passage in cell culture in the presence of GS-441524, the parent nucleoside of RDV, including multiple combinations of nsp12-RdRp amino acid substitutions including: V166A, S759A, V792I, and C799F/R. Lineages containing S759A demonstrated 7-to- 9-fold decreased sensitivity to RDV by EC_50_. Introduction of the SARS-CoV-2 co-selected S759A and V792I mutations at identical nsp12-RdRp residues in the *Betacoronavirus*, murine hepatitis virus (MHV), conferred up to 38-fold increase in RDV EC_50_ but also incurred a replication defect compared to WT virus. Biochemical analyses of SARS-CoV-2 nsp12 with S759A and V792I mutations demonstrated distinct and complementary molecular mechanisms of RDV resistance. This study provides important insights into potential evolutionary pathways leading to RDV resistance, identifies viral determinants and molecular mechanisms of RDV resistance, and forms the basis for surveillance for early indicators for potential RDV resistance.

## Results

### SARS-CoV-2 acquires phenotypic resistance to RDV during passage with GS-441524

We previously reported that RDV resistance mutations selected in MHV conferred resistance in SARS-CoV(*5*). To identify viral genetic pathways to RDV resistance in SARS-CoV-2, we passaged the WA-1 clinical isolate (MN985325)(*22*) in Vero E6 cells in the presence of DMSO vehicle or GS-441524, which is metabolized to the same active nucleoside triphosphate as the prodrug RDV, is able to achieve higher concentrations of active triphosphate than RDV in Vero E6 cells(*7*). Virus was passaged in three independent and parallel series, resulting in three GS-441524-passaged lineages and three DMSO-passaged lineages (Fig. S1). An increase in cytopathic effect (CPE) was observed in the all three GS-441524-passaged lineages between passages 10 and 13. To determine whether this shift represented selection for resistance, we tested the sensitivity of each drug- and DMSO-passaged lineage to RDV in the human lung cell line, A549-hACE2, at passage 9 (P9) and passage 13 (P13) by quantifying the relative change in viral genome copy number in cell culture supernatant (Fig. 1, Table S1). All three P9 GS-441524- passaged lineages were less sensitive to RDV than P9 DMSO lineages. GS-441524 lineage 1 appeared moderately less sensitive to RDV, with a 2.6-fold increase in EC_50_ at P9. Lineage 1 RDV sensitivity decreased further by P13, with a 10.4-fold increase in EC_50_. GS-441524 lineage 2 demonstrated minimal change in susceptibility to RDV at P9, with a 1.5-fold increase in EC_50_ compared to DMSO-passaged lineages and its sensitivity further decreased modestly by P13, with a 2.7-fold increase in EC_50_ compared to DMSO-passaged lineages. At P9, GS-441524 lineage 3 demonstrated a 1.7-fold increase in EC_50_ compared to the DMSO-passaged lineages. Sensitivity of GS-441524 lineage 3 decreased further by P13, with an 8-fold increase in EC_50_ compared to DMSO-passaged lineage 1. We next tested the replication of all the DMSO and GS-441524- passage lineages in the absence of RDV. Compared to the clear replication advantage observed in the presence of RDV, all three GS-441524-passaged P13 lineages demonstrated delayed and impaired replication with ∼0.5-1 log decrease in peak titer compared to DMSO-passaged control virus in the absence of drug. Thus, selection for phenotypic resistance conferred a replication defect in all three lineages.

**Fig. 1.**
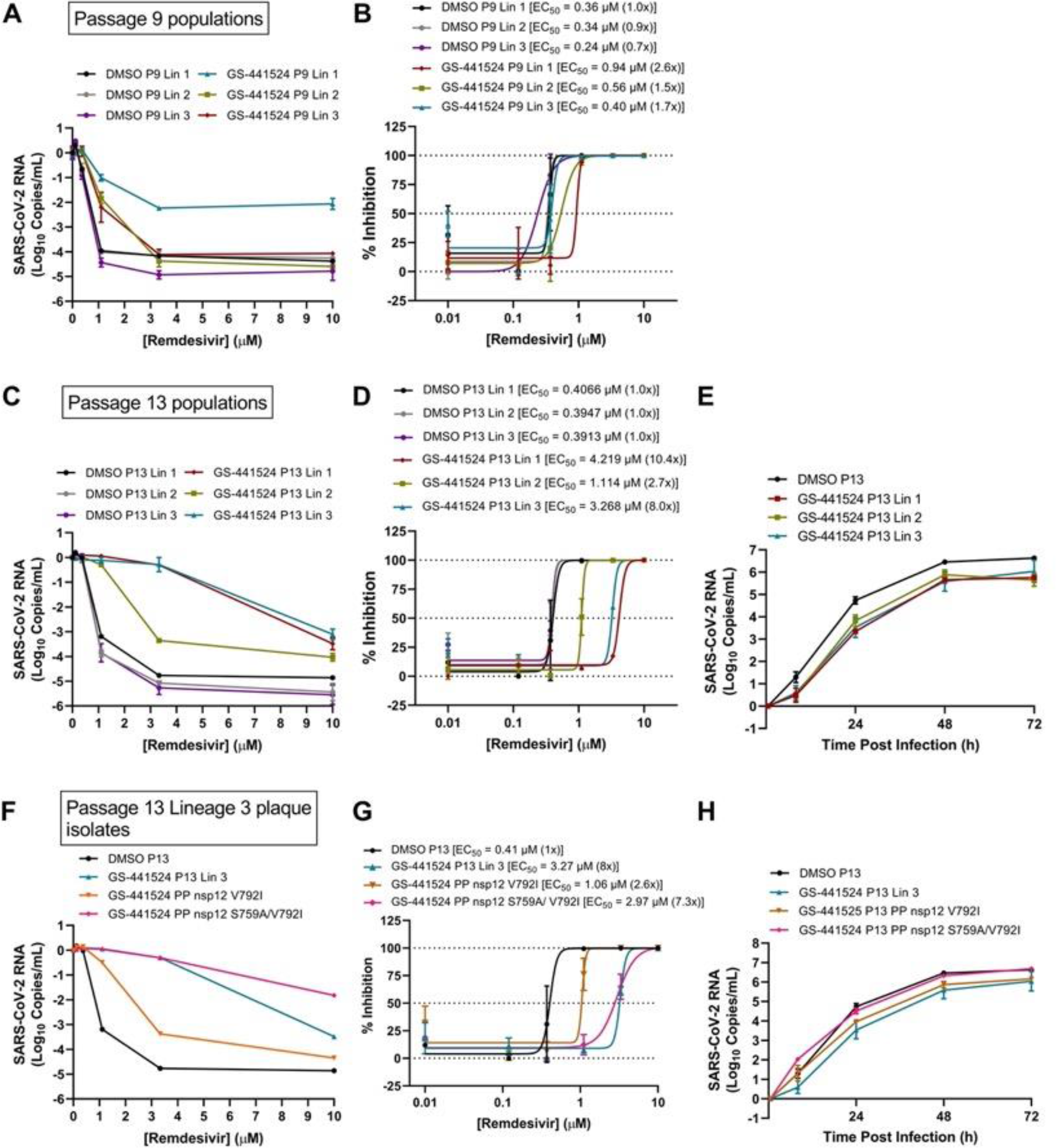
(See also Data file S1) SARS-CoV-2 RDV resistance and replication after serial passaging. SARS-CoV-2 was serially passaged in the presence and absence of GS-441524 in Vero-E6 cells in triplicate lineages. **(A)** Sensitivity of P9 lineages to RDV in A549-hACE2 cells as determined by change in genome copy number. **(B)** Percent inhibition calculated from genome copy number *(A)* and fold-change in EC50 compared to vehicle (DMSO)-passaged lineage 1 at P9. **(C)** Sensitivity of P13 lineages to RDV in A549-hACE2 cells as determined by change in genome copy number. **(D)** Percent inhibition calculated from genome copy number *(C)* and fold- change in EC50 compared to vehicle-passaged lineage 1 at P13. **(E)** Replication kinetics of P13 drug-passaged viral lineages compared to vehicle-passaged lineage 1. **(F)** Sensitivity to RDV of plaque-pick (PP) isolates from GS-441524-passaged lineage 3 and vehicle-passaged lineage 1 population viruses and input virus in A549-hACE2 cells as determined by change in genome copy number. Plaque-picks (PP) from lineage 3 were isolated and expanded in presence of 1uM GS- 441524. **(G)** Percent inhibition calculated from raw genome copy number *(F)* and fold-change in EC50 compared to vehicle DMSO-passaged lineage 1. **(H)**Replication kinetics of plaque isolates tested in *(F)* and *(G)*.

**Table 1.**
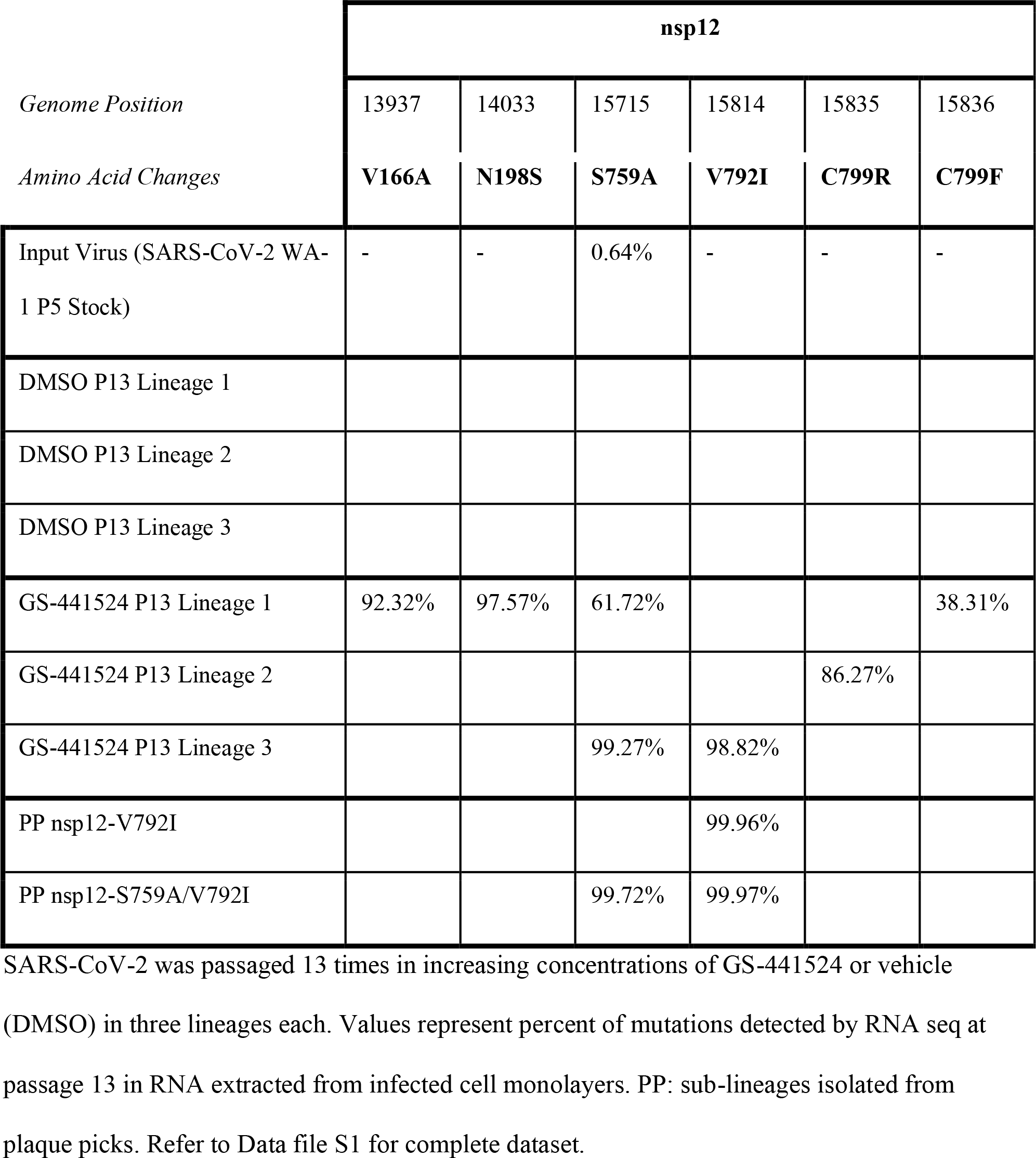
nsp12 non-synonymous mutations present at >15% of populations of serially passaged SARS-CoV-2.

### Identification of candidate resistance mutations in nsp12-RdRp

To identify candidate resistance mutations, we performed short-read Illumina poly(A) RNA-sequencing (RNA-seq) on RNA purified from infected cell monolayers of all six lineages at passages 0 (input), 6, 9, and 13. (Table 1, Data file S1). Numerous low frequency nucleotide mutations (0.01 – 5%) were detected in all six lineages at P13 (Data file S1). Of the non-synonymous (NS) mutations present at >15% frequency, mutations in spike were present in both DMSO- and drug-passaged lineages and likely represent cell culture adaptation (Data file S1). In contrast, six NS nsp12-RdRp mutations were detected with >15% frequency in GS-441524- passaged lineages but not in any of the DMSO-passaged lineages (Table 1). By P13, these nsp12 NS mutations were dominant (>50%) in GS-441524-passaged populations. GS-441524 lineage 1 encoded NS nsp12-RdRp mutations V166A (92%), N198S (97%), S759A (62%) and C799F (38%); lineage 2 encoded only C799R; and lineage 3 encoded S759A (99%) and V792I (99%).

GS-441524 lineage 1 and 3 populations were more resistant than GS-441524 lineage 2 based on both extent of CPE and the increased RDV EC_50_ (Fig. S1, Fig. 1). Of the selected substitutions, only S759A was detected in the original SARS-CoV-2 WA-1 P5 stock virus population at 0.64%. The S759A was not detected at any level in any of the lineages passaged in the DMSO vehicle (Table 1, Data file S1), suggesting a lack of positive selection in absence of RDV. Overall, these results identified distinct combinations of a limited number of nsp12-RdRp NS mutations associated with independent RDV-resistant lineages.

To look for presence of the identified *in vitro* GS-441524-selected nsp12 mutations in circulating clinical isolates we analyzed the consensus sequences of >6 million clinical isolates of SARS-CoV-2 submitted to the GISAID Database(*23*) prior to January 4, 2021 (Table S2). In the absence of RDV selection, the S759A mutation was identified in submitted consensus sequences, including the Delta variant only in a single isolate while the other nsp12 mutations were detected at a frequency less than 0.02%, and none of the nsp12 mutations were observed among >130,000 Omicron variant consensus sequences submitted as of this writing. A clear limitation of this dataset is that the details of isolation and raw sequence data are not available. Consensus sequences most likely represent nucleotides present at >50% and that any single nucleotide polymorphisms present <50% in the population would not be represented in this analysis. For example, we detected the S759A substitution at 0.64% in the expanded clinical WA-1 isolate received from the CDC, which may have importance for its eventual selection. Similarly, in the GS-441524 lineage 1 passage 13, C799F was present at 38%, a level that would not be reported in consensus sequence. Despite this limitation, the analysis can allow us to conclude that in the absence of RDV-selective pressure the *in vitro* identified nsp12 mutations were not present as dominant variants or propagated in circulating SARS-CoV-2 including Delta and Omicron variants. This supports our hypothesis that the identified nsp12 mutations likely were associated with GS-441524 selective pressure, a hypothesis we next tested.

**Table 2.** Selectivity values against RDV-TP incorporation by SARS-CoV-2 WT and a set of mutants containing single amino acid substitution at residue S759

| RdRp | $V_{\max}^a$ | | $K_m^b$ | | $V_{\max} / K_m$ | | Selectivity <sup>c</sup> | Discrimination <sup>d</sup> |
| --- | --- | --- | --- | --- | --- | --- | --- | --- |
|  | ATP | RDV-TP | ATP | RDV-TP | ATP | RDV-TP |  |  |
| **WT**<br>n=10 <sup>e</sup> | 0.86<br>±0.016 <sup>f</sup> | 0.86<br>±0.015 | 0.058<br>±0.0041 | 0.022<br>±0.0015 | 15 | 39 | 0.38 | Ref. <sup>g</sup> |
| **S759A**<br>n=3 | 0.92<br>±0.008 | 0.91<br>±0.006 | 0.19<br>±0.0075 | 0.74<br>±0.025 | 4.8 | 1.2 | 4 | 10.5 |
| **V792I**<br>n=3 | 0.87<br>±0.039 | 0.90<br>±0.031 | 0.017<br>±0.0040 | 0.023<br>±0.0035 | 51 | 39 | 1.3 | 3.4 |

<sup>a</sup> $V_{\max}$ is a Michaelis–Menten parameter reflecting the maximal velocity of nucleotide incorporation; reported as a raw value of product fraction of the incorporated nucleotide. <sup>b</sup> $K_m$ is a Michaelis–Menten parameter reflecting the concentration ( $\mu\text{M}$ ) of the nucleotide substrate at which the velocity of nucleotide incorporation is half of $V_{\max}$ . <sup>c</sup> Selectivity of a viral RNA polymerase for a nucleotide substrate analogue is calculated as the ratio of the $V_{\max}/K_m$ values for NTP and NTP analogue, respectively; as such, it is a unitless value. <sup>d</sup> The discrimination index is calculated as a ratio of selectivity of the mutant over wild type. <sup>e</sup> All reported values have been calculated on the basis of a 9-data point experiment repeated the indicated number of times (n). <sup>f</sup> Standard error associated with the fit. <sup>g</sup> Reference.

### nsp12-S759A and -V792I are associated with in vitro RDV resistance

The S759A residue substitution was selected in GS-441524 lineages 1 and 3, which demonstrated the most CPE and EC_50_ increase of GS-441524-passaged lineages. In both lineages, the S759A substitution emerged as a dominant change in the population in combination with at least one other substitution; GS-441524 lineage 3 contained only S759A and V792I. To define the contribution of S759A and V792I to RDV resistance in the context of adapted infectious virus, we plaque-picked (PP), expanded, and sequenced sub-lineages derived from GS-441524-passaged lineage 3 (Table 1, Data file S1). We isolated a sub-lineage containing V792I alone and another sub-lineage containing S759A in combination with V792I. Importantly, in these two sub-lineages, no other NS mutations were detected at >5% elsewhere in replicase genes (Data file S1). In RDV sensitivity assays, the S759A/V792I-containing sub-lineage was indistinguishable from the GS- 441524-passaged lineage 3 population at RDV concentrations up to 3.3μM, with a 7.3-fold increase in EC_50_ over the DMSO-passaged control as compared to an 8-fold increase in EC_50_ for the GS-441524 lineage 3 P13 population virus (Fig.1, Table S1). Thus, the S759A/V792I plaque-picked sub-lineage recapitulated resistance phenotype of the GS-441524 lineage 3 P13 population. In contrast, the V792I-containing sub-lineage displayed a low-level resistance phenotype corresponding to a 2.6-fold increase in EC_50_ compared to the DMSO-passaged virus.

Finally, in the absence of RDV, the V792I and S759A/V792I plaque-picked sub-lineages demonstrated replication kinetics similar to the DMSO-passaged control virus rather than to the GS-441524 lineage 3 population virus (Fig. 1H), suggesting differential replication efficiency among subpopulations of GS-441524-passaged virus.

### Selection and emergence of candidate resistance mutations and combinations

To define viral genetic progression toward resistance, we quantified the relative frequency of RDV resistance-associated nsp12 mutations in the GS-441524 lineages 1-3 at P6, P9, and P13 by RNA-seq (Fig.2 A-C) and direct nanopore MinION sequencing of DNA amplicons spanning nsp12 coding domain (Fig.2 D-F). The data results between the two methods were consistent in the emergence and patterns. In lineage 1, N198F was present in the population at >87% by P6 and thereafter, while V166A, S759A, and C799F were less abundant at P6 but by P13 were prominent. In lineage 2, only C799R was detected at any passage by RNA-seq or nanopore amplicon sequencing. In lineage 3, V792I became dominant in the P9 population, while S759A was detected at ∼2.5% by nanopore amplicon sequencing at P6 and P9, but by P13 was >65%. In both lineage 1 and 3, emergence of S759A correlated with an increase in EC_50_ (2.6-to-10.4-fold in lineage 1; 1.7-to-8.0-fold in lineage 3).

**Fig. 2.**
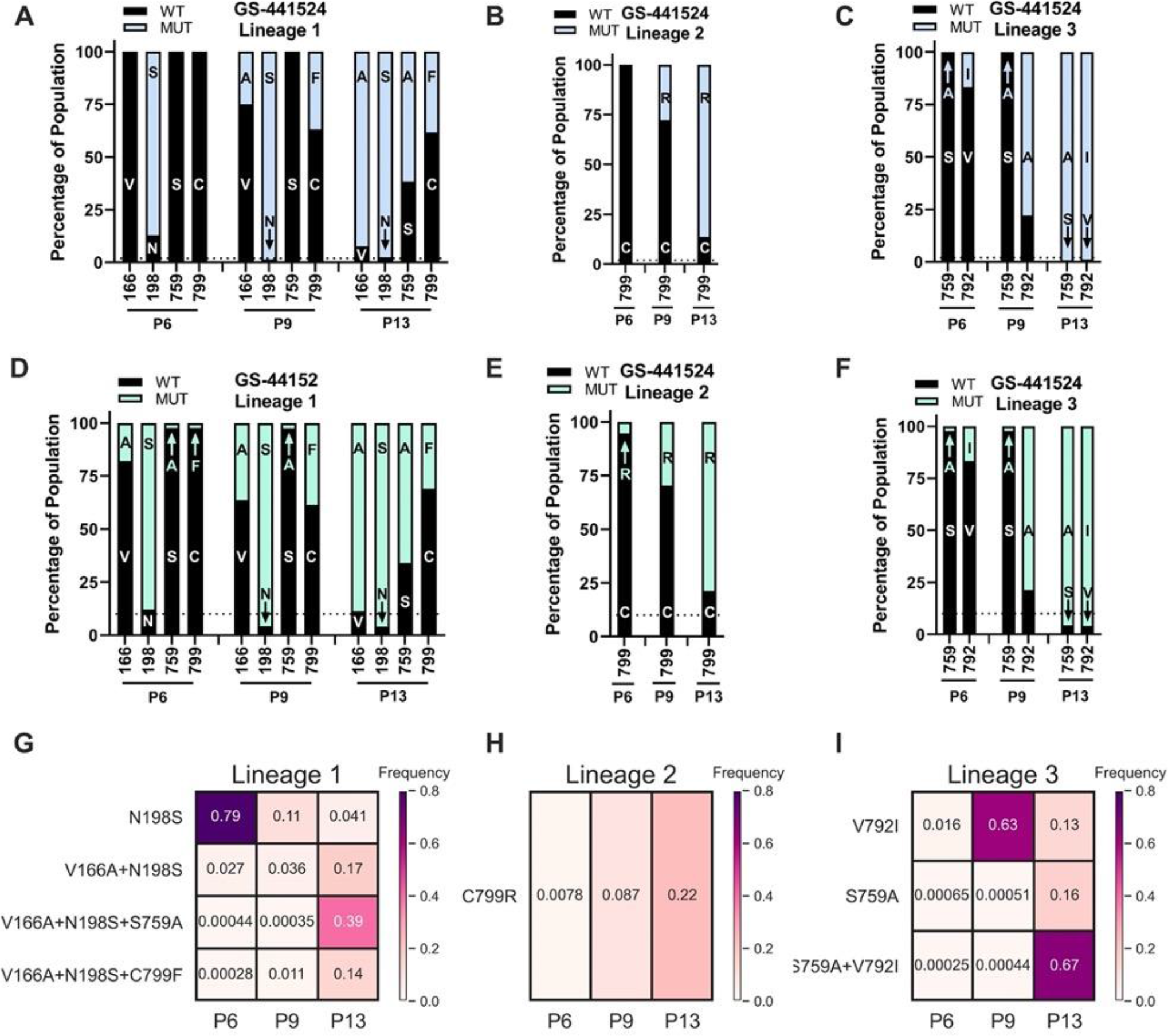
Evolution and intramolecular linkage of nsp12 mutations. SARS-CoV-2 was passaged 13 times in increasing concentrations of GS-441524 in 3 lineages. RNA from infected cell monolayers was subjected to Illumina RNA sequencing and Oxford nanopore MinION sequencing. **(A,B,C) RNA-seq percent of nsp12 mutations** in lineages 1, 2 and 3. **(D,E,F) Nanopore amplicon sequencing percent of nsp12 mutations** in lineages 1, 2 and 3) **(G,E,H) Frequency of single and combined sets of nsp12 mutations** in single viral genomes in lineages 1, 2, and 3. Variants were mapped according to their genomic position and frequency, expressed as a percentage of the total reads mapped to that position.

To determine the frequency with which resistance-associated mutations were present individually or in haplotypic linkages, we employed long-read nanopore (MinION) sequencing across full-length nsp12 amplicons and developed the bioinformatics pipeline, *MutALink*, to quantify the absolute abundance of each mutation alone and in combination with other mutations that occurred at >15% frequency within by P13 (Fig. 2 G-I). Of the nsp12-RdRp amino acid substitutions identified in lineage 1, the abundance of combined V166A/N198S/S759A underwent the greatest increase with increasing drug concentration. In lineage 2, the C799R mutation had no detected linkage to another mutation. In lineage 3, the abundance of combined S759A/V792I demonstrated the greatest increase with increasing drug concentration (39%). Thus, S759A predominantly existed in combination with either N198S+V166A (lineage 1) or V792I (lineage 3).

### Structural modeling of S759A predicts altered RDV interactions

The potential of RDV resistance-associated nsp12 mutations to accommodate the RDV-TP substrate was evaluated using a structural model of the pre-incorporation state in which the NTP substrate is base paired to the template and coordinated with two Mg_++_ ions in the active site (Fig. 3). The model was generated based on the SARS-CoV-2 RdRp cryo-EM structure 6XEZ(*24*), but was influenced by several other polymerase structures(*25–27*). From this model, we determined that the amino acid substitutions which arose during serial drug passage were clustered in three general locations: S759A was located in the RdRp catalytic S_759_DD motif C, in close proximity to the incoming NTP; V166A, V792I and C799F/R were located on or adjacent to motif D, a common structural element of polymerases important to the dynamics of NTP incorporation; and N198S was located on the protein surface behind the NiRAN site, a position having no known or predicted role in RNA synthesis. Focusing on the active site, several polar residues form a pocket containing T687, A688 and N691 on Motif B and S759 on motif C that can easily accommodate the RDV-TP 1’-cyano group. Some variation in the sidechain conformations of T687 and S759 was observed across the available cryo-EM structures and may be dependent on the state of RNA and substrate binding. A computational assessment of these conformations within the RDV-TP pre-incorporation model suggested that a state which oriented the hydroxyls toward the RDV-TP 1’-cyano was optimal. The resulting hydrogen bonds were weak (2.5 Å for T287 and 2.8 Å for S759), but this overall favorable interaction suggested a potential advantage of RDV-TP over natural substrates. Modeling the S759A mutation showed a loss of one of these hydrogen bonds, and from that we predicted decreased binding affinity of RDV-TP in the pre-incorporation complex relative to ATP. The effect of other observed mutations was more difficult to predict structurally. A conformational search of the nsp12 loops 163-168 and 790-800 for both WT and mutant amino acids predicted only modest shifts for V166A+S759A and S759A+V792I. In contrast, the C799R and V166A+C799F mutations predicted clear changes in the conformation of motif D. As motif D is thought to be important to the closing of the polymerase active site once the NTP is positioned for incorporation(*28*), any mutation affecting dynamics of this loop could potentially impact incorporation rates.

**Fig. 3.**
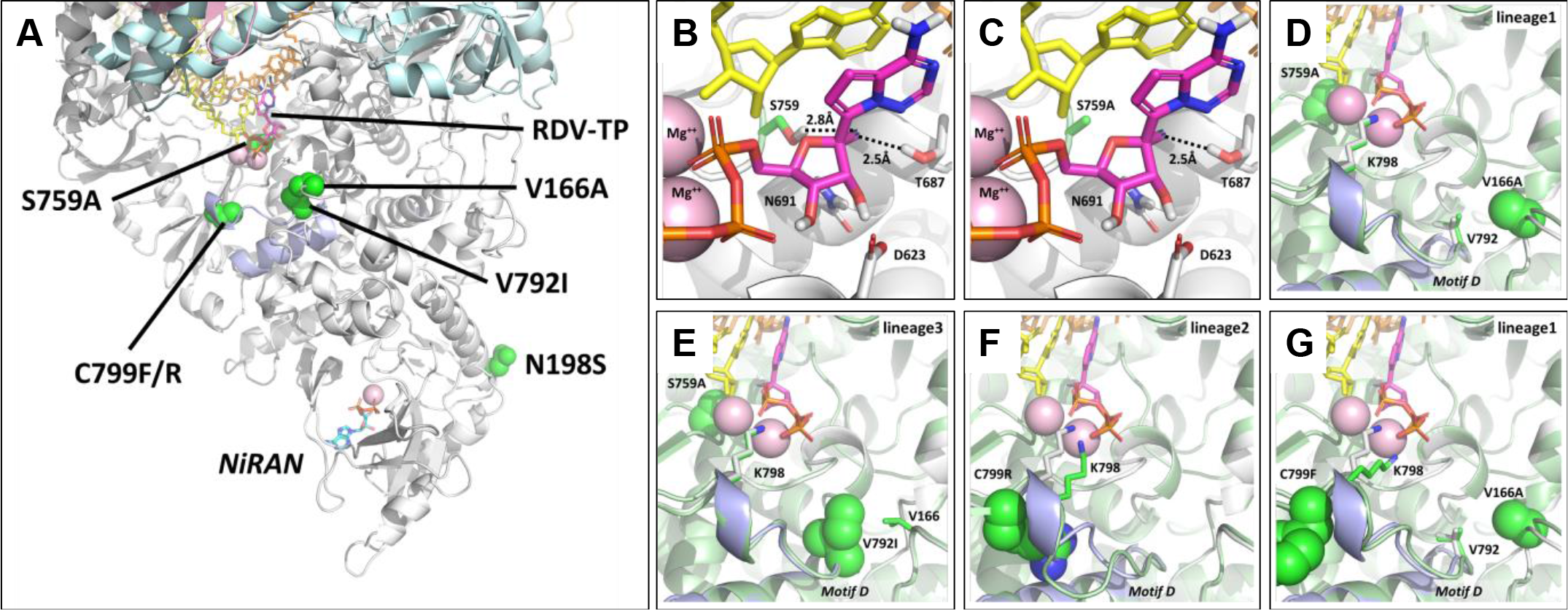
Structural modelling predictions. **(A)** Map of observed nsp12 amino acid substitutions on a model of the SARS-CoV-2/RDV-TP pre-incorporation complex. nsp12 is shown in white, nsp7 in pink, nsp8 in cyan, the primer strand in yellow, the template strand in orange, RDV-TP in magenta, and mutations in green. S759A is in the active site, while V166A, V792A and C799F/R are adjacent to the active site, clustered around Motif D (in blue). N198S does not appear to impact either the NiRAN or Pol sites. **(B, C)** Detail of the RDV-TP pre-incorporation model, highlighting the polar residues that interact with the 2’OH and the 1’CN. S759 is seen to be in close contact with the 1’CN, forming a favorable interaction that is lost with the S759A mutation. **(D)** Model of the lineage 1 mutations V166A and S759A (green) overlaid on the WT structure (white). V166A is in direct contact with V792 and may impact the dynamics of Motif D. **(E)** Model of the similar lineage 3 mutations V792I and S759A (green) overlaid on the WT structure (white). **(F)** Model of the lineage 2 mutation C799R (green) overlaid on the WT structure (white). The mutation is predicted to alter the conformation of Motif D, impacting how K798 interacts with the substrate γ-phosphate. **(G)** Model of the lineage 1 mutations V166A and C799F, which are also seen to alter the conformation of Motif D and the position of K798.

### SARS-CoV-2 S759A and V792I mutation homologues confer RDV resistance in recombinant Betacoronavirus murine hepatitis virus

In all subsequent experiments we targeted S759A and V792I for genetic and biochemical analysis, since: 1) they were associated with the most increase in measured resistance; 2) they could be isolated alone (V792I) and together (S759A/V792I) in virus clones; and 3) S759A, located in the nsp12-RdRp LS_759_DD active motif modeled a plausible change in interaction with RNA. To directly test the capacity of S759A and V792I substitutions to mediate RDV resistance in isogenic virus backgrounds devoid of other mutations, we engineered individual S-to-A and V-to-I changes at the aligned identical and structurally orthologous S755 and V788 residues in the nsp12-RdRp of the *Betacoronavirus* murine hepatitis virus (MHV) (Fig. 4). Viable MHV mutants encoding S755A, V788I, or S755A+V788I were compared with WT MHV in replication assays in the absence of RDV. While all mutant viruses ultimately attained peak titers similar to WT, there were significant defects in mutant virus replication kinetics. The MHV S755A and V788I mutants demonstrated a 2-hour delay to exponential replication compared to WT MHV and a 12- hour delay to peak titer. The S755A/V788I double mutant conferred a more protracted 4-hour delay to exponential replication and 16-hour delay to peak titer. Analysis of sensitivity to RDV demonstrated that MHV mutants were less sensitive to RDV than WT MHV based on reduction in infectious viral titer and genome copy number. MHV mutants encoding S755A or V788I alone demonstrated small decreases in sensitivity to RDV compared to WT MHV by EC_50_ calculation. For S755A, there was a 2.2-fold increase in EC_50_ by infectious virus titer (plaque assay) and 2.7-fold increase based on RNA genome copy number. Corresponding values for V788I were 1.3-fold and 1.8-fold increase in EC_50_, respectively. In contrast, the combined S755A+V788I mutant demonstrated a of 38.3-fold increase in EC_50_ compared to WT based on infectious viral titer and 24.9-fold increase in EC_50_ based on genome copy number. Further, the combination S755A/V788I mutant demonstrated complete resistance to RDV at a concentration of RDV that caused >99% inhibition of WT MHV (0.6 mM). Thus, GS-441524-selected SARS-CoV-2 S759A and V792I- associated phenotypes were transferable to MHV and mediated significant RDV resistance in the MHV genetic background. The results also demonstrated that the substitutions, when tested in isolation or in combination, impaired virus replication efficiency in the absence of RDV, consistent with our previous study demonstrating that RDV resistance selection in MHV was achieved at the expense of viral fitness(*5*).

**Fig. 4.**
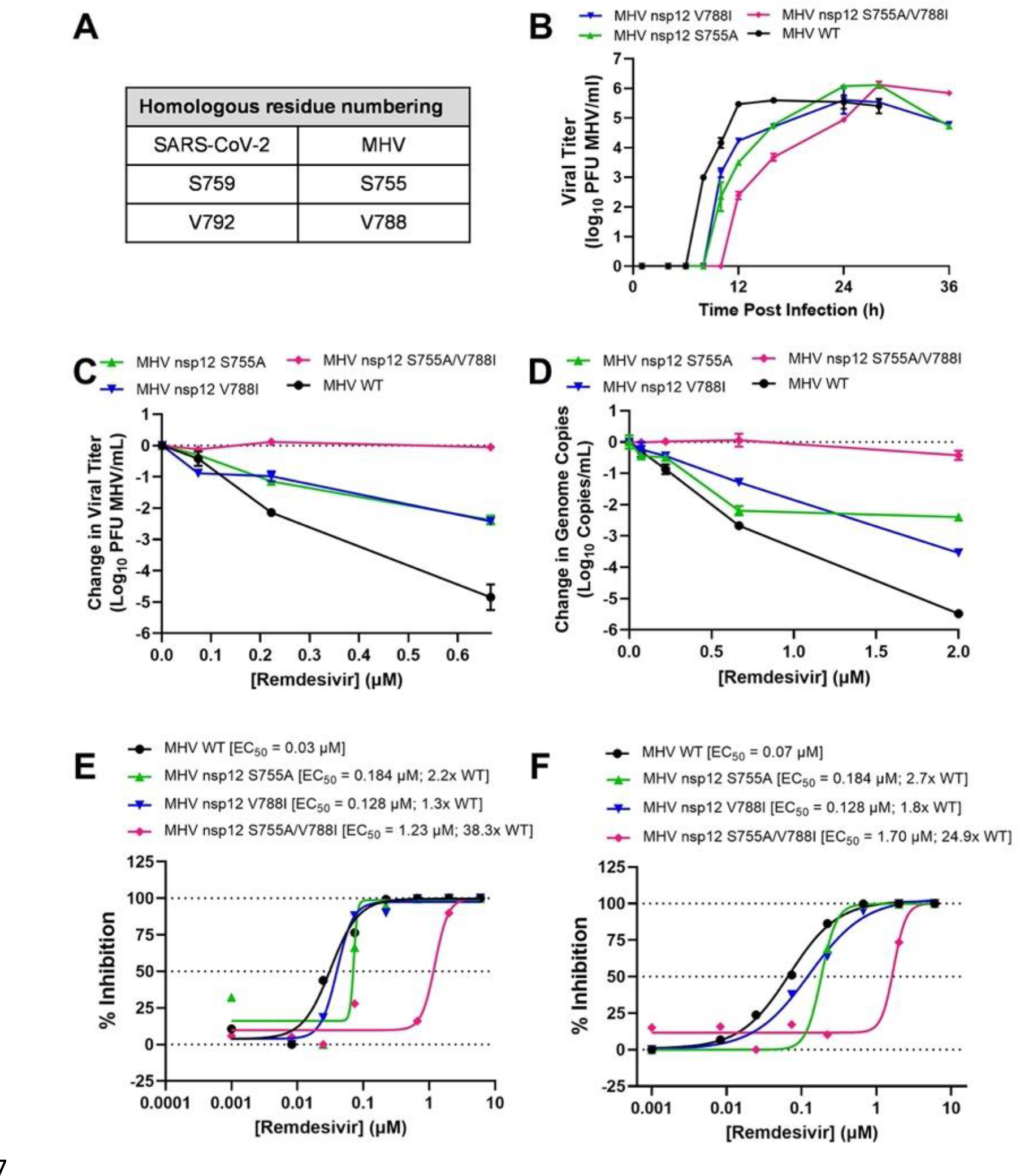
Effect of SARS-CoV-2 resistance mutations in MHV. **(A)** Candidate resistance mutations identified in SARS-CoV-2 were engineered at conserved homologous positions in the MHV infectious clone. WT MHV and mutant viruses were tested against RDV in murine delayed brain tumor (DBT9) cells. **(B)** Viral replication kinetics. **(C)** Change in infectious viral titer by plaque assay. **(D)** Change in genome copy number by qRT-PCR. **(E)** Percent inhibition and EC_50_ calculated using infectious virus titers from *(B).* **(F)** Percent inhibition calculated using genome copy numbers from (C).

### SARS-CoV-2 nsp12 S759A and V792I substitutions mediate RDV resistance by independent and complementary biochemical mechanisms

We next sought to define the mechanisms of S759A and V792I resistance. For biochemical analysis, we expressed and purified SARS-CoV-2 WT, S759A, or V792I RdRp complexes consisting of nsp7, nsp8, and nsp12, using approaches previously described(*8*). A key characteristic of RDV-TP as an inhibitor of WT SARS-CoV-2 is that it is incorporated with higher efficiency than its natural ATP counterpart by the viral RdRp(*8, 29*). To determine if a change in selective substrate usage could explain the effect of S759A and V792I, we determined the efficiency of incorporation of ATP over RDV-TP by measuring *V*_max_/*K*_m_ for single nucleotide incorporation events under steady-state conditions (Fig. 5, Table 2). ATP and RDV-TP incorporation were monitored with a model primer/template using WT RdRp and the S759A and V792I mutants. Consistent with previous reports(*8, 29*), the selectivity measured with the WT enzyme was 0.38, indicating that RDV-TP was preferred over ATP (Table 2). In contrast, the selectivity value for the S759A mutant was 4, demonstrating a preferred use of ATP over RDV-TP. This shift was driven primarily by a marked reduction in the use of RDV-TP as a substrate (∼33-fold decrease in *V*_max_/*K*_m_). The efficiency of incorporation of ATP also was compromised with S759A to a lesser degree (∼3.1-fold decrease in *V*_max_/*K*_m_). When corrected for the difference in ATP usage, the S759A mutant showed a 10.5-fold reduced use of RDV-TP as substrate relative to the use of ATP. In contrast, the RdRp expressing V792I demonstrated increased capacity to incorporate ATP (∼3-fold), while the increase in RDV-TP substrate usage was less pronounced.

**Fig. 5.**
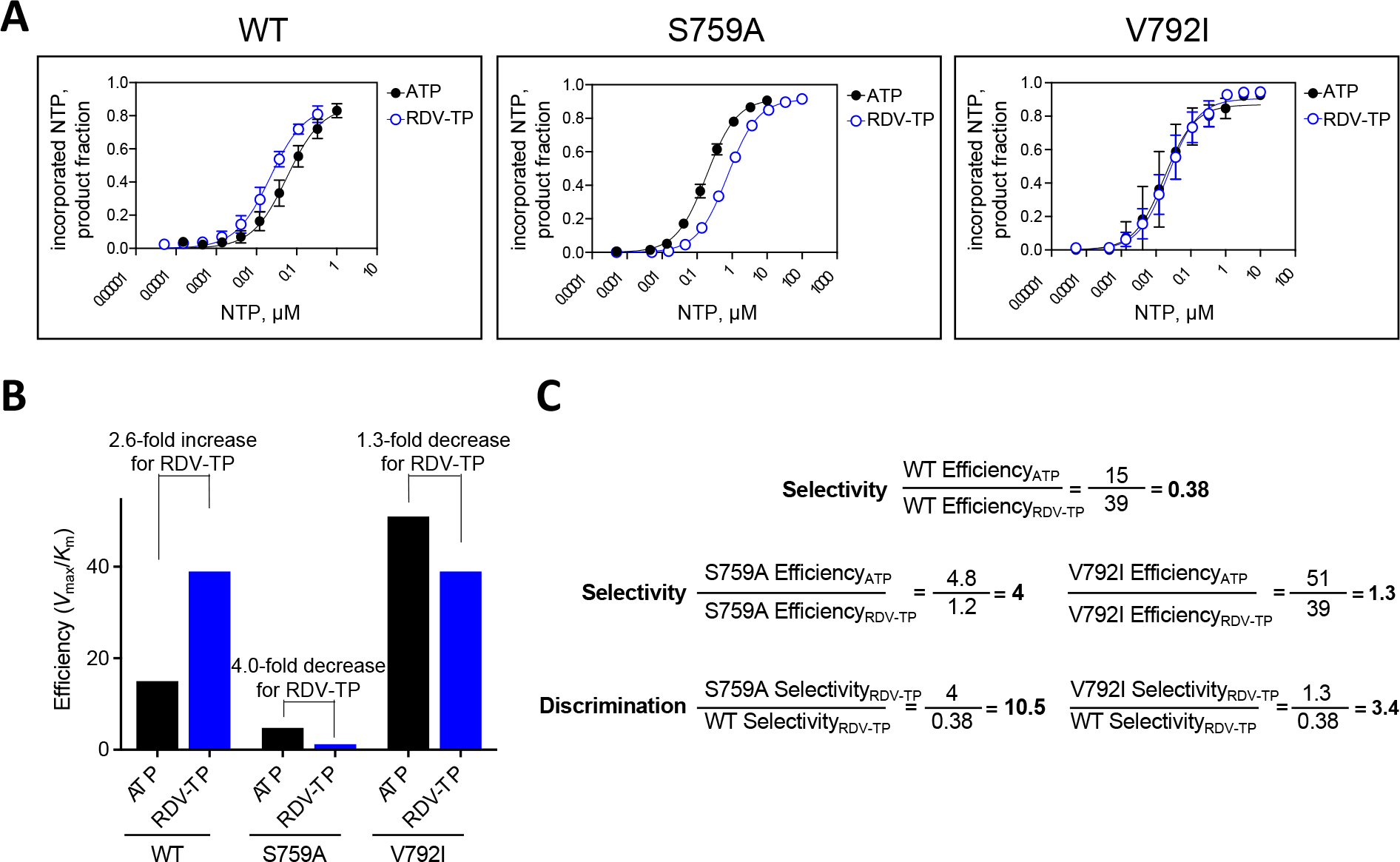
Efficiency of RDV-TP incorporation by WT, S759A, and V792I mutant SARS-CoV-2 RdRp complexes. (A) Graphical representation of the data shown in figure Fig.S2. Best fit lines illustrate fitting of the data points to Michaelis-Menten kinetics function using GraphPad prism 7.0 (GraphPad Software, Inc., San Diego, CA). Error bars illustrate standard deviation of the data. All data represent at least three independent experiments. (B) Graphic representation of efficiencies of incorporation (ATP and RDV-TP) and selectivity (ATP over RDV-TP) of the mutant enzymes corrected for differences in ATP incorporation. (C) Calculation of the discrimination value against RDV-TP across mutant enzymes.

This resulted in a selectivity value of ∼0.81, only a 2.1-fold change compared with the WT RdRp. Thus, the S759A mutant discriminated against the inhibitor RDV-TP at the level of nucleotide incorporation much more effectively than V792I.

Next, we used a second biochemical assay to assess both the effect of changes in selective incorporation of ATP versus RDV-TP as well as the inhibitory effects of the incorporated RDV-MP on primer extension. We utilized a polyU template and increasing RDV-TP concentrations to enhance the resistant phenotypes by allowing multiple incorporations of RDV-TP, which could potentially magnify effects of the mutations (Fig. S3). In this *in vitro* model for WT RdRp, initially increasing RDV-TP concentration resulted in increased chain-termination. However, further increases in RDV-TP concentration resulted in an increase in full-length product due to efficient RDV-TP substrate incorporation on the polyU template that overcame delayed chain termination. The V792I substitution showed a very similar pattern to WT with only marginal increases in full-length product formation. Conversely, both the S759A and S759A/V792I mutants demonstrated delayed chain-termination only at higher concentrations compared to WT, and no rebound in full-length product formation was observed as RDV-TP concentrations increased. S759A and S759A/V792I are indistinguishable in this assay and the phenotype is driven by S759A and the reduced usage of RDV-TP as a substrate. In contrast, the V792I substitution alone did not demonstrate a significant effect on selective incorporation of RDV-TP, delayed chain-termination, or overcoming delayed chain-termination (Table 2, Fig 5, Fig. S3).

This result indicated that the contribution of V792I to RDV resistance is likely based on a different mechanism.

We previously reported that the MHV-V553L RDV resistance mutation, when tested as the homologous V557L substitution in a SARS-CoV-2 biochemical system, conferred low-level RDV resistance by improving incorporation of UTP opposite the RDV-MP in the template and thereby reducing template-dependent inhibition(*12*). To test a potential effect of V792I, S759A, and S759A/V792I on template-dependent inhibition, we prepared WT template-A in which a single AMP was embedded, and template-R in which a single RDV-MP was embedded (Fig. 6). All enzymes were equilibrated so that the same amount of product was present in the absence of UTP (Fig. 6B, lane “0”, product 10). The S759A mutant behaved almost identically to WT in this reaction, while V792I alone or S759A/V792I together lowered the UTP concentration needed to overcome template-dependent inhibition (Fig. 6C, F). Thus, distinct and complementary mechanisms of resistance were associated with S759A and V792I and the two residue substitutions combined provided an advantage to RNA synthesis in the presence of RDV.

**Fig. 6.**
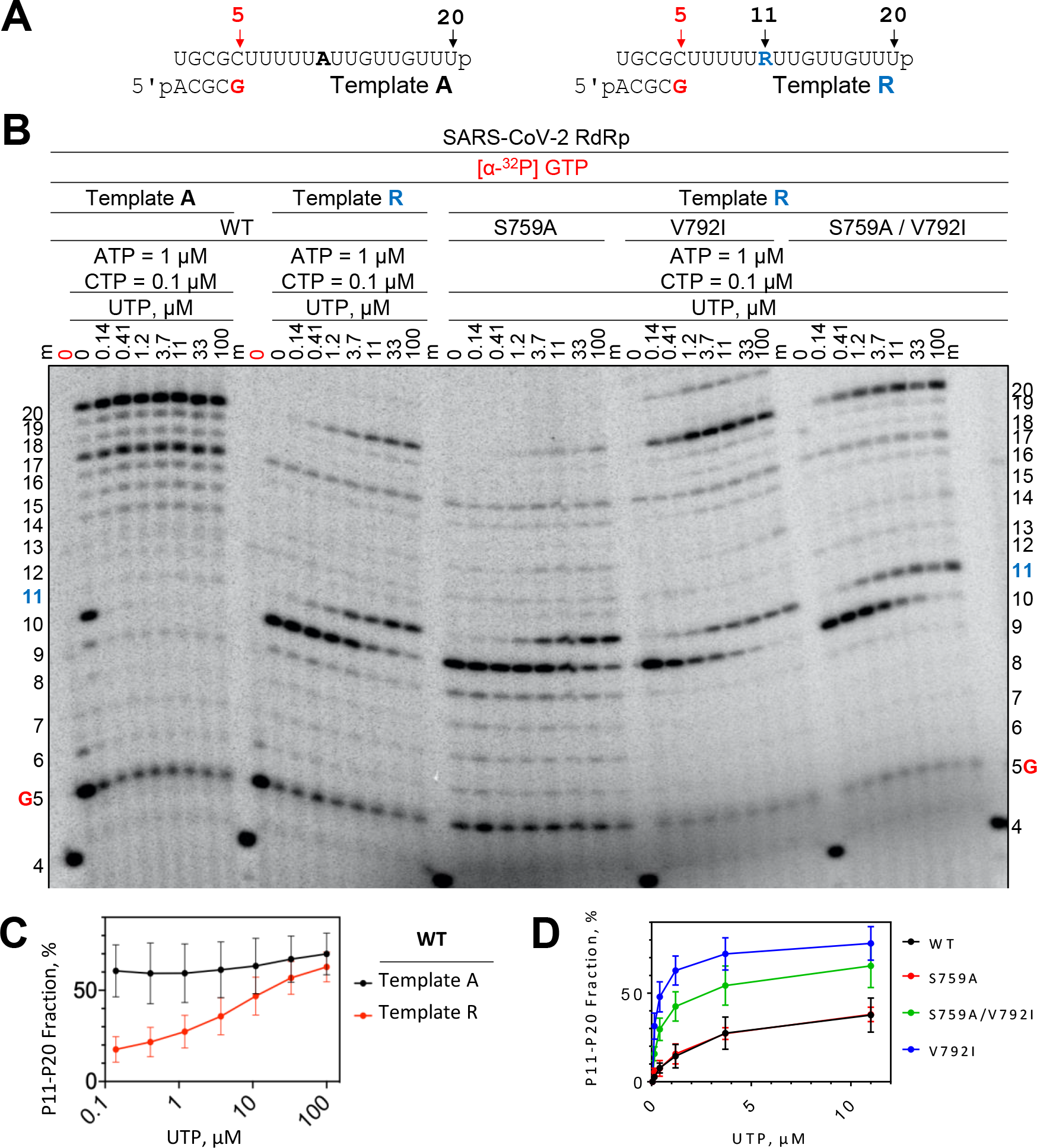
RNA synthesis by SARS-CoV-2 WT and mutant S759A, V792I, and S759A/V792I mutant RdRp complexes. **(A)** RNA primer/template sequences used are shown. **(B)** RDV-MP is embedded at position 11 in the template R strand while AMP is in the same position on the template A strand. RNA products were synthesized by the WT or mutant SARS-CoV-2 RdRps in a reaction mixture containing the primer/template pair, MgCl_2_, and indicated NTP concentrations. G (red) indicates the incorporation of [α-^32^P] GTP at position 5 and 4 indicates the migration pattern of 5’-^32^P-labeled 4-nt primer is used as a size marker. The 0 point in red indicates a reaction where [α-^32^P] GTP was the only NTP present to control for contaminating NTPs in the template preparations. **(C,D)** Graphical representations of the fraction of RNA synthesis beyond position 11 with respect to total RNA products formed. **(C)** Comparison of reactions using template A and template R with WT RdRp and increasing concentrations of UTP. **(D)** Comparisons of WT and mutant enzymes. Data corresponding to UTP = 33 and 100 µM were excluded to focus on the differences in the lower concentration range.

## Discussion

We here show that SARS-CoV-2 is capable of evolving reduced susceptibility to GS- 441524/RDV via substitutions in the nsp12 RdRp at, or in close proximity to, the RdRP S_759_DD active motif. Distinct sets of novel mutations within the RdRp arose in three separate lineages with differing degrees of population resistance, with lineage 3 co-evolving S759A and V792I substitutions that together in plaque isolates demonstrated the greatest extent of RDV resistance. Introduction of the substitutions at homologous positions in the structurally conserved MHV nsp12-RdRp (S755A and V788I) confirmed the resistance phenotype and its transferability across divergent CoVs. Biochemical kinetic studies of the mutations in the expressed RdRp complex consisting of nsp7, 8 and 12 demonstrated that S759A improved nsp12 discrimination against incorporation of RDV-TP, while V792I reduced template-dependent inhibition of RNA synthesis mediated by incorporated RMP, thereby complementing the effects of S759A. These results provide a mechanistic explanation for the co-selection and emergence of these mutations.

Although RdRp mutations previously have been reported in MHV(*5*) and SARS-CoV- 2(*30, 31*) associated with RDV resistance, this is the first report of an amino acid substitution in a CoV nsp12-RdRp S_759_DD catalytic motif, and which mediates the largest magnitude of RDV resistance observed to date. Notably, mutations are well known at the structurally equivalent HIV- 1 reverse transcriptase (RT) YM_184_DD active motif that confer resistance to nucleoside analogs(*32*). The HIV-1 RT YM_184_DD motif is relatively conserved among reverse transcriptase enzymes and M184V or M184I within this region has a significant effect on RT catalytic activity and may confer high-level resistance (>100-fold) to lamivudine (Epivir) and emtricitabine (Emtriva)(*31, 33, 34*). In our study, replacing the conserved S759 in the S_759_DD motif of the RdRp with an alanine resulted in decreased sensitivity to RDV, yet exerting diminished impact compared to similar changes in the HIV-1 RT. Nevertheless, these data pinpoint S759A—the product of a single nucleotide change and tolerated amino acid substitution in the RdRp—as a likely key determinant of RDV resistance. Finally, it remains to be determined if our *in vitro* selected S759A, V792I, or other mutations emerge *in vivo*under selection. Here, the HIV-1 example is potentially informative as the RT M184V/I substitutions first identified *in vitro* have repeatedly been confirmed to be selected *in vivo*(*35*).

The results of this study, along with our published biochemical and genetic studies, suggest that there are multiple potential genetic pathways to SARS-CoV-2 RDV resistance. These pathways may evolve both common and unique determinants within and across divergent CoVs. There remain many questions to pursue in understanding the relationship of RDV with the uniquely complex CoV multi-protein replicase, and the likely equally complex pathways to resistance *in vitro* and *in vivo*. Our previously reported MHV nsp12-RdRp RDV resistance substitutions F476L and V553L(*5*) were not detected at SARS-CoV-2 homologous F480 and V557 residues in any of the three GS-441524-passaged lineages in this study. However, our previous biochemical studies demonstrated that the SARS-CoV-2 V557L change did reduce template-dependent inhibition of RNA synthesis(*12*). In contrast, while the SARS-CoV-2 RDV resistance mutations S759A and V792I were not identified during MHV passage, introduction of the homologous substitutions in recombinant MHV mutants yielded clear reduced susceptibility RDV alone and combined. Results from another *in vitro* SARS-CoV-2 passage study with RDV linked nsp12-E802D, a residue change not observed in our studies, to low-level RDV resistance(*36*). GS-441524 forms the identical active metabolite as RDV in cells and acts through the same mechanism of RdRp inhibition but may have different intracellular pharmacokinetics and triphosphate levels(*7, 37*), which could theoretically contribute to differential outcomes observed in the *in vitro* resistance selection experiments performed with RDV vs. GS-441524.

Further, it will be critical to test mutations that are selected in other proteins of the CoV replicase complex (Data file S1), specifically in the nsp14 exonuclease, a key determinant of CoV high fidelity replication (proofreading), and native resistance nucleoside analogs such as Ribavirin and 5-Fluorouracil, as well as low-level native resistance to RDV in the MHV model(*5, 38, 39*).

Finally, it will be important to determine if the different individual and combined nsp12 mutations herein identified confer different extents of RDV resistance and fitness cost in SARS-CoV-2 compared to the SARS-CoV-2 lineages or the recombinant isogenic MHV background. These direct genetic studies of SARS-CoV-2 were in process when RDV received FDA Emergency Use Authorization (EUA) and FDA Approval for treatment of COVID-19 in October 2020. The FDA approval while stating the importance of and requiring data on SARS-CoV-2 RDV resistance determinants and potential, paradoxically triggered a halt to any newly initiated genetic studies of RDV resistance in SARS-CoV-2 using NIH or other US government support(*40–42*). This necessitated our targeting of reverse genetic studies using the non-human *betacoronavirus* MHV.

Our results also emphasize the need for additional research to determine the potential for *in vivo* resistance emergence and impact of resistance in various clinical settings and patient populations. A case report of an immunocompromised COVID-19 patient who responded poorly to RDV described a single mutation in nsp12-RdRp, but neither causal effect nor mechanism was demonstrated(*21*). Our results would predict that the barriers to RDV resistance emergence are significant but not insurmountable. The passage-selected SARS-CoV-2 RDV-resistant lineages and the targeted engineered MHV mutants displayed either impaired replication or no advantage compared to the parallel vehicle-passaged or recombinant WT controls, suggesting that development of RDV resistance in SARS-CoV-2 may confer a significant fitness cost, and consistent with our previous findings on MHV RDV-resistance mutants(*5*). Further, since RDV is administered intravenously, emergence of clinical resistance during treatment of an individual likely would be disfavored by the highly controlled duration of administration and rapid and profound reduction in virus titer. Taken together, these factors would predict substantial barriers to RDV resistance in natural variants and treated patients. Our analysis of the >6 million consensus sequences deposited to the GSAID database also demonstrate very low prevalence in global SARS-CoV-2 isolates, including Delta and Omicron variants, of the nsp12 substitutions identified in this study(*43*) (Table S2). While it is encouraging that natural variants to date have not propagated confirmed RDV resistance mutations at consensus levels, these substitutions might arise as minority variants. The possibility of RDV extended use in chronically infected or immunosuppressed patients also may increase the opportunities for SARS-CoV-2 to overcome genetic barriers and adapt for increased fitness. Our results create a reference for surveillance for RDV resistance, and support the need to pursue combination therapies targeting the RdRp through different mechanisms(*38, 44*), as well as inhibiting other replicase functions such as protease activities(*45*).

## Materials and Methods

### Data and Code Availability

The bioinformatic pipeline utilized for all RNA-seq datasets is available at https://github.com/DenisonLabVU/CoVariant.git. The Nanopore data analysis pipeline is available in the https://github.com/DenisonLabVU/MutALink.git repository. All sequencing datasets are publicly available at the NCBI Sequence Read Archive (SRA) under BioProject PRJNA787945 (RNA-seq) and PRJNA787608 (Nanopore).

### Cells and viruses

Vero E6 cells were obtained from USAMRIID and cultured in Dulbecco’s modified Eagle medium (DMEM) (Gibco) supplemented with 10% fetal bovine serum (FBS) (Gibco), 100 U/ml penicillin (Gibco), 100 mg/ml streptomycin (Gibco), and 0.25 μM amphotericin B (Corning). A549 cells overexpressing the human ACE2 receptor (A549-hACE2) (*46*) were cultured in DMEM supplemented with 10% FBS, 100 U/ml penicillin, 100 mg/ml streptomycin, and 1% MEM Non-Essential Amino Acids Solution (Gibco). Murine astrocytoma delayed brain tumor (DBT) cells and baby hamster kidney 21 cells expressing the MHV receptor (BHK-R)(*47*) were maintained in DMEM containing 10% FBS (Invitrogen), 100 U/ml penicillin, 100 mg/ml streptomycin, HEPES (Gibco), and 0.25 μM amphotericin B. BHK-R cells were further supplemented with 0.8 mg/ml G418 (Mediatech). A P3 stock of the SARS-CoV- 2/human/USA/WA-CDC-WA1/2020 isolate (GenBank accession no. MN985325.1) was obtained from the CDC and passed twice in Vero E6 cells to generate a high-titer P5 stock for experiments described in this manuscript. All work with MHV was performed using the recombinant WT strain MHV-A59 (GenBank accession no. AY910861)(*47*). Mutant MHV viruses were generated using QuikChange mutagenesis performed according to the manufacturer’s protocol to generate mutations in MHV individual genome cDNA fragment plasmids using the previously described infectious clone reverse-genetics system(*47*). Mutants were recovered in BHK-R cells following electroporation of *in vitro*-transcribed genomic RNA. All fragments containing mutations were Sanger sequenced to ensure mutations were present before use in further studies (GeneWiz, South Plainfield, NJ). RDV and GS-441524 were synthesized by the Department of Medicinal Chemistry, Gilead Sciences (Foster City, CA).

### Selection of RDV resistance

Infection was initiated in 6-well tissue-culture plates (Corning) at an MOI of 0.01 PFU SARS-CoV-2 per cell in sextuplicate. Three wells of Vero E6 cells were treated with 0.5 μM GS-441524, and three other wells were treated with 0.1% DMSO (vehicle controls), each well representing one lineage. Once cell monolayers demonstrated at least 40% CPE or after 72 h, cell culture supernatant was harvested and a constant volume of 20 μL supernatant was added to a fresh monolayer to initiate the subsequent passage. All lineages were maintained until passage 13 (P13). At P13, GS-441524 lineage 2 was reduced to 3 μM to allow for virus recovery, whereas lineage 1 and 3 replicated in 9 μM GS-441524. P13 virus lineages were titered by plaque assay, and levels of resistance were determined in an antiviral activity assay. In addition, RNA was harvested from infected cell supernatant using TRIzol LS reagent (Invitrogen) and cell monolayers using TRIzol reagent (Invitrogen) for viral population sequencing. Passages from GS-441524-treated lineages were subjected to viral plaque isolation by standard plaque assay in the absence of GS-441524. Plaque picks (PP) were expanded in Vero E6 cultures supplemented with 1 μM GS-441524. Cultures were harvested when CPE was >50% or after 72 h. Supernatant RNA was collected for Sanger sequencing and total monolayer RNA was harvested for RNA-seq.

### Antiviral activity assays

A549-hACE2 cells were seeded at 5 x 10_4_ cells per well in 48-well plates (Corning) and allowed to adhere for 16-24 h. RDV (20 μM in DMSO stock) was serially diluted in DMSO to achieve 1000x final concentration and diluted to final 1x concentration in culture medium up to 2 h before start of infection. Cells were adsorbed with MOI = 0.01 PFU/cell of passaged virus lineages in gel saline for 30 min at 37°C and gently rocked manually every 10 minutes to redistribute the inoculum. Viral inoculum was removed, and cells were washed with pre-warmed PBS containing CaCl_2_ and MgCl_2_ (PBS +/+) (Corning). Medium containing dilutions of RDV or vehicle control (0.1% DMSO) was added and following incubation at 37°C/5% CO_2_ for 48 h, cell culture supernatants were harvested and processed for viral genomic RNA quantification by RT-qPCR. Data represent the means of two independent experiments consisting of 2 replicates each.

### Viral replication assays

A549-hACE2 or DBT-9 cells were seeded at 1 x 10_5_ cells per well in 24-well plates (Corning) and allowed to reach confluence within 24 h. A549-hACE2 cells were adsorbed with MOI = 0.01 PFU/ml SARS-CoV-2 passaged population virus or plaque-isolated sub-lineages. DBT-9 cells were adsorbed with MOI = 0.01 PFU/ml WT MHV (derived from the infectious clone) or with MHV recombinantly engineered to contain putative RDV-resistance mutations in the isogenic background. Cells were adsorbed with virus for 30 min at 37°C/5% CO_2_, with manual rocking every 10 min to redistribute the viral inoculum, after which the inoculum was removed, cells were washed with pre-warmed PBS +/+, and fresh medium without drug was added. Cultures were incubated at 37°C/5% CO_2_, supernatants were harvested at indicated times post infection, and MHV infectious titers were determined via plaque assay as previously described(*48*). Viral genomic RNA in culture supernatants was quantified by RT-qPCR.

### Quantification of SARS-CoV-2 infectious titer

Approximately 1 x 10_6_ Vero E6 cells/well were seeded in 6-well plates and allowed to reach confluence within 24 h. Medium was removed, and 100 µL of 10-fold serial dilutions of virus-containing supernatants in gelatin saline (0.3% [wt/vol] gelatin in PBS +/+) was adsorbed in duplicate wells for 30 min at 37°C/5% CO_2_. Plates were rocked manually every 10 minutes to redistribute inoculum. Cells were overlaid with DMEM containing 1% agar and incubated at 37°C/5% CO_2_. Plaques were enumerated in unstained monolayers at 48-72 h post infection.

### Quantification of viral RNA

Cell culture supernatants were harvested in TRIzol LS reagent, and RNA was purified following phase separation by chloroform as recommended by the manufacturer. RNA in the aqueous phase was collected and further purified using a KingFisher II automated nucleic acid extraction system (ThermoFisher Scientific) according to manufacturer’s protocol. Viral RNA was quantified by RT-qPCR on a StepOnePlus Real-Time PCR System (Applied Biosystems) using TaqMan Fast Virus 1-Step Master Mix chemistry (Applied Biosystems). SARS-CoV-2 genomic RNA was amplified and detected using forward (5’- CGTGTAGTCTTTAATGGTGTTTCC-3’) and reverse (5’-GCACATCACTACGCAACTTTAG-3’) primers and probe (5’-FAM-TTTGAAGAAGCTGCGCTGTGCAC-BHQ-1-3’) specific for the nsp4 gene. RNA copy numbers were interpolated from a standard curve produced with serial 10-fold dilutions of nsp4 gene RNA. Briefly, SARS-CoV-2 cloned nsp4 gene cDNA served as template to PCR-amplify a 1062 bp product using forward (5’- TAATACGACTCACTATAGGCTGCTGAATGTACAATTTT-3’) and reverse (5’-CTGCAAAACAGCTGAGGTGATAGAG-3’) primers that appended a T7 RNA polymerase promoter to the 5’ end. PCR product was column purified (Promega) for subsequent *in vitro* transcription of nsp4 RNA using mMESSAGE mMACHINE T7 Transcription Kit (ThermoFisher Scientific) according to manufacturer’s protocol. Nsp4 RNA was purified using the RNeasy mini kit (Qiagen) according to manufacturer’s protocol, and copy number was calculated using SciencePrimer.com copy number calculator. RNA copy numbers from MHV infections were quantified as previously described(*49*).

### Illumina sequencing

Total RNA was extracted from P9 and P13 monolayers using TRIzol according to the manufacturer’s instructions. For RNA-Seq, total RNA underwent poly(A) selection followed by NovaSeq PE150 sequencing (Illumina) at 15 million reads per sample at the VUMC core facility, VANTAGE. Reads were aligned to the reference genome (MT020881.1), and mutations were identified, quantified, and annotated using the in-house pipeline, *CoVariant*. Amino acid locations were confirmed through sequence alignment using MacVector and CLC Workbench (QIAGEN).

### Nanopore amplicon sequencing

5 μL of RNA from infected cell monolayers was reverse transcribed using random hexamers and Superscript III (ThermoFisher) to generate the first cDNA strand for each sample according to manufacturer’s protocols. Nsp12 amplicons 2796 bp in size were generated with first-round EasyA (Agilent) PCR using tailed primers according to manufacturer’s protocols (forward = 5’- TTTCTGTTGGTGCTGATATTGCCTGTAGATGCTGCTAAAGC-3’; reverse = 5’- ACTTGCCTGTCGCTCTATCTTCTGACATCACAACCTGGAGC-3’) and confirmed by gel electrophoresis. PCR products were purified by the Wizard SV Gel and PCR Clean-Up System (Promega) and quantified using the Qubit dsDNA HS assay (ThermoFisher). For each sample, 1 μg of DNA (505.7 fmol) was used for barcoding PCR according to manufacturer’s protocols for the EXP-PBC001 kit (Oxford Nanopore Technologies). Barcoded amplicons were purified by the Wizard SV Gel and PCR Clean-Up System (Promega) and quantified using the Qubit dsDNA HS assay. Amplicons were pooled using 112 ng of amplicon DNA per sample for a total of 1 μg of amplicon DNA. Sequencing of the library prep was performed according to manufacturer’s protocols using the SQK-LSK110 kit (Oxford Nanopore Technologies). The pooled library was loaded onto a quality-checked MinION flowcell with 1491 functional sequencing pores, and sequencing was performed using the *MinKNOW* GUI over 72 hours.

### Nanopore genetic linkage analysis

Mutation linkage was determined using a newly developed, in-house pipeline, *MutALink*. Analysis was directed by sequential custom Bash shell scripts that direct each module of the pipeline. The first module performs basecalling and alignment.

Specifically, following sequencing, raw FAST5 files were basecalled and demultiplexed using *Guppy v5.0.11* (Oxford Nanopore Technologies). Pass FASTQ files were aligned to the SARS-CoV-2 genome (MT020881.1) for each sample using *minimap2*(*50*), and alignments were processed and filtered for reads containing sequences across all of nsp12 using *SAMtools*(*51*). Alignment statistics were generated using *NanoStat*. The second module of the *MutALink* pipeline calls and quantifies variant allele frequencies for candidate variants using *Nanopolish*(*52*). The last module, genotype quantification, filters different combinations of candidate variants and generates outputs for each lineage using a custom batch script, variant-specific javascript files (V166A.js, N198S.js, V792I.js, S759A.js, and C799F.js), and the *samjdk* package from *jvarkit* (*53*) using a separate Bash shell script for each lineage. Read counts were corrected manually for duplicate counting between combinations, and the frequency of each genotype in each sample passage compared to total mapped reads was reported and visualized using the Python package, *seaborn*(*54*).

### Structural modeling

The model of the SARS-CoV-2 pre-incorporation polymerase complex was built on the cryo-EM PDB structure 6XEZ(*24*) by examination of several post-incorporation/pre-translocation SARS-CoV-2 RdRp structures compared to a number of pre-incorporation complexes of similar viral RdRps (i.e., HCV, norovirus and poliovirus)(*25–27*). First ATP was positioned in the active site, as were two Mg_++_ ions. The corresponding template base at position +1 was modified from A to U. D618, D760 and D761 were optimized to coordinate the metal ions, and a conformational search was done on key sidechains in the active site, including K545, R553, R555, D623, S682, T687, N691, D759 and K798(*55*). These residues, as well as metals, ATP, and primer/template nucleotides P_-1_, T_-1_ and T_+1_ were then minimized(*56*). Once optimized, ATP was modified to RDV-TP and the structure was minimized again. Mutations were analyzed by conducting a conformational search of all residues within 5 Å of the mutation and minimizing. In the case of V166A, V792I and C799R/F, a conformational search of the loops 163-168 and 790-800 was also conducted.

### Protein expression and purification

SARS–CoV-2 RdRp WT and mutant proteins (S759A and V792I) were expressed and purified as reported previously(*8, 9, 12*). The pFastBac-1 (Invitrogen, Burlington, Ontario, Canada) plasmid with codon-optimized synthetic DNA sequences (GenScript, Piscataway, NJ) coding for a portion of 1ab polyproteins of SARS-CoV-2 (NCBI: QHD43415.1), containing only nsp5, nsp7, nsp8, and nsp12, was used as starting material for protein expression in insect cells (Sf9, Invitrogen). We employed the MultiBac (Geneva Biotech, Indianapolis, IN) system for protein expression in insect cells according to published protocols(*57, 58*).

### Single NTP incorporation and the effect of primer-embedded RDV-MP

NTP incorporation by SARS-CoV-2 RdRp WT and mutants, data acquisition and quantification were done as reported(*8, 9, 12*). Enzyme concentration was 150 nM for both single and multiple nucleotide incorporation assays, respectively. RNA synthesis incubation time was 10 min. Single nucleotide incorporation assays were used to determine the preference for the natural nucleotide over RDV-TP. The selectivity value was calculated as a ratio of the incorporation efficiencies of the natural nucleotide over the nucleotide analog. The discrimination value was calculated as a ratio of mutant to WT selectivity. The efficiency of nucleotide incorporation was determined by the ratio of Michaelis–Menten constants *V*_max_ over *K*_m_ as previously reported(*8, 9, 12*).

### Evaluation of RNA synthesis across the RNA template with embedded RDV-MP

RNA synthesis assays using SARS-CoV-2 RdRp complex on an RNA template with an embedded RDV-MP or adenosine at equivalent positions, data acquisition and quantification were done as previously described with the following adjustments: (1) enzyme concentration of WT RdRp was increased to 250 nM and (2) mutant RdRp concentration was adjusted such that activity was equivalent to WT. Two independent preparations of RDV-embedded RNA templates and at least three independent preparations of SARS-CoV-2 WT and mutant enzymes were used.

### Mathematical and statistical analyses

The EC_50_ value was calculated in GraphPad Prism 8 as the concentration at which there was a 50% decrease in viral replication relative to vehicle alone (0% inhibition). Dose-response curves were fit based using four-parameter non-linear regression. All statistical tests were executed using GraphPad Prism 8. Statistical details of experiments are described in the figure legends.

## Supplementary Materials

Fig. S1. Serial passaging of SARS-CoV-2.

Fig. S2. Selective incorporation of RDV-TP by WT and mutant S759A, and V792I SARS-CoV-2 RdRp complexes.

Fig. S3. Competition between RDV-TP and natural NTPs in SARS-CoV-2 WT and mutant S759A, V792I, and S759A/V792I RdRp complexes.

Table S1. EC50 and fold change for GS-441524 and vehicle-passaged virus lineages.

Table S2. Number of times the nsp12 amino acid substitutions were detected in SARS-CoV-2 sequences deposited to GISAID database

Data file S1. Mutations present at >1% frequency in populations of serially passaged SARS-CoV- 2.

## Acknowledgements

We thank Dr. Natalie Thornburg at the Centers for Disease Control and Prevention in Atlanta, USA for providing the WA-1 SARS-CoV-2 used in this study. Finally, we thank VUMC and UNC Environmental Health and Safety personnel and institutional biosafety committees for reviewing and approving the work described herein. All RDV passage and forward genetic selection studies with SARS-CoV-2 were completed before the October 2020 FDA EUA and approval for remdesivir.

## Author Contributions

LJS, AJP, HWL, CJG, and EPT conceived, designed, and performed experiments and managed and coordinated responsibility for research activity, planning, and execution. LJS, AJP, MG, and JKP, wrote the manuscript. LJS, AJP, HWL, CG, EPT, JG, ASG, TMH, XL, JL, and JKP performed experiments. TPS, RSB, MG, and MRD directed the funded programs and oversaw experimental design and interpretation. MG, TPS, RSB, MRD, DKP, and TC edited the manuscript.

## Data and Materials Availability

All data are available in the main text or the supplementary materials.

## References

1. CDC, COVID Data Tracker. Cent. Dis. Control Prev. (2020), (available at https://covid.cdc.gov/covid-data-tracker).

2. WHO Coronavirus (COVID-19) Dashboard, (available at https://covid19.who.int).

3. T. K. Warren, R. Jordan, M. K. Lo, A. S. Ray, R. L. Mackman, V. Soloveva, D. Siegel, M. Perron, R. Bannister, H. C. Hui, N. Larson, R. Strickley, J. Wells, K. S. Stuthman, S. A. V. Tongeren, N. L. Garza, G. Donnelly, A. C. Shurtleff, C. J. Retterer, D. Gharaibeh, R. Zamani, T. Kenny, B. P. Eaton, E. Grimes, L. S. Welch, L. Gomba, C. L. Wilhelmsen, D. K. Nichols, J. E. Nuss, E. R. Nagle, J. R. Kugelman, G. Palacios, E. Doerffler, S. Neville, E. Carra, M. O. Clarke, L. Zhang, W. Lew, B. Ross, Q. Wang, K. Chun, L. Wolfe, D. Babusis, Y. Park, K. M. Stray, I. Trancheva, J. Y. Feng, O. Barauskas, Y. Xu, P. Wong, M. R. Braun, M. Flint, L. K. McMullan, S.-S. Chen, R. Fearns, S. Swaminathan, D. L. Mayers, C. F. Spiropoulou, W. A. Lee, S. T. Nichol, T. Cihlar, S. Bavari, Therapeutic efficacy of the small molecule GS-5734 against Ebola virus in rhesus monkeys. Nature. 531, 381–385 (2016).

4. T. P. Sheahan, A. C. Sims, R. L. Graham, V. D. Menachery, L. E. Gralinski, J. B. Case, S. R. Leist, K. Pyrc, J. Y. Feng, I. Trantcheva, R. Bannister, Y. Park, D. Babusis, M. O. Clarke, R. L. Mackman, J. E. Spahn, C. A. Palmiotti, D. Siegel, A. S. Ray, T. Cihlar, R. Jordan, M. R. Denison, R. S. Baric, Broad-spectrum antiviral GS-5734 inhibits both epidemic and zoonotic coronaviruses. Sci. Transl. Med. 9, eaal3653 (2017).

5. M. L. Agostini, E. L. Andres, A. C. Sims, R. L. Graham, T. P. Sheahan, X. Lu, E. C. Smith, J. B. Case, J. Y. Feng, R. Jordan, A. S. Ray, T. Cihlar, D. Siegel, R. L. Mackman, M. O. Clarke, R. S. Baric, M. R. Denison, Coronavirus Susceptibility to the Antiviral Remdesivir (GS-5734) Is Mediated by the Viral Polymerase and the Proofreading Exoribonuclease. mBio. 9, e00221–18 (2018).

6. A. J. Brown, J. J. Won, R. L. Graham, K. H. Dinnon, A. C. Sims, J. Y. Feng, T. Cihlar, M. R. Denison, R. S. Baric, T. P. Sheahan, Broad spectrum antiviral remdesivir inhibits human endemic and zoonotic deltacoronaviruses with a highly divergent RNA dependent RNA polymerase. Antiviral Res. 169, 104541 (2019).

7. A. J. Pruijssers, A. S. George, A. Schäfer, S. R. Leist, L. E. Gralinksi, K. H. Dinnon, B. L. Yount, M. L. Agostini, L. J. Stevens, J. D. Chappell, X. Lu, T. M. Hughes, K. Gully, D. R. Martinez, A. J. Brown, R. L. Graham, J. K. Perry, V. Du Pont, J. Pitts, B. Ma, D. Babusis, E. Murakami, J. Y. Feng, J. P. Bilello, D. P. Porter, T. Cihlar, R. S. Baric, M. R. Denison, T. P. Sheahan, Remdesivir Inhibits SARS-CoV-2 in Human Lung Cells and Chimeric SARS-CoV Expressing the SARS-CoV-2 RNA Polymerase in Mice. Cell Rep. 32, 107940 (2020).

8. C. J. Gordon, E. P. Tchesnokov, E. Woolner, J. K. Perry, J. Y. Feng, D. P. Porter, M. Gotte, Remdesivir is a direct-acting antiviral that inhibits RNA-dependent RNA polymerase from severe acute respiratory syndrome coronavirus 2 with high potency. J. Biol. Chem. (2020), doi:10.1074/jbc.RA120.013679.

9. C. J. Gordon, E. P. Tchesnokov, J. Y. Feng, D. P. Porter, M. Gotte, The antiviral compound remdesivir potently inhibits RNA-dependent RNA polymerase from Middle East respiratory syndrome coronavirus. J. Biol. Chem. (2020), doi:10.1074/jbc.AC120.013056.

10. J. P. K. Bravo, T. L. Dangerfield, D. W. Taylor, K. A. Johnson, Remdesivir is a delayed translocation inhibitor of SARS-CoV-2 replication. Mol. Cell. 81, 1548–1552.e4 (2021).

11. G. Kokic, H. S. Hillen, D. Tegunov, C. Dienemann, F. Seitz, J. Schmitzova, L. Farnung, A. Siewert, C. Höbartner, P. Cramer, Mechanism of SARS-CoV-2 polymerase stalling by remdesivir. Nat. Commun. 12, 279 (2021).

12. E. P. Tchesnokov, C. J. Gordon, E. Woolner, D. Kocinkova, J. K. Perry, J. Y. Feng, D. P. Porter, M. Götte, Template-dependent inhibition of coronavirus RNA-dependent RNA polymerase by remdesivir reveals a second mechanism of action. J. Biol. Chem. 295, 16156– 16165 (2020).

13. M. Seifert, S. C. Bera, P. van Nies, R. N. Kirchdoerfer, A. Shannon, T.-T.-N. Le, X. Meng, H. Xia, J. M. Wood, L. D. Harris, F. S. Papini, J. J. Arnold, S. Almo, T. L. Grove, P.-Y. Shi, Y. Xiang, B. Canard, M. Depken, C. E. Cameron, D. Dulin, Inhibition of SARS-CoV-2 polymerase by nucleotide analogs from a single-molecule perspective. eLife. 10, e70968 (2021).

14. E. de Wit, F. Feldmann, J. Cronin, R. Jordan, A. Okumura, T. Thomas, D. Scott, T. Cihlar, H. Feldmann, Prophylactic and therapeutic remdesivir (GS-5734) treatment in the rhesus macaque model of MERS-CoV infection. Proc. Natl. Acad. Sci. 117, 6771–6776 (2020).

15. A. T. P. Sheahan, A. C. Sims, S. R. Leist, A. Schäfer, J. Won, A. J. Brown, S. A. Montgomery, A. Hogg, D. Babusis, M. O. Clarke, J. E. Spahn, L. Bauer, S. Sellers, D. Porter, J. Y. Feng, T. Cihlar, R. Jordan, M. R. Denison, R. S. Baric, Comparative therapeutic efficacy of remdesivir and combination lopinavir, ritonavir, and interferon beta against MERS-CoV. Nat. Commun. 11, 222 (2020).

16. D. R. Martinez, A. Schäfer, S. R. Leist, D. Li, K. Gully, B. Yount, J. Y. Feng, E. Bunyan, D. P. Porter, T. Cihlar, S. A. Montgomery, B. F. Haynes, R. S. Baric, M. C. Nussenzweig, T. P. Sheahan, Prevention and therapy of SARS-CoV-2 and the B.1.351 variant in mice. Cell Rep. 36, 109450 (2021).

17. J. H. Beigel, K. M. Tomashek, L. E. Dodd, A. K. Mehta, B. S. Zingman, A. C. Kalil, E. Hohmann, H. Y. Chu, A. Luetkemeyer, S. Kline, D. Lopez de Castilla, R. W. Finberg, K. Dierberg, V. Tapson, L. Hsieh, T. F. Patterson, R. Paredes, D. A. Sweeney, W. R. Short, G. Touloumi, D. C. Lye, N. Ohmagari, M.-D. Oh, G. M. Ruiz-Palacios, T. Benfield, G. Fätkenheuer, M. G. Kortepeter, R. L. Atmar, C. B. Creech, J. Lundgren, A. G. Babiker, S. Pett, J. D. Neaton, T. H. Burgess, T. Bonnett, M. Green, M. Makowski, A. Osinusi, S. Nayak, H. C. Lane, ACTT-1 Study Group Members, Remdesivir for the Treatment of Covid-19 - Preliminary Report. N. Engl. J. Med. (2020), doi:10.1056/NEJMoa2007764.

18. C. D. Spinner, R. L. Gottlieb, G. J. Criner, J. R. A. López, A. M. Cattelan, A. S. Viladomiu, O. Ogbuagu, P. Malhotra, K. M. Mullane, A. Castagna, L. Y. A. Chai, M. Roestenberg, O. T. Y. Tsang, E. Bernasconi, P. L. Turnier, S.-C. Chang, D. SenGupta, R. H. Hyland, A. O. Osinusi, H. Cao, C. Blair, H. Wang, A. Gaggar, D. M. Brainard, M. J. McPhail, S. Bhagani, M. Y. Ahn, A. J. Sanyal, G. Huhn, F. M. Marty, for the G.-U.-540-5774 Investigators, Effect of Remdesivir vs Standard Care on Clinical Status at 11 Days in Patients With Moderate COVID-19: A Randomized Clinical Trial. JAMA. 324, 1048–1057 (2020).

19. Gilead’s Veklury® (Remdesivir) Associated With a Reduction in Mortality Rate in Hospitalized Patients With COVID-19 Across Three Analyses of Large Retrospective Real-World Data Sets, (available at https://www.gilead.com/news-and-press/press-room/press-releases/2021/6/gileads-veklury-remdesivir-associated-with-a-reduction-in-mortality-rate-in-hospitalized-patients-with-covid19-across-three-analyses-of-large-ret).

20. R. L. Gottlieb, C. E. Vaca, R. Paredes, J. Mera, B. J. Webb, G. Perez, G. Oguchi, P. Ryan, B. U. Nielsen, M. Brown, A. Hidalgo, Y. Sachdeva, S. Mittal, O. Osiyemi, J. Skarbinski, K. Juneja, R. H. Hyland, A. Osinusi, S. Chen, G. Camus, M. Abdelghany, S. Davies, N. Behenna-Renton, F. Duff, F. M. Marty, M. J. Katz, A. A. Ginde, S. M. Brown, J. T. Schiffer, J. A. Hill, GS-US-540-9012 (PINETREE) Investigators, Early Remdesivir to Prevent Progression to Severe Covid-19 in Outpatients. N. Engl. J. Med. (2021), doi:10.1056/NEJMoa2116846.

21. M. Martinot, A. Jary, S. Fafi-Kremer, V. Leducq, H. Delagreverie, M. Garnier, J. Pacanowski, A. Mékinian, F. Pirenne, P. Tiberghien, V. Calvez, C. Humbrecht, A.-G. Marcelin, K. Lacombe, Emerging RNA-Dependent RNA Polymerase Mutation in a Remdesivir-Treated B-cell Immunodeficient Patient With Protracted Coronavirus Disease 2019. Clin. Infect. Dis. (2020), doi:10.1093/cid/ciaa1474.

22. J. Harcourt, A. Tamin, X. Lu, S. Kamili, S. K. Sakthivel, J. Murray, K. Queen, Y. Tao, C. R. Paden, J. Zhang, Y. Li, A. Uehara, H. Wang, C. Goldsmith, H. A. Bullock, L. Wang, B. Whitaker, B. Lynch, R. Gautam, C. Schindewolf, K. G. Lokugamage, D. Scharton, J. A. Plante, D. Mirchandani, S. G. Widen, K. Narayanan, S. Makino, T. G. Ksiazek, K. S. Plante, S. C. Weaver, S. Lindstrom, S. Tong, V. D. Menachery, N. J. Thornburg, Early Release - Severe Acute Respiratory Syndrome Coronavirus 2 from Patient with 2019 Novel Coronavirus Disease, United States - Volume 26, Number 6—June 2020 - Emerging Infectious Diseases journal - CDC, doi:10.3201/eid2606.200516.

23. GISAID - Initiative, (available at https://www.gisaid.org/).

24. J. Chen, B. Malone, E. Llewellyn, M. Grasso, P. M. M. Shelton, P. D. B. Olinares, K. Maruthi, E. T. Eng, H. Vatandaslar, B. T. Chait, T. M. Kapoor, S. A. Darst, E. A. Campbell, Structural Basis for Helicase-Polymerase Coupling in the SARS-CoV-2 Replication- Transcription Complex. Cell. 182, 1560–1573.e13 (2020).

25. T. C. Appleby, J. K. Perry, E. Murakami, O. Barauskas, J. Feng, A. Cho, D. Fox, D. R. Wetmore, M. E. McGrath, A. S. Ray, M. J. Sofia, S. Swaminathan, T. E. Edwards, Viral replication. Structural basis for RNA replication by the hepatitis C virus polymerase. Science. 347, 771–775 (2015).

26. D. F. Zamyatkin, F. Parra, J. M. M. Alonso, D. A. Harki, B. R. Peterson, P. Grochulski, K. K.-S. Ng, Structural Insights into Mechanisms of Catalysis and Inhibition in Norwalk Virus Polymerase*. J. Biol. Chem. 283, 7705–7712 (2008).

27. P. Gong, O. B. Peersen, Structural basis for active site closure by the poliovirus RNA- dependent RNA polymerase. Proc. Natl. Acad. Sci. U. S. A. 107, 22505–22510 (2010).

28. X. Yang, E. D. Smidansky, K. R. Maksimchuk, D. Lum, J. L. Welch, J. J. Arnold, C. E. Cameron, D. D. Boehr, Motif D of viral RNA-dependent RNA polymerases determines efficiency and fidelity of nucleotide addition. Struct. Lond. Engl. 1993. 20, 1519–1527 (2012).

29. T. L. Dangerfield, N. Z. Huang, K. A. Johnson, Remdesivir Is Effective in Combating COVID-19 because It Is a Better Substrate than ATP for the Viral RNA-Dependent RNA Polymerase. iScience. 23, 101849 (2020).

30. M. Pachetti, B. Marini, F. Benedetti, F. Giudici, E. Mauro, P. Storici, C. Masciovecchio, S. Angeletti, M. Ciccozzi, R. C. Gallo, D. Zella, R. Ippodrino, Emerging SARS-CoV-2 mutation hot spots include a novel RNA-dependent-RNA polymerase variant. J. Transl. Med. 18, 179 (2020).

31. A. M. Szemiel, A. Merits, R. J. Orton, O. A. MacLean, R. M. Pinto, A. Wickenhagen, G. Lieber, M. L. Turnbull, S. Wang, W. Furnon, N. M. Suarez, D. Mair, A. da S. Filipe, B. J. Willett, S. J. Wilson, A. H. Patel, E. C. Thomson, M. Palmarini, A. Kohl, M. E. Stewart, In vitro selection of Remdesivir resistance suggests evolutionary predictability of SARS-CoV- 2. PLOS Pathog. 17, e1009929 (2021).

32. S. Garforth, C. Lwatula, V. Prasad, The Lysine 65 Residue in HIV-1 Reverse Transcriptase Function and in Nucleoside Analog Drug Resistance. Viruses. 6, 4080–4094 (2014).

33. P. L. Boyer, H.-Q. Gao, P. K. Clark, S. G. Sarafianos, E. Arnold, S. H. Hughes, YADD Mutants of Human Immunodeficiency Virus Type 1 and Moloney Murine Leukemia Virus Reverse Transcriptase Are Resistant to Lamivudine Triphosphate (3TCTP) In Vitro. J. Virol. 75, 6321–6328 (2001).

34. K. Diallo, B. Brenner, M. Oliveira, D. Moisi, M. Detorio, M. Götte, M. A. Wainberg, The M184V Substitution in Human Immunodeficiency Virus Type 1 Reverse Transcriptase Delays the Development of Resistance to Amprenavir and Efavirenz in Subtype B and C Clinical Isolates. Antimicrob. Agents Chemother. 47, 2376–2379 (2003).

35. R. F. Schinazi, R. M. Lloyd, M. H. Nguyen, D. L. Cannon, A. McMillan, N. Ilksoy, C. K. Chu, D. C. Liotta, H. Z. Bazmi, J. W. Mellors, Characterization of human immunodeficiency viruses resistant to oxathiolane-cytosine nucleosides. Antimicrob. Agents Chemother. 37, 875–881 (1993).

36. A. M. Szemiel, A. Merits, R. J. Orton, O. MacLean, R. M. Pinto, A. Wickenhagen, G. Lieber, M. L. Turnbull, S. Wang, D. Mair, A. da S. Filipe, B. J. Willett, S. J. Wilson, A. H. Patel, E. C. Thomson, M. Palmarini, A. Kohl, M. E. Stewart, bioRxiv, in press, doi:10.1101/2021.02.01.429199.

37. R. L. Mackman, H. C. Hui, M. Perron, E. Murakami, C. Palmiotti, G. Lee, K. Stray, L. Zhang, B. Goyal, K. Chun, D. Byun, D. Siegel, S. Simonovich, V. Du Pont, J. Pitts, D. Babusis, A. Vijjapurapu, X. Lu, C. Kim, X. Zhao, J. Chan, B. Ma, D. Lye, A. Vandersteen, S. Wortman, K. T. Barrett, M. Toteva, R. Jordan, R. Subramanian, J. P. Bilello, T. Cihlar, Prodrugs of a 1’-CN-4-Aza-7,9-dideazaadenosine C-Nucleoside Leading to the Discovery of Remdesivir (GS-5734) as a Potent Inhibitor of Respiratory Syncytial Virus with Efficacy in the African Green Monkey Model of RSV. J. Med. Chem. 64, 5001–5017 (2021).

38. M. L. Agostini, A. J. Pruijssers, J. D. Chappell, J. Gribble, X. Lu, E. L. Andres, G. R. Bluemling, M. A. Lockwood, T. P. Sheahan, A. C. Sims, M. G. Natchus, M. Saindane, A. A. Kolykhalov, G. R. Painter, R. S. Baric, M. R. Denison, Small-Molecule Antiviral β-d-N4- Hydroxycytidine Inhibits a Proofreading-Intact Coronavirus with a High Genetic Barrier to Resistance. J. Virol. 93 (2019), doi:10.1128/JVI.01348-19.

39. K. Graepel, X. Lu, J. B. Case, N. R. Sexton, E. C. Smith, M. R. Denison, Proofreading- deficient coronaviruses adapt for increased fitness over long-term passage without reversion of exoribonuclease-inactivating mutations (2017), doi:10.1101/175562.

40. NIH Guidelines for Research Involving Recombinant or Synthetic Nucleic Acid Molecules (NIH Guidelines) - April 2019, 149 (2019).

41. Research Involving Enhanced Potential Pandemic Pathogens. *Natl. Inst. Health NIH* (2021), (available at https://www.nih.gov/news-events/research-involving-potential-pandemic-pathogens).

42. Framework for Guiding Funding Decisions about Proposed Research Involving Enhanced Potential Pandemic Pathogens. https://www.phe.gov/s3/dualuse/Documents/P3CO.pdf, (available at https://www.phe.gov/s3/dualuse/Documents/P3CO.pdf).

43. COVID CG, (available at https://covidcg.org/?tab=location&selectedProtein=nsp12%20-%20RdRp&residueCoordinates=1,932&coordinateMode=protein&startDate=2019-12-15&submStartDate=2019-12-15&submEndDate=2021-09-23&region=Africa&region=Asia&region=Europe&region=North%20America&region=Oceania&region=South%20America).

44. T. P. Sheahan, A. C. Sims, S. Zhou, R. L. Graham, A. J. Pruijssers, M. L. Agostini, S. R. Leist, A. Schäfer, K. H. Dinnon, L. J. Stevens, J. D. Chappell, X. Lu, T. M. Hughes, A. S. George, C. S. Hill, S. A. Montgomery, A. J. Brown, G. R. Bluemling, M. G. Natchus, M. Saindane, A. A. Kolykhalov, G. Painter, J. Harcourt, A. Tamin, N. J. Thornburg, R. Swanstrom, M. R. Denison, R. S. Baric, An orally bioavailable broad-spectrum antiviral inhibits SARS-CoV-2 in human airway epithelial cell cultures and multiple coronaviruses in mice. Sci. Transl. Med. (2020), doi:10.1126/scitranslmed.abb5883.

45. M. D. Hall, J. M. Anderson, A. Anderson, D. Baker, J. Bradner, K. R. Brimacombe, E. A. Campbell, K. S. Corbett, K. Carter, S. Cherry, L. Chiang, T. Cihlar, E. de Wit, M. Denison, M. Disney, C. V. Fletcher, S. L. Ford-Scheimer, M. Götte, A. C. Grossman, F. G. Hayden, D. J. Hazuda, C. A. Lanteri, H. Marston, A. D. Mesecar, S. Moore, J. O. Nwankwo, J. O’Rear, G. Painter, K. Singh Saikatendu, C. A. Schiffer, T. P. Sheahan, P.-Y. Shi, H. D. Smyth, M. J. Sofia, M. Weetall, S. K. Weller, R. Whitley, A. S. Fauci, C. P. Austin, F. S. Collins, A. J. Conley, M. I. Davis, Report of the National Institutes of Health SARS-CoV-2 Antiviral Therapeutics Summit. J. Infect. Dis. 224, S1–S21 (2021).

46. Y. J. Hou, S. Chiba, P. Halfmann, C. Ehre, M. Kuroda, K. H. Dinnon, S. R. Leist, A. Schäfer, N. Nakajima, K. Takahashi, R. E. Lee, T. M. Mascenik, R. Graham, C. E. Edwards, L. V. Tse, K. Okuda, A. J. Markmann, L. Bartelt, A. de Silva, D. M. Margolis, R. C. Boucher, S. H. Randell, T. Suzuki, L. E. Gralinski, Y. Kawaoka, R. S. Baric, SARS-CoV-2 D614G variant exhibits efficient replication ex vivo and transmission in vivo. Science. 370, 1464– 1468 (2020).

47. B. Yount, M. R. Denison, S. R. Weiss, R. S. Baric, Systematic Assembly of a Full-Length Infectious cDNA of Mouse Hepatitis Virus Strain A59. J. Virol. 76, 11065–11078 (2002).

48. L. D. Eckerle, X. Lu, S. M. Sperry, L. Choi, M. R. Denison, High Fidelity of Murine Hepatitis Virus Replication Is Decreased in nsp14 Exoribonuclease Mutants. J. Virol. 81, 12135– 12144 (2007).

49. E. C. Smith, H. Blanc, M. Vignuzzi, M. R. Denison, Coronaviruses Lacking Exoribonuclease Activity Are Susceptible to Lethal Mutagenesis: Evidence for Proofreading and Potential Therapeutics. PLoS Pathog. 9, e1003565 (2013).

50. H. Li, Minimap2: pairwise alignment for nucleotide sequences. Bioinformatics. 34, 3094– 3100 (2018).

51. P. Danecek, J. K. Bonfield, J. Liddle, J. Marshall, V. Ohan, M. O. Pollard, A. Whitwham, T. Keane, S. A. McCarthy, R. M. Davies, H. Li, Twelve years of SAMtools and BCFtools. GigaScience. 10 (2021), doi:10.1093/gigascience/giab008.

52. N. J. Loman, J. Quick, J. T. Simpson, A complete bacterial genome assembled de novo using only nanopore sequencing data. Nat. Methods. 12, 733–735 (2015).

53. P. Lindenbaum, JVarkit: java-based utilities for Bioinformatics (2015), doi:10.6084/m9.figshare.1425030.v1.

54. M. L. Waskom, seaborn: statistical data visualization. J. Open Source Softw. 6, 3021 (2021).

55. Schrödinger Release 2021-2: Prime, Schrödinger, LLC, New York, NY, 2021.

56. Schrödinger Release 2021-2: MacroModel, Schrödinger, LLC, New York, NY, 2021.

57. I. Berger, D. J. Fitzgerald, T. J. Richmond, Baculovirus expression system for heterologous multiprotein complexes. Nat. Biotechnol. 22, 1583–1587 (2004).

58. C. Bieniossek, T. J. Richmond, I. Berger, Curr. Protoc. Protein Sci., in press, doi:10.1002/0471140864.ps0520s51.

