## Supplemental Figures and Tables for "Distinct genetic determinants and mechanisms of SARS-CoV-2 resistance to remdesivir"

Supplementary Figures:

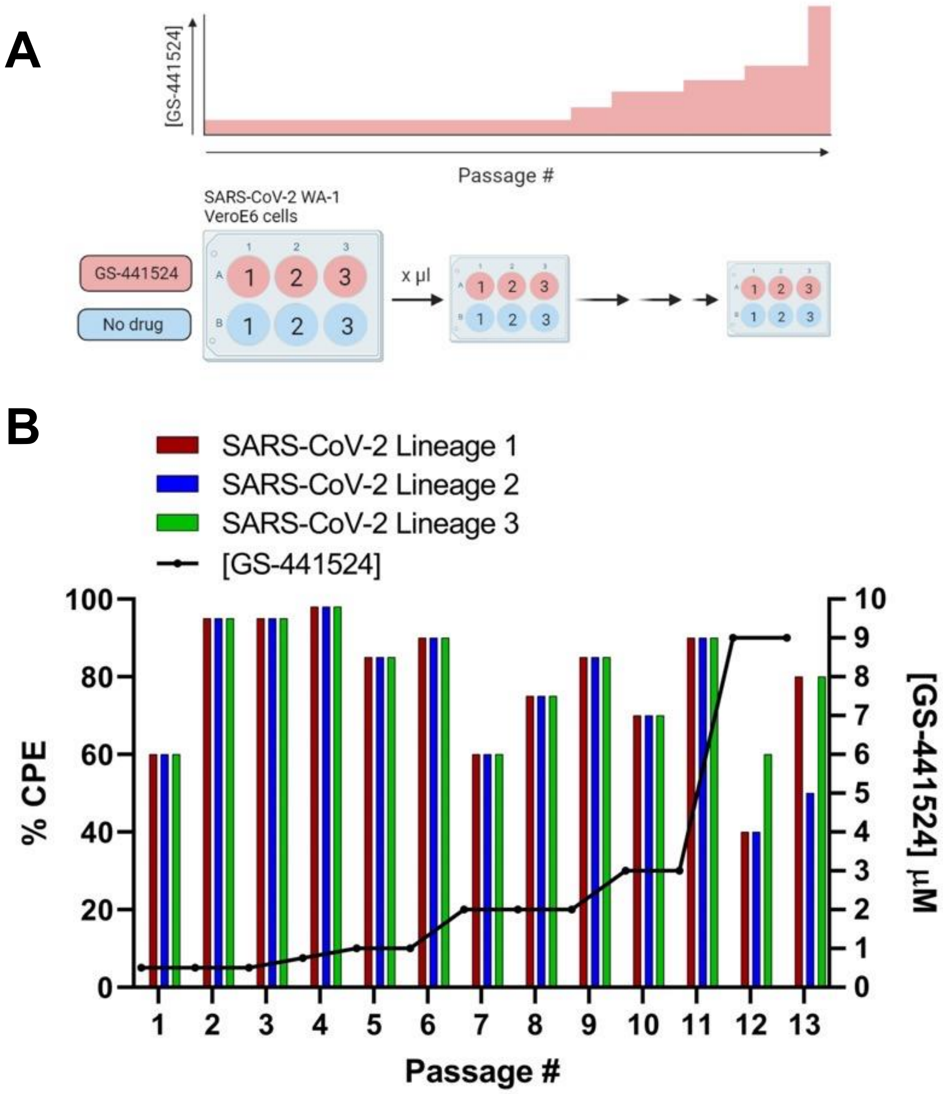


**Passage 13 Lineage 1 & 3: 9 µM**
**Passage 13 Lineage 2: 3 µM**

**Fig. S1. Serial passaging of SARS-CoV-2.** SARS-CoV-2 WA-1 isolate (culture passage 5) was additionally passaged 13x in the presence of GS-441524 or vehicle (DMSO) in VeroE6 cells, three separate lineages each. **(A)** Schematic representation of viral passaging. **(B)** Percentage of cell monolayer with viral cytopathic effect (CPE) is shown on left Y axis and GS-441524 concentration on right y axis.

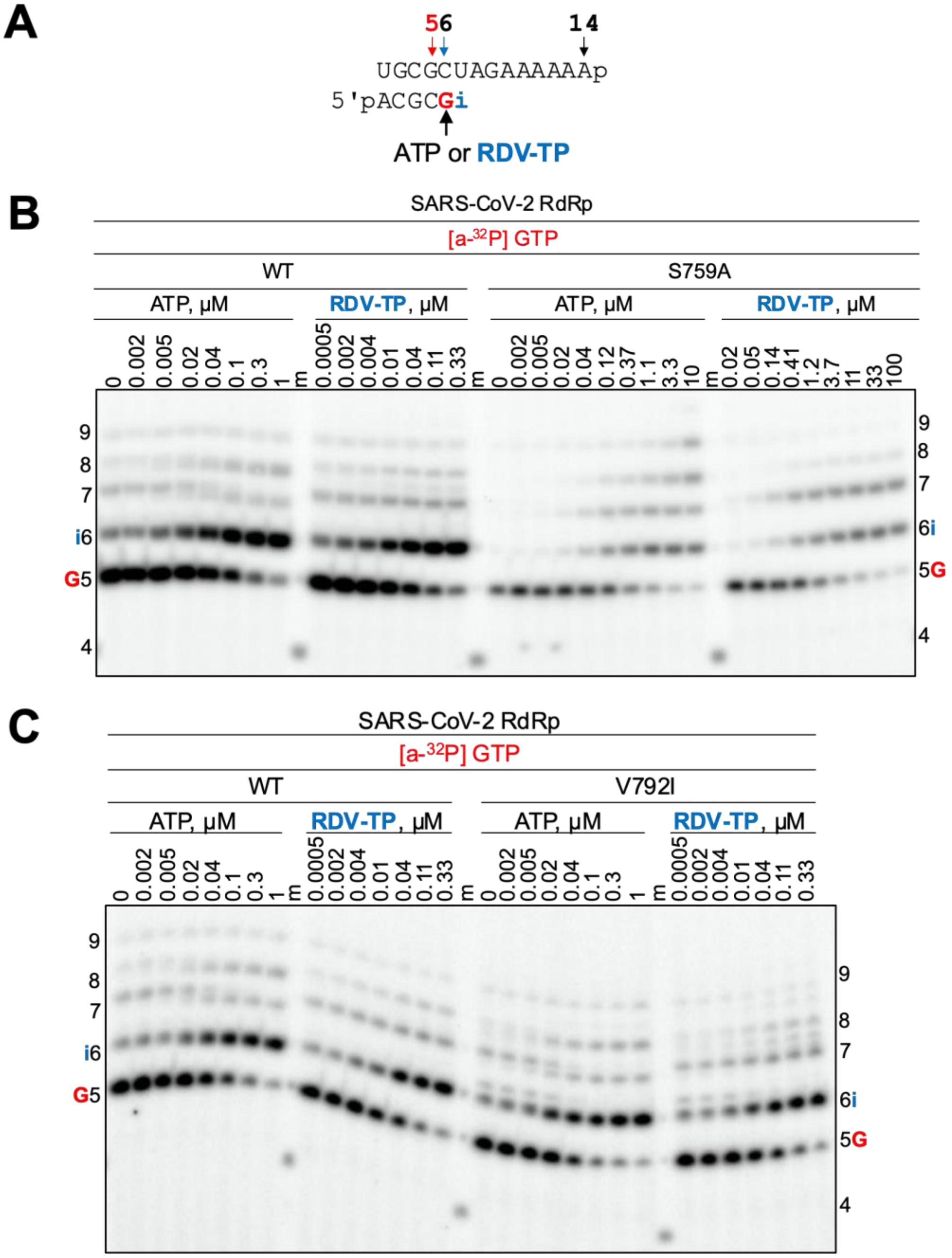


**Fig. S2. Selective incorporation of RDV-TP by WT and mutant S759A, and V792I SARS-CoV-2 RdRp complexes. (A)** RNA primer/template sequence used to determine the efficiency of ATP or RDV-TP incorporation at position 6 (i). G indicates incorporation of [α-$^{32}$P] GTP at position 5 (red). **(B)** Migration patterns of the products of ATP or RDV-TP incorporation reactions with WT and S759A SARS-CoV-2 RdRp complexes. Main products emerge at position 6 and mismatches are seen at positions 7, 8, and 9. The 5'-$^{32}$P-labeled 4-nt primer (4) is used as a size marker (m). **(C)** Migration patterns of the reactions with WT and V792I SARS-CoV-2 RdRp complexes under conditions as described above.

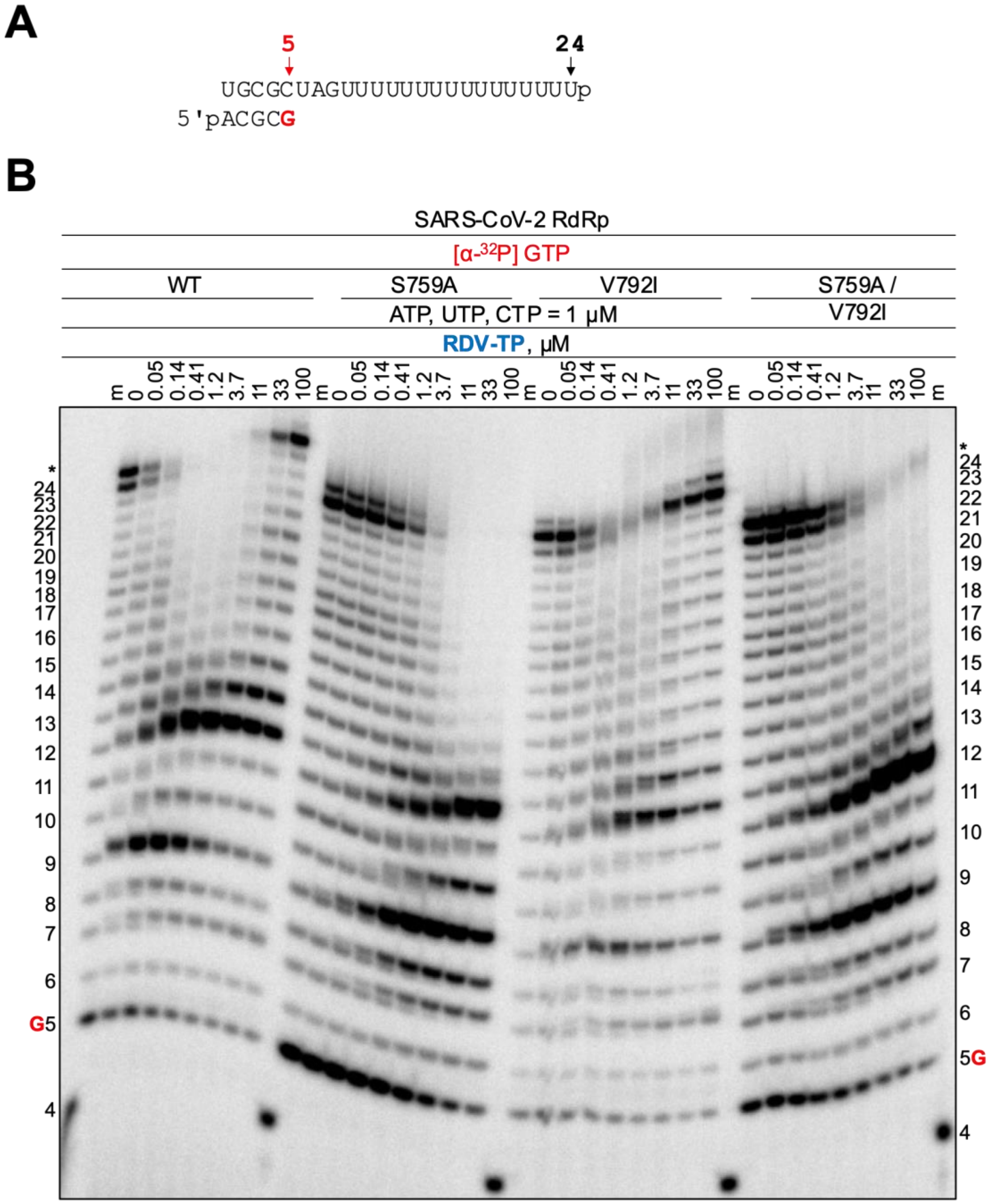


**Fig. S3. Competition between RDV-TP and natural NTPs in SARS-CoV-2 WT and mutant S759A, V792I, and S759A/V792I RdRp complexes. (A)** RNA primer/template sequence used is shown. **(B)** NTPs (ATP, UTP, and CTP) were supplemented at a constant concentration per reaction while RDV-TP concentrations varied as indicated. Denaturing PAGE migration patterns of RNA products are shown. G(red) indicates the incorporation of [α-$^{32}$P] GTP at position 5 and 4 indicates the migration pattern of 5'-$^{32}$P-labeled 4-nt primer is used as a size marker (m). The asterisk (*) indicates products formed due to terminal transferase activity.

**Table S1. $EC_{50}$ and fold change for GS-441524 and vehicle-passaged virus lineages.**

| Passage | $EC_{50}$ [mM] n=2 | |
|---|---|---|
| **DMSO** | | |
| P9 Lineage 1 | 0.36 | |
| P9 Lineage 2 | 0.34 | |
| P9 Lineage 3 | 0.24 | |
| P13 Lineage 1 | 0.41 | |
| P13 Lineage 2 | 0.39 | |
| P13 Lineage 3 | 0.39 | |
| **GS-441524** | | **Fold Change** |
| P9 Lineage 1 | 0.94 | 2.6 |
| P9 Lineage 2 | 0.56 | 1.5 |
| P9 Lineage 3 | 0.40 | 1.7 |
| P13 Lineage 1 | 4.22 | 10.4 |
| P13 Lineage 2 | 1.11 | 2.7 |
| P13 Lineage 3 | 3.27 | 8.0 |
| PP nsp12-V792I | 1.06 | 2.6 |
| PP nsp12-S759A/V792I | 2.97 | 7.3 |

SARS-CoV-2 was passaged 13 times in increasing concentrations of GS-441524 or vehicle (DMSO). Fold-change represents RDV $EC_{50}$ ratio of drug-passaged to vehicle-passaged virus tested in A549-hACE2 cells.

**Table S2.** Number of times the nsp12 amino acid substitutions were detected in SARS-CoV-2 sequences deposited to GISAID database

| **Residue Substitution** | **Mutation Frequency in all Sequences, % (N)** [a] | **Mutation Frequency in Omicron Sequences, % (N)** [b] | **Mutation Frequency in Delta Sequences, % (N)** [c] |
|---|---|---|---|
| V166A | 0.002% (107) | 0 | 0.0007% (26) |
| N198S | 0.013% (845) | 0.0007% (1) | 0.015% (580) |
| S759A | 0.000% (1) | 0 | 0 |
| V792I | 0.002% (105) | 0 | 0.0002% (8) |
| C799F | 0.001% (70) | 0 | 0.0006% (24) |
| C799R | 0.000% (4) | 0 | 0.0000% (1) |

[a] Total N = 6,741,412 genomes in GISAID on 01/04/22 including Omicron and Delta variants (https://www.gisaid.org).
[b] N = 138,061 Omicron genomes in GISAID on 01/04/22 (https://www.gisaid.org).
[c] N = 3,924,580 Delta genomes in GISAID on 01/04/22 (https://www.gisaid.org).
