## Supplementary material for "Distinct genetic determinants and mechanisms of SARS-CoV-2 resistance to remdesivir": Data file S1

Stock: SARS-CoV-2 WA-1 P5

P6
DMSO Lineage 1
DMSO Lineage 2
DMSO Lineage 3
Lineage 1
Lineage 2
Lineage 3

P9
DMSO Lineage 1
DMSO Lineage 2
DMSO Lineage 3
Lineage 1
Lineage 2
Lineage 3

P13
DMSO Lineage 1
DMSO Lineage 2
DMSO Lineage 3
Lineage 1
Lineage 2
Lineage 3

Plaque Picks
PP nsp12-V792I
PP nsp12-S759A/V792I

SARS-CoV-2 WA-1 P5

| Genome | Position | Reference | Variant | Frequency | Gene |
|---|---|---|---|---|---|
| MT020881.1 | 23606 | C | T | 0.251005942 | S protein |
| MT020881.1 | 23607 | G | T | 0.161503920 | S protein |
| MT020881.1 | 27761 | T | C | 0.150011556 | ORF7ab |
| MT020881.1 | 27758 | G | A | 0.149107837 | ORF7ab |
| MT020881.1 | 27760 | T | A | 0.148228285 | ORF7ab |
| MT020881.1 | 76 | T | A | 0.122589717 | 5UTR |
| MT020881.1 | 21784 | T | A | 0.104840574 | S protein |
| MT020881.1 | 23607 | G | A | 0.089976872 | S protein |
| MT020881.1 | 22482 | C | T | 0.081310239 | S protein |
| MT020881.1 | 8498 | G | T | 0.075248888 | nsp3 |
| MT020881.1 | 17827 | C | A | 0.074706148 | nsp13 |
| MT020881.1 | 23525 | C | T | 0.068291194 | S protein |
| MT020881.1 | 23618 | A | G | 0.051795811 | S protein |
| MT020881.1 | 23606 | C | G | 0.047776187 | S protein |
| MT020881.1 | 14679 | T | C | 0.040207705 | nsp12 |
| MT020881.1 | 7028 | T | G | 0.028092765 | nsp3 |
| MT020881.1 | 15771 | T | C | 0.027986887 | nsp12 |
| MT020881.1 | 2145 | T | C | 0.025855484 | nsp2 |
| MT020881.1 | 15357 | T | C | 0.025596155 | nsp12 |
| MT020881.1 | 13422 | T | C | 0.024352129 | nsp10 |
| MT020881.1 | 11566 | T | A | 0.024234779 | nsp6 |
| MT020881.1 | 28853 | T | A | 0.022593737 | N protein |
| MT020881.1 | 3315 | C | T | 0.022557838 | nsp3 |
| MT020881.1 | 22795 | T | C | 0.021536480 | S protein |
| MT020881.1 | 17142 | T | C | 0.020273097 | nsp13 |
| MT020881.1 | 15771 | T | A | 0.020224582 | nsp12 |
| MT020881.1 | 8554 | T | C | 0.019951230 | nsp3 |
| MT020881.1 | 10451 | A | G | 0.017072229 | nsp5 |
| MT020881.1 | 26542 | C | T | 0.016632726 | M protein |
| MT020881.1 | 1420 | C | T | 0.015591539 | nsp2 |
| MT020881.1 | 559 | T | C | 0.015247879 | nsp1 |
| MT020881.1 | 1963 | T | A | 0.014656659 | nsp2 |
| MT020881.1 | 22990 | T | C | 0.014584609 | S protein |
| MT020881.1 | 19044 | T | A | 0.014502629 | nsp14 |
| MT020881.1 | 5457 | C | T | 0.014492353 | nsp3 |
| MT020881.1 | 15909 | T | C | 0.014149534 | nsp12 |
| MT020881.1 | 14679 | T | A | 0.013949465 | nsp12 |
| MT020881.1 | 22673 | T | C | 0.013818317 | S protein |
| MT020881.1 | 7562 | T | C | 0.013508950 | nsp3 |
| MT020881.1 | 7527 | T | C | 0.013490816 | nsp3 |
| MT020881.1 | 19896 | T | C | 0.013454902 | nsp15 |
| MT020881.1 | 1963 | T | G | 0.013380729 | nsp2 |
| MT020881.1 | 13947 | A | G | 0.013379266 | nsp12 |

| | | | | | |
|---|---|---|---|---|---|
| MT020881.1 | 5389 | T | C | 0.013313374 | nsp3 |
| MT020881.1 | 15771 | T | G | 0.012976605 | nsp12 |
| MT020881.1 | 2473 | A | G | 0.012963443 | nsp2 |
| MT020881.1 | 2860 | T | C | 0.012644054 | nsp3 |
| MT020881.1 | 23128 | A | G | 0.012287512 | S protein |
| MT020881.1 | 9812 | T | A | 0.012184508 | nsp4 |
| MT020881.1 | 11083 | G | T | 0.012152502 | nsp6 |
| MT020881.1 | 13542 | T | G | 0.011980268 | nsp12 |
| MT020881.1 | 6405 | T | C | 0.011788507 | nsp3 |
| MT020881.1 | 1683 | T | C | 0.011652884 | nsp2 |
| MT020881.1 | 5207 | T | C | 0.011652188 | nsp3 |
| MT020881.1 | 4147 | T | C | 0.011618817 | nsp3 |
| MT020881.1 | 20471 | A | G | 0.011540121 | nsp15 |
| MT020881.1 | 3625 | A | G | 0.011387749 | nsp3 |
| MT020881.1 | 7474 | T | C | 0.011198152 | nsp3 |
| MT020881.1 | 16576 | T | C | 0.011107297 | nsp13 |
| MT020881.1 | 1442 | T | A | 0.011059060 | nsp2 |
| MT020881.1 | 4507 | T | C | 0.011007469 | nsp3 |
| MT020881.1 | 1783 | T | C | 0.010718948 | nsp2 |
| MT020881.1 | 11223 | T | C | 0.010531520 | nsp6 |
| MT020881.1 | 16646 | C | T | 0.010379287 | nsp13 |
| MT020881.1 | 2719 | T | C | 0.010294529 | nsp2 |
| MT020881.1 | 8383 | T | C | 0.010267621 | nsp3 |
| MT020881.1 | 10620 | A | G | 0.010094735 | nsp5 |
| MT020881.1 | 13048 | T | C | 0.010009008 | nsp10 |
| MT020881.1 | 16209 | T | A | 0.010006447 | nsp12 |

DMSO P6 Lineage 1

| Genome | Position | Reference | Variant | Frequency | Gene |
|---|---|---|---|---|---|
| MT020881.1 | 21784 | T | A | 0.98635775 | S protein |
| MT020881.1 | 8498 | G | T | 0.97849026 | nsp3 |
| MT020881.1 | 23618 | A | G | 0.67948474 | S protein |
| MT020881.1 | 76 | T | A | 0.20414308 | 5UTR |
| MT020881.1 | 1820 | G | A | 0.16050798 | nsp2 |
| MT020881.1 | 1426 | C | T | 0.12027195 | nsp2 |
| MT020881.1 | 23607 | G | A | 0.11722839 | S protein |
| MT020881.1 | 23525 | C | T | 0.09437509 | S protein |
| MT020881.1 | 29529 | C | T | 0.05475789 | N protein |
| MT020881.1 | 28285 | T | C | 0.0447518 | N protein |
| MT020881.1 | 27761 | T | C | 0.04382594 | ORF7ab |
| MT020881.1 | 27760 | T | A | 0.04322871 | ORF7ab |
| MT020881.1 | 27758 | G | A | 0.04275513 | ORF7ab |
| MT020881.1 | 10162 | T | G | 0.04155673 | nsp5 |
| MT020881.1 | 2083 | T | A | 0.04139651 | nsp2 |
| MT020881.1 | 23606 | C | T | 0.03542332 | S protein |
| MT020881.1 | 23616 | G | A | 0.03128747 | S protein |
| MT020881.1 | 15771 | T | C | 0.02510849 | nsp12 |
| MT020881.1 | 14679 | T | C | 0.02440581 | nsp12 |
| MT020881.1 | 9004 | T | C | 0.02419675 | nsp4 |
| MT020881.1 | 15357 | T | C | 0.02355528 | nsp12 |
| MT020881.1 | 11566 | T | A | 0.02326365 | nsp6 |
| MT020881.1 | 7764 | C | T | 0.02134886 | nsp3 |
| MT020881.1 | 15835 | T | C | 0.0204111 | nsp12 |
| MT020881.1 | 11448 | A | C | 0.01948558 | nsp6 |
| MT020881.1 | 13422 | T | C | 0.01896977 | nsp10 |
| MT020881.1 | 559 | T | C | 0.01720335 | nsp1 |
| MT020881.1 | 2145 | T | C | 0.01705321 | nsp2 |
| MT020881.1 | 23607 | G | T | 0.01576101 | S protein |
| MT020881.1 | 10451 | A | G | 0.01509434 | nsp5 |
| MT020881.1 | 27888 | A | C | 0.01442052 | ORF7ab |
| MT020881.1 | 2473 | A | G | 0.01435949 | nsp2 |
| MT020881.1 | 7562 | T | C | 0.01426719 | nsp3 |
| MT020881.1 | 20135 | T | C | 0.01397064 | nsp15 |
| MT020881.1 | 16576 | T | C | 0.01388038 | nsp13 |
| MT020881.1 | 15909 | T | C | 0.01375776 | nsp12 |
| MT020881.1 | 17142 | T | C | 0.01338944 | nsp13 |
| MT020881.1 | 25267 | C | T | 0.01264515 | S protein |
| MT020881.1 | 7527 | T | C | 0.01183432 | nsp3 |
| MT020881.1 | 8383 | T | C | 0.01170836 | nsp3 |
| MT020881.1 | 5207 | T | C | 0.01169337 | nsp3 |

| | | | | | |
|---|---|---|---|---|---|
| MT020881.1 | 13542 | T | G | 0.01097026 | nsp12 |
| MT020881.1 | 19044 | T | A | 0.01072574 | nsp14 |
| MT020881.1 | 10620 | A | G | 0.01065283 | nsp5 |
| MT020881.1 | 22678 | A | G | 0.01045188 | S protein |
| MT020881.1 | 23606 | C | G | 0.01040889 | S protein |
| MT020881.1 | 19896 | T | C | 0.01033089 | nsp15 |
| MT020881.1 | 1045 | A | G | 0.01023535 | nsp2 |
| MT020881.1 | 1629 | A | G | 0.01017394 | nsp2 |
| MT020881.1 | 22114 | T | C | 0.01011402 | S protein |
| MT020881.1 | 5641 | A | C | 0.01005103 | nsp3 |

DMSO P6 Lineage 2

| Genome | Position | Reference | Variant | Frequency | Gene |
|---|---|---|---|---|---|
| MT020881.1 | 21784 | T | A | 0.63647023 | S protein |
| MT020881.1 | 23607 | G | T | 0.50393076 | S protein |
| MT020881.1 | 8498 | G | T | 0.46117112 | nsp3 |
| MT020881.1 | 22482 | C | T | 0.41329642 | S protein |
| MT020881.1 | 17827 | C | A | 0.39352009 | nsp13 |
| MT020881.1 | 26309 | C | A | 0.39025245 | E protein |
| MT020881.1 | 23525 | C | T | 0.19714466 | S protein |
| MT020881.1 | 23607 | G | A | 0.13618286 | S protein |
| MT020881.1 | 23618 | A | G | 0.12091566 | S protein |
| MT020881.1 | 27761 | T | C | 0.11563984 | ORF7ab |
| MT020881.1 | 27758 | G | A | 0.11297286 | ORF7ab |
| MT020881.1 | 27760 | T | A | 0.11275916 | ORF7ab |
| MT020881.1 | 76 | T | A | 0.10725851 | 5UTR |
| MT020881.1 | 23606 | C | T | 0.09218149 | S protein |
| MT020881.1 | 28892 | C | T | 0.05017395 | N protein |
| MT020881.1 | 19547 | C | T | 0.03271509 | nsp14 |
| MT020881.1 | 26490 | T | G | 0.03243706 | NA |
| MT020881.1 | 23467 | T | G | 0.03075518 | S protein |
| MT020881.1 | 26542 | C | T | 0.02817749 | M protein |
| MT020881.1 | 17748 | T | A | 0.02505785 | nsp13 |
| MT020881.1 | 26358 | A | C | 0.02255686 | E protein |
| MT020881.1 | 29700 | A | G | 0.0225233 | 3UTR |
| MT020881.1 | 23689 | T | G | 0.02048865 | S protein |
| MT020881.1 | 28853 | T | A | 0.01992257 | N protein |
| MT020881.1 | 26183 | A | G | 0.01962206 | ORF3a |
| MT020881.1 | 23606 | C | G | 0.01766746 | S protein |
| MT020881.1 | 26353 | C | T | 0.01748067 | E protein |
| MT020881.1 | 10335 | C | T | 0.01635646 | nsp5 |
| MT020881.1 | 23672 | T | C | 0.01615536 | S protein |
| MT020881.1 | 13542 | T | G | 0.01566179 | nsp12 |
| MT020881.1 | 10162 | T | G | 0.01253396 | nsp5 |
| MT020881.1 | 1442 | T | G | 0.01203834 | nsp2 |
| MT020881.1 | 8554 | T | C | 0.01051679 | nsp3 |
| MT020881.1 | 19044 | T | A | 0.01029537 | nsp14 |

DMSO P6 Lineage 3

| Genome | Position | Reference | Variant | Frequency | Gene |
|---|---|---|---|---|---|
| MT020881.1 | 21784 | T | A | 0.99339493 | S protein |
| MT020881.1 | 8498 | G | T | 0.98150943 | nsp3 |
| MT020881.1 | 23618 | A | G | 0.71515911 | S protein |
| MT020881.1 | 76 | T | A | 0.23834247 | 5UTR |
| MT020881.1 | 27761 | T | C | 0.12697839 | ORF7ab |
| MT020881.1 | 27758 | G | A | 0.12387231 | ORF7ab |
| MT020881.1 | 27760 | T | A | 0.12379683 | ORF7ab |
| MT020881.1 | 23525 | C | T | 0.11096221 | S protein |
| MT020881.1 | 23606 | C | T | 0.10347138 | S protein |
| MT020881.1 | 23607 | G | A | 0.05314169 | S protein |
| MT020881.1 | 18556 | A | G | 0.04888731 | nsp14 |
| MT020881.1 | 19182 | A | G | 0.04132139 | nsp14 |
| MT020881.1 | 15357 | T | C | 0.0302028 | nsp12 |
| MT020881.1 | 11083 | G | T | 0.02875716 | nsp6 |
| MT020881.1 | 22303 | T | C | 0.0246724 | S protein |
| MT020881.1 | 10162 | T | G | 0.02417176 | nsp5 |
| MT020881.1 | 13099 | A | G | 0.02171937 | nsp10 |
| MT020881.1 | 28096 | A | T | 0.02110714 | ORF8 |
| MT020881.1 | 28853 | T | A | 0.02046345 | N protein |
| MT020881.1 | 29806 | A | G | 0.01960722 | 3UTR |
| MT020881.1 | 25526 | G | T | 0.0177885 | ORF3a |
| MT020881.1 | 8554 | T | C | 0.01715497 | nsp3 |
| MT020881.1 | 10451 | A | G | 0.01686747 | nsp5 |
| MT020881.1 | 26341 | A | C | 0.01555957 | E protein |
| MT020881.1 | 22114 | T | C | 0.01542901 | S protein |
| MT020881.1 | 22678 | A | G | 0.01493975 | S protein |
| MT020881.1 | 15771 | T | A | 0.01435072 | nsp12 |
| MT020881.1 | 13542 | T | G | 0.01417454 | nsp12 |
| MT020881.1 | 19044 | T | A | 0.01367376 | nsp14 |
| MT020881.1 | 8821 | A | G | 0.01356064 | nsp4 |
| MT020881.1 | 1963 | T | A | 0.01311855 | nsp2 |
| MT020881.1 | 2473 | A | G | 0.01244946 | nsp2 |
| MT020881.1 | 13947 | A | G | 0.01230001 | nsp12 |
| MT020881.1 | 16939 | T | C | 0.01205163 | nsp13 |
| MT020881.1 | 916 | A | C | 0.01199588 | nsp2 |
| MT020881.1 | 8383 | T | C | 0.01198288 | nsp3 |
| MT020881.1 | 29386 | C | A | 0.01190859 | N protein |
| MT020881.1 | 23607 | G | T | 0.01168779 | S protein |
| MT020881.1 | 4291 | A | G | 0.01126526 | nsp3 |
| MT020881.1 | 1442 | T | A | 0.01043552 | nsp2 |
| MT020881.1 | 4507 | T | C | 0.01022854 | nsp3 |
| MT020881.1 | 7562 | T | C | 0.01017474 | nsp3 |
| MT020881.1 | 16576 | T | C | 0.01002633 | nsp13 |

GS-441524 P6 Lineage 1

| Genome | Position | Reference | Variant | Frequency | Gene |
|---|---|---|---|---|---|
| MT020881.1 | 23607 | G | A | 0.926568519 | S protein |
| MT020881.1 | 14033 | A | G | 0.871043376 | nsp12 |
| MT020881.1 | 76 | T | A | 0.169581886 | 5UTR |
| MT020881.1 | 23607 | G | T | 0.135638809 | S protein |
| MT020881.1 | 8498 | G | T | 0.058857809 | nsp3 |
| MT020881.1 | 23606 | C | T | 0.042932004 | S protein |
| MT020881.1 | 27761 | T | C | 0.041871699 | ORF7ab |
| MT020881.1 | 27760 | T | A | 0.041425054 | ORF7ab |
| MT020881.1 | 27758 | G | A | 0.041032396 | ORF7ab |
| MT020881.1 | 15357 | T | C | 0.029446972 | nsp12 |
| MT020881.1 | 15771 | T | C | 0.028234168 | nsp12 |
| MT020881.1 | 28306 | T | C | 0.028014629 | N protein |
| MT020881.1 | 1204 | C | T | 0.027839077 | nsp2 |
| MT020881.1 | 13422 | T | C | 0.022599663 | nsp10 |
| MT020881.1 | 13542 | T | G | 0.021770266 | nsp12 |
| MT020881.1 | 23618 | A | G | 0.0206395 | S protein |
| MT020881.1 | 17142 | T | C | 0.018315018 | nsp13 |
| MT020881.1 | 22678 | A | G | 0.017551206 | S protein |
| MT020881.1 | 3625 | A | G | 0.016523673 | nsp3 |
| MT020881.1 | 17146 | A | T | 0.016331894 | nsp13 |
| MT020881.1 | 14679 | T | A | 0.015176715 | nsp12 |
| MT020881.1 | 22114 | T | C | 0.015154341 | S protein |
| MT020881.1 | 8554 | T | C | 0.015091116 | nsp3 |
| MT020881.1 | 26234 | A | T | 0.014755982 | NA |
| MT020881.1 | 2473 | A | G | 0.014449391 | nsp2 |
| MT020881.1 | 26351 | C | T | 0.014078297 | E protein |
| MT020881.1 | 20135 | T | C | 0.013094925 | nsp15 |
| MT020881.1 | 1963 | T | G | 0.01307914 | nsp2 |
| MT020881.1 | 1442 | T | A | 0.01305483 | nsp2 |
| MT020881.1 | 25704 | T | A | 0.013052819 | ORF3a |
| MT020881.1 | 10620 | A | G | 0.012960437 | nsp5 |
| MT020881.1 | 19896 | T | C | 0.01271515 | nsp15 |
| MT020881.1 | 19044 | T | A | 0.012601078 | nsp14 |
| MT020881.1 | 15909 | T | C | 0.012594841 | nsp12 |
| MT020881.1 | 25923 | T | C | 0.012477741 | ORF3a |
| MT020881.1 | 1963 | T | A | 0.01227629 | nsp2 |
| MT020881.1 | 13947 | A | G | 0.011625654 | nsp12 |
| MT020881.1 | 7562 | T | C | 0.011242433 | nsp3 |
| MT020881.1 | 5389 | T | C | 0.011203814 | nsp3 |
| MT020881.1 | 9812 | T | A | 0.011030972 | nsp4 |
| MT020881.1 | 25201 | A | G | 0.010951498 | S protein |
| MT020881.1 | 1442 | T | G | 0.010271903 | nsp2 |

| | |
|---|---|
| 21563 | 25384 |

GS-441524 P6 Lineage 2

| Genome | Position | Reference | Variant | Frequency | Gene |
|---|---|---|---|---|---|
| MT020881.1 | 21784 | T | A | 0.91906532 | S protein |
| MT020881.1 | 8498 | G | T | 0.75733545 | nsp3 |
| MT020881.1 | 1820 | G | A | 0.56241215 | nsp2 |
| MT020881.1 | 23607 | G | T | 0.54546146 | S protein |
| MT020881.1 | 23618 | A | G | 0.23734912 | S protein |
| MT020881.1 | 76 | T | A | 0.17664461 | 5UTR |
| MT020881.1 | 27761 | T | C | 0.17412326 | ORF7ab |
| MT020881.1 | 27760 | T | A | 0.16809074 | ORF7ab |
| MT020881.1 | 27758 | G | A | 0.16500954 | ORF7ab |
| MT020881.1 | 12297 | A | T | 0.15748031 | nsp8 |
| MT020881.1 | 23606 | C | T | 0.11301006 | S protein |
| MT020881.1 | 23607 | G | A | 0.10545745 | S protein |
| MT020881.1 | 22320 | A | G | 0.05955077 | S protein |
| MT020881.1 | 23525 | C | T | 0.05578677 | S protein |
| MT020881.1 | 20895 | A | G | 0.03843302 | nsp16 |
| MT020881.1 | 14679 | T | C | 0.03723643 | nsp12 |
| MT020881.1 | 11566 | T | A | 0.03280543 | nsp6 |
| MT020881.1 | 15357 | T | C | 0.02971188 | nsp12 |
| MT020881.1 | 26351 | C | T | 0.02970398 | E protein |
| MT020881.1 | 15771 | T | C | 0.02804456 | nsp12 |
| MT020881.1 | 14033 | A | G | 0.0231272 | nsp12 |
| MT020881.1 | 10162 | T | G | 0.02260327 | nsp5 |
| MT020881.1 | 26261 | C | T | 0.02195275 | E protein |
| MT020881.1 | 13542 | T | G | 0.02011096 | nsp12 |
| MT020881.1 | 2057 | A | C | 0.01999504 | nsp2 |
| MT020881.1 | 7562 | T | C | 0.01792115 | nsp3 |
| MT020881.1 | 8554 | T | C | 0.01705426 | nsp3 |
| MT020881.1 | 26258 | T | G | 0.01544618 | E protein |
| MT020881.1 | 26234 | A | T | 0.01536865 | NA |
| MT020881.1 | 15909 | T | C | 0.01526447 | nsp12 |
| MT020881.1 | 7527 | T | C | 0.01520468 | nsp3 |
| MT020881.1 | 16576 | T | C | 0.01493411 | nsp13 |
| MT020881.1 | 20135 | T | C | 0.01437186 | nsp15 |
| MT020881.1 | 2473 | A | G | 0.01419142 | nsp2 |
| MT020881.1 | 25923 | T | C | 0.01402276 | ORF3a |
| MT020881.1 | 19044 | T | A | 0.01366897 | nsp14 |
| MT020881.1 | 13947 | A | G | 0.01309586 | nsp12 |
| MT020881.1 | 14019 | T | C | 0.01305057 | nsp12 |
| MT020881.1 | 2860 | T | C | 0.01240164 | nsp3 |
| MT020881.1 | 22678 | A | G | 0.01226411 | S protein |
| MT020881.1 | 13936 | G | T | 0.0119258 | nsp12 |
| MT020881.1 | 7474 | T | C | 0.01178361 | nsp3 |
| MT020881.1 | 23606 | C | G | 0.01174284 | S protein |

| | | | | | |
|---|---|---|---|---|---|
| MT020881.1 | 16581 | A | G | 0.01154957 | nsp13 |
| MT020881.1 | 25704 | T | A | 0.01105638 | ORF3a |
| MT020881.1 | 19896 | T | C | 0.01069278 | nsp15 |
| MT020881.1 | 1963 | T | G | 0.01051673 | nsp2 |
| MT020881.1 | 6405 | T | C | 0.01043566 | nsp3 |
| MT020881.1 | 22673 | T | C | 0.01024452 | S protein |
| MT020881.1 | 22114 | T | C | 0.01016067 | S protein |
| MT020881.1 | 661 | T | C | 0.01003658 | nsp1 |

å

GS-441524 Lineage 3

| Genome | Position | Reference | Variant | Frequency | Gene |
|---|---|---|---|---|---|
| MT020881.1 | 21784 | T | A | 0.93804628 | S protein |
| MT020881.1 | 8498 | G | T | 0.59668267 | nsp3 |
| MT020881.1 | 23606 | C | T | 0.46067939 | S protein |
| MT020881.1 | 27761 | T | C | 0.40953832 | ORF7ab |
| MT020881.1 | 27758 | G | A | 0.40505768 | ORF7ab |
| MT020881.1 | 27760 | T | A | 0.40331286 | ORF7ab |
| MT020881.1 | 23618 | A | G | 0.19608917 | S protein |
| MT020881.1 | 76 | T | A | 0.15585224 | 5UTR |
| MT020881.1 | 10162 | T | G | 0.08818011 | nsp5 |
| MT020881.1 | 23616 | G | A | 0.08724114 | S protein |
| MT020881.1 | 23607 | G | A | 0.08523134 | S protein |
| MT020881.1 | 12747 | C | T | 0.08414171 | nsp9 |
| MT020881.1 | 23607 | G | T | 0.08093647 | S protein |
| MT020881.1 | 23525 | C | T | 0.05522892 | S protein |
| MT020881.1 | 7028 | T | G | 0.04795082 | nsp3 |
| MT020881.1 | 6106 | T | C | 0.03207889 | nsp3 |
| MT020881.1 | 15357 | T | C | 0.02777778 | nsp12 |
| MT020881.1 | 11566 | T | A | 0.02551834 | nsp6 |
| MT020881.1 | 14033 | A | G | 0.01860465 | nsp12 |
| MT020881.1 | 3625 | A | G | 0.01841141 | nsp3 |
| MT020881.1 | 13542 | T | G | 0.01816746 | nsp12 |
| MT020881.1 | 13422 | T | C | 0.01747612 | nsp10 |
| MT020881.1 | 821 | G | A | 0.01653026 | nsp2 |
| MT020881.1 | 15771 | T | A | 0.01618743 | nsp12 |
| MT020881.1 | 17142 | T | C | 0.0161584 | nsp13 |
| MT020881.1 | 19044 | T | A | 0.01607127 | nsp14 |
| MT020881.1 | 22678 | A | G | 0.01597824 | S protein |
| MT020881.1 | 26234 | A | T | 0.01569049 | NA |
| MT020881.1 | 26233 | G | C | 0.01538697 | NA |
| MT020881.1 | 18756 | G | A | 0.0146202 | nsp14 |
| MT020881.1 | 8884 | A | G | 0.01452885 | nsp4 |
| MT020881.1 | 25923 | T | C | 0.01366778 | ORF3a |
| MT020881.1 | 22114 | T | C | 0.01357418 | S protein |
| MT020881.1 | 10620 | A | G | 0.01314902 | nsp5 |
| MT020881.1 | 2473 | A | G | 0.01254921 | nsp2 |
| MT020881.1 | 13947 | A | G | 0.01183588 | nsp12 |
| MT020881.1 | 8554 | T | C | 0.01126634 | nsp3 |
| MT020881.1 | 20136 | A | G | 0.01119913 | nsp15 |
| MT020881.1 | 25704 | T | A | 0.011172 | ORF3a |
| MT020881.1 | 7474 | T | C | 0.010682 | nsp3 |
| MT020881.1 | 19896 | T | C | 0.01055856 | nsp15 |
| MT020881.1 | 12015 | T | G | 0.0104712 | nsp7 |
| MT020881.1 | 4291 | A | G | 0.01046671 | nsp3 |

| | | | | | |
|---|---|---|---|---|---|
| MT020881.1 | 1963 | T | G | 0.01014713 | nsp2 |
| MT020881.1 | 559 | T | C | 0.01011138 | nsp1 |
| MT020881.1 | 5389 | T | C | 0.01007484 | nsp3 |
| MT020881.1 | 667 | T | A | 0.01005333 | nsp1 |

DMSO P9 Lineage 1

| Genome | Position | Reference | Variant | Frequency | Gene |
| --- | --- | --- | --- | --- | --- |
| MT020881.1 | 21784 | T | A | 0.99882805 | S protein |
| MT020881.1 | 8498 | G | T | 0.9788961 | nsp3 |
| MT020881.1 | 23618 | A | G | 0.83984188 | S protein |
| MT020881.1 | 76 | T | A | 0.25878203 | 5UTR |
| MT020881.1 | 1820 | G | A | 0.14866809 | nsp2 |
| MT020881.1 | 23525 | C | T | 0.14494764 | S protein |
| MT020881.1 | 11448 | A | C | 0.12431863 | nsp6 |
| MT020881.1 | 7764 | C | T | 0.11845316 | nsp3 |
| MT020881.1 | 1426 | C | T | 0.11030856 | nsp2 |
| MT020881.1 | 23607 | G | A | 0.05871849 | S protein |
| MT020881.1 | 26387 | A | G | 0.05170814 | E protein |
| MT020881.1 | 10162 | T | G | 0.04917249 | nsp5 |
| MT020881.1 | 28285 | T | C | 0.04831466 | N protein |
| MT020881.1 | 27761 | T | C | 0.0479562 | ORF7ab |
| MT020881.1 | 27760 | T | A | 0.04764591 | ORF7ab |
| MT020881.1 | 27758 | G | A | 0.04748836 | ORF7ab |
| MT020881.1 | 2083 | T | A | 0.04199925 | nsp2 |
| MT020881.1 | 11566 | T | A | 0.03891736 | nsp6 |
| MT020881.1 | 14679 | T | C | 0.03827918 | nsp12 |
| MT020881.1 | 15771 | T | C | 0.03268982 | nsp12 |
| MT020881.1 | 15357 | T | C | 0.02809573 | nsp12 |
| MT020881.1 | 29529 | C | T | 0.02302013 | N protein |
| MT020881.1 | 10684 | T | G | 0.02278137 | nsp5 |
| MT020881.1 | 15771 | T | A | 0.02237876 | nsp12 |
| MT020881.1 | 23616 | G | A | 0.02128209 | S protein |
| MT020881.1 | 11477 | A | G | 0.01992032 | nsp6 |
| MT020881.1 | 13457 | T | C | 0.01961127 | nsp12 |
| MT020881.1 | 22990 | T | C | 0.01919492 | S protein |
| MT020881.1 | 3625 | A | G | 0.01914211 | nsp3 |
| MT020881.1 | 10451 | A | G | 0.01892272 | nsp5 |
| MT020881.1 | 24566 | C | G | 0.01837739 | S protein |
| MT020881.1 | 19374 | C | T | 0.01837711 | nsp14 |
| MT020881.1 | 13422 | T | C | 0.01774763 | nsp10 |
| MT020881.1 | 5457 | C | T | 0.0174522 | nsp3 |
| MT020881.1 | 11566 | T | G | 0.01622951 | nsp6 |
| MT020881.1 | 8554 | T | C | 0.01558586 | nsp3 |
| MT020881.1 | 22678 | A | G | 0.01552197 | S protein |
| MT020881.1 | 20135 | T | C | 0.01506192 | nsp15 |
| MT020881.1 | 25267 | C | T | 0.01459296 | S protein |
| MT020881.1 | 5389 | T | C | 0.0135807 | nsp3 |
| MT020881.1 | 10620 | A | G | 0.01314438 | nsp5 |
| MT020881.1 | 11083 | G | T | 0.01272412 | nsp6 |
| MT020881.1 | 19896 | T | C | 0.01218276 | nsp15 |

| | | | | | |
|---|---|---|---|---|---|
| MT020881.1 | 4147 | T | C | 0.01197227 | nsp3 |
| MT020881.1 | 14679 | T | A | 0.01189197 | nsp12 |
| MT020881.1 | 19044 | T | A | 0.01186381 | nsp14 |
| MT020881.1 | 13947 | A | G | 0.01183899 | nsp12 |
| MT020881.1 | 13542 | T | G | 0.01141455 | nsp12 |
| MT020881.1 | 559 | T | C | 0.01097217 | nsp1 |
| MT020881.1 | 6405 | T | C | 0.0108492 | nsp3 |
| MT020881.1 | 15909 | T | C | 0.01066695 | nsp12 |
| MT020881.1 | 8383 | T | C | 0.01064718 | nsp3 |
| MT020881.1 | 11223 | T | C | 0.01043841 | nsp6 |
| MT020881.1 | 16209 | T | A | 0.01026065 | nsp12 |
| MT020881.1 | 7922 | T | C | 0.01006957 | nsp3 |
| MT020881.1 | 2473 | A | G | 0.01 | nsp2 |

DMSO P9 Lineage 2

| Genome | Position | Reference | Variant | Frequency | Gene |
|---|---|---|---|---|---|
| MT020881.1 | 21784 | T | A | 0.9446024 | S protein |
| MT020881.1 | 8498 | G | T | 0.5190032 | nsp3 |
| MT020881.1 | 23618 | A | G | 0.50993507 | S protein |
| MT020881.1 | 27761 | T | C | 0.38709677 | ORF7ab |
| MT020881.1 | 27758 | G | A | 0.38306081 | ORF7ab |
| MT020881.1 | 27760 | T | A | 0.38061007 | ORF7ab |
| MT020881.1 | 19547 | C | T | 0.3450469 | nsp14 |
| MT020881.1 | 23525 | C | T | 0.24707727 | S protein |
| MT020881.1 | 76 | T | A | 0.19238716 | 5UTR |
| MT020881.1 | 22579 | T | C | 0.12591793 | S protein |
| MT020881.1 | 23607 | G | T | 0.10460125 | S protein |
| MT020881.1 | 26309 | C | A | 0.09482042 | E protein |
| MT020881.1 | 23607 | G | A | 0.08034938 | S protein |
| MT020881.1 | 28892 | C | T | 0.07759487 | N protein |
| MT020881.1 | 17827 | C | A | 0.05664344 | nsp13 |
| MT020881.1 | 22482 | C | T | 0.04747472 | S protein |
| MT020881.1 | 29700 | A | G | 0.04467326 | 3UTR |
| MT020881.1 | 23606 | C | G | 0.03857445 | S protein |
| MT020881.1 | 23606 | C | T | 0.0375829 | S protein |
| MT020881.1 | 26542 | C | T | 0.03385511 | M protein |
| MT020881.1 | 17748 | T | A | 0.0327578 | nsp13 |
| MT020881.1 | 23689 | T | G | 0.03132198 | S protein |
| MT020881.1 | 14679 | T | C | 0.02886094 | nsp12 |
| MT020881.1 | 11566 | T | A | 0.0287589 | nsp6 |
| MT020881.1 | 28853 | T | A | 0.02689215 | N protein |
| MT020881.1 | 23467 | T | G | 0.02664086 | S protein |
| MT020881.1 | 15357 | T | C | 0.02576452 | nsp12 |
| MT020881.1 | 15771 | T | C | 0.02507347 | nsp12 |
| MT020881.1 | 10451 | A | G | 0.01948643 | nsp5 |
| MT020881.1 | 10335 | C | T | 0.01842671 | nsp5 |
| MT020881.1 | 13422 | T | C | 0.01782375 | nsp10 |
| MT020881.1 | 8554 | T | C | 0.01739253 | nsp3 |
| MT020881.1 | 7562 | T | C | 0.01738495 | nsp3 |
| MT020881.1 | 26353 | C | T | 0.01677746 | E protein |
| MT020881.1 | 26183 | A | G | 0.01675061 | ORF3a |
| MT020881.1 | 2473 | A | G | 0.01634998 | nsp2 |
| MT020881.1 | 17142 | T | C | 0.01633406 | nsp13 |
| MT020881.1 | 26439 | G | A | 0.0163287 | E protein |
| MT020881.1 | 26895 | C | T | 0.01610926 | M protein |
| MT020881.1 | 21846 | C | T | 0.01523472 | S protein |
| MT020881.1 | 23672 | T | C | 0.01453415 | S protein |
| MT020881.1 | 20135 | T | C | 0.01416639 | nsp15 |
| MT020881.1 | 23616 | G | A | 0.01416547 | S protein |

| | | | | | |
|---|---|---|---|---|---|
| MT020881.1 | 22678 | A | G | 0.01400389 | S protein |
| MT020881.1 | 16576 | T | C | 0.01380368 | nsp13 |
| MT020881.1 | 21789 | C | T | 0.01374898 | S protein |
| MT020881.1 | 14033 | A | G | 0.01370601 | nsp12 |
| MT020881.1 | 15909 | T | C | 0.01358098 | nsp12 |
| MT020881.1 | 5389 | T | C | 0.01357597 | nsp3 |
| MT020881.1 | 7474 | T | C | 0.01324594 | nsp3 |
| MT020881.1 | 4507 | T | C | 0.0129413 | nsp3 |
| MT020881.1 | 8383 | T | C | 0.01280559 | nsp3 |
| MT020881.1 | 19044 | T | A | 0.01279958 | nsp14 |
| MT020881.1 | 13542 | T | G | 0.01276225 | nsp12 |
| MT020881.1 | 22114 | T | C | 0.01276034 | S protein |
| MT020881.1 | 22990 | T | C | 0.01244628 | S protein |
| MT020881.1 | 11522 | T | G | 0.0122449 | nsp6 |
| MT020881.1 | 11223 | T | C | 0.01216246 | nsp6 |
| MT020881.1 | 5207 | T | C | 0.01198596 | nsp3 |
| MT020881.1 | 1442 | T | A | 0.01194348 | nsp2 |
| MT020881.1 | 1963 | T | G | 0.01178276 | nsp2 |
| MT020881.1 | 1683 | T | C | 0.01142857 | nsp2 |
| MT020881.1 | 7527 | T | C | 0.01139114 | nsp3 |
| MT020881.1 | 19896 | T | C | 0.01069165 | nsp15 |
| MT020881.1 | 10620 | A | G | 0.01066018 | nsp5 |
| MT020881.1 | 16209 | T | A | 0.01058779 | nsp12 |
| MT020881.1 | 20471 | A | G | 0.01051172 | nsp15 |
| MT020881.1 | 26260 | T | G | 0.01047996 | E protein |
| MT020881.1 | 10210 | T | A | 0.01039247 | nsp5 |
| MT020881.1 | 13947 | A | G | 0.01026644 | nsp12 |
| MT020881.1 | 17895 | T | C | 0.01023783 | nsp13 |
| MT020881.1 | 12296 | C | A | 0.01018115 | nsp8 |
| MT020881.1 | 1045 | A | G | 0.01011556 | nsp2 |
| MT020881.1 | 22673 | T | C | 0.01003691 | S protein |

DMSO P9 Lineage 3

| Genome | Position | Reference | Variant | Frequency | Gene |
|---|---|---|---|---|---|
| MT020881.1 | 21784 | T | A | 0.99879178 | S protein |
| MT020881.1 | 8498 | G | T | 0.99043062 | nsp3 |
| MT020881.1 | 23618 | A | G | 0.91706195 | S protein |
| MT020881.1 | 23525 | C | T | 0.34959932 | S protein |
| MT020881.1 | 76 | T | A | 0.2161939 | 5UTR |
| MT020881.1 | 27761 | T | C | 0.1966881 | ORF7ab |
| MT020881.1 | 27758 | G | A | 0.1929416 | ORF7ab |
| MT020881.1 | 27760 | T | A | 0.19264939 | ORF7ab |
| MT020881.1 | 28096 | A | T | 0.10249115 | ORF8 |
| MT020881.1 | 25526 | G | T | 0.09689608 | ORF3a |
| MT020881.1 | 10162 | T | G | 0.08065476 | nsp5 |
| MT020881.1 | 13099 | A | G | 0.07999281 | nsp10 |
| MT020881.1 | 14679 | T | C | 0.03126503 | nsp12 |
| MT020881.1 | 15357 | T | C | 0.02995551 | nsp12 |
| MT020881.1 | 18556 | A | G | 0.02845482 | nsp14 |
| MT020881.1 | 19182 | A | G | 0.02758222 | nsp14 |
| MT020881.1 | 11566 | T | A | 0.02707334 | nsp6 |
| MT020881.1 | 26341 | A | C | 0.02514476 | E protein |
| MT020881.1 | 9429 | T | C | 0.02361061 | nsp4 |
| MT020881.1 | 11471 | C | T | 0.0221342 | nsp6 |
| MT020881.1 | 17142 | T | C | 0.02202119 | nsp13 |
| MT020881.1 | 23606 | C | T | 0.02166142 | S protein |
| MT020881.1 | 13422 | T | C | 0.02151451 | nsp10 |
| MT020881.1 | 4654 | T | G | 0.02056457 | nsp3 |
| MT020881.1 | 22303 | T | C | 0.02000565 | S protein |
| MT020881.1 | 15771 | T | A | 0.01968504 | nsp12 |
| MT020881.1 | 23607 | G | A | 0.01936188 | S protein |
| MT020881.1 | 7562 | T | C | 0.01851852 | nsp3 |
| MT020881.1 | 25443 | A | C | 0.01766962 | ORF3a |
| MT020881.1 | 8554 | T | C | 0.01747361 | nsp3 |
| MT020881.1 | 13542 | T | G | 0.01675714 | nsp12 |
| MT020881.1 | 10451 | A | G | 0.01646091 | nsp5 |
| MT020881.1 | 20135 | T | C | 0.01561514 | nsp15 |
| MT020881.1 | 8383 | T | C | 0.01560349 | nsp3 |
| MT020881.1 | 11379 | C | T | 0.01556886 | nsp6 |
| MT020881.1 | 16576 | T | C | 0.01438985 | nsp13 |
| MT020881.1 | 1783 | T | C | 0.01405928 | nsp2 |
| MT020881.1 | 22990 | T | C | 0.01344026 | S protein |
| MT020881.1 | 3625 | A | G | 0.01335784 | nsp3 |
| MT020881.1 | 22678 | A | G | 0.01313378 | S protein |
| MT020881.1 | 1820 | G | A | 0.01313307 | nsp2 |
| MT020881.1 | 7474 | T | C | 0.01245791 | nsp3 |
| MT020881.1 | 5207 | T | C | 0.01245634 | nsp3 |

| | | | | | |
|---|---|---|---|---|---|
| MT020881.1 | 19044 | T | A | 0.012379 | nsp14 |
| MT020881.1 | 7527 | T | C | 0.01189643 | nsp3 |
| MT020881.1 | 11083 | G | T | 0.01154618 | nsp6 |
| MT020881.1 | 916 | A | C | 0.0115277 | nsp2 |
| MT020881.1 | 1683 | T | C | 0.01110391 | nsp2 |
| MT020881.1 | 11522 | T | G | 0.01079812 | nsp6 |
| MT020881.1 | 22114 | T | C | 0.01071894 | S protein |
| MT020881.1 | 2473 | A | G | 0.01046106 | nsp2 |
| MT020881.1 | 28411 | C | T | 0.01039178 | N protein |

GS-441524 P9 Lineage 1

| Genome | Position | Reference | Variant | Frequency | Gene |
|---|---|---|---|---|---|
| MT020881.1 | 23607 | G | A | 0.98557875 | S protein |
| MT020881.1 | 14033 | A | G | 0.98522039 | nsp12 |
| MT020881.1 | 15836 | G | T | 0.36880402 | nsp12 |
| MT020881.1 | 13937 | T | C | 0.24981719 | nsp12 |
| MT020881.1 | 76 | T | A | 0.21189503 | 5UTR |
| MT020881.1 | 13847 | A | G | 0.13342371 | nsp12 |
| MT020881.1 | 9952 | T | G | 0.12187317 | nsp4 |
| MT020881.1 | 15296 | A | T | 0.11994643 | nsp12 |
| MT020881.1 | 3634 | C | A | 0.0986234 | nsp3 |
| MT020881.1 | 15222 | C | T | 0.04883466 | nsp12 |
| MT020881.1 | 986 | T | C | 0.04565635 | nsp2 |
| MT020881.1 | 13847 | A | T | 0.04075347 | nsp12 |
| MT020881.1 | 29839 | A | C | 0.03311743 | 3UTR |
| MT020881.1 | 15835 | T | C | 0.03125404 | nsp12 |
| MT020881.1 | 11566 | T | A | 0.02841638 | nsp6 |
| MT020881.1 | 5869 | C | T | 0.02740446 | nsp3 |
| MT020881.1 | 15357 | T | C | 0.02592004 | nsp12 |
| MT020881.1 | 26978 | T | C | 0.02540155 | M protein |
| MT020881.1 | 26151 | C | A | 0.02098913 | ORF3a |
| MT020881.1 | 28310 | C | A | 0.01956355 | N protein |
| MT020881.1 | 27980 | A | C | 0.01925094 | ORF8 |
| MT020881.1 | 19667 | A | G | 0.01923332 | nsp15 |
| MT020881.1 | 10451 | A | G | 0.01902959 | nsp5 |
| MT020881.1 | 22678 | A | G | 0.01765192 | S protein |
| MT020881.1 | 15837 | T | G | 0.01755611 | nsp12 |
| MT020881.1 | 13542 | T | G | 0.01709996 | nsp12 |
| MT020881.1 | 8498 | G | T | 0.01688771 | nsp3 |
| MT020881.1 | 14831 | G | A | 0.01683432 | nsp12 |
| MT020881.1 | 13422 | T | C | 0.01599271 | nsp10 |
| MT020881.1 | 15771 | T | A | 0.0154727 | nsp12 |
| MT020881.1 | 27629 | C | A | 0.01537301 | ORF7ab |
| MT020881.1 | 1963 | T | G | 0.01498783 | nsp2 |
| MT020881.1 | 8667 | A | C | 0.01495341 | nsp4 |
| MT020881.1 | 8554 | T | C | 0.01482159 | nsp3 |
| MT020881.1 | 15909 | T | C | 0.01442684 | nsp12 |
| MT020881.1 | 8821 | A | G | 0.01427532 | nsp4 |
| MT020881.1 | 8600 | G | T | 0.01383148 | nsp4 |
| MT020881.1 | 19044 | T | A | 0.01353948 | nsp14 |
| MT020881.1 | 13947 | A | G | 0.01308127 | nsp12 |
| MT020881.1 | 17142 | T | C | 0.01288924 | nsp13 |
| MT020881.1 | 1963 | T | A | 0.01258537 | nsp2 |
| MT020881.1 | 1442 | T | G | 0.01250499 | nsp2 |
| MT020881.1 | 4775 | A | G | 0.01232997 | nsp3 |

| | | | | | |
|---|---|---|---|---|---|
| MT020881.1 | 11566 | T | G | 0.01213884 | nsp6 |
| MT020881.1 | 2473 | A | G | 0.01179023 | nsp2 |
| MT020881.1 | 493 | T | A | 0.0117665 | nsp1 |
| MT020881.1 | 19896 | T | C | 0.01173099 | nsp15 |
| MT020881.1 | 4855 | A | G | 0.01151029 | nsp3 |
| MT020881.1 | 12296 | C | A | 0.01149696 | nsp8 |
| MT020881.1 | 28171 | C | A | 0.011281 | ORF8 |
| MT020881.1 | 16209 | T | A | 0.01113347 | nsp12 |
| MT020881.1 | 11233 | T | C | 0.01096983 | nsp6 |
| MT020881.1 | 1442 | T | A | 0.01092605 | nsp2 |
| MT020881.1 | 22034 | A | G | 0.0108246 | S protein |
| MT020881.1 | 10620 | A | G | 0.01082085 | nsp5 |
| MT020881.1 | 7922 | T | C | 0.01070639 | nsp3 |
| MT020881.1 | 11083 | G | T | 0.01058582 | nsp6 |
| MT020881.1 | 7562 | T | C | 0.01055855 | nsp3 |
| MT020881.1 | 6156 | A | C | 0.01017488 | nsp3 |
| MT020881.1 | 7009 | T | G | 0.0100306 | nsp3 |

GS-441524 P9 Lineage 2

| Genome | Position | Reference | Variant | Frequency | Gene |
|---|---|---|---|---|---|
| MT020881.1 | 21784 | T | A | 0.99853299 | S protein |
| MT020881.1 | 23607 | G | T | 0.96901318 | S protein |
| MT020881.1 | 8498 | G | T | 0.96363636 | nsp3 |
| MT020881.1 | 1820 | G | A | 0.85776136 | nsp2 |
| MT020881.1 | 15822 | G | T | 0.39120445 | nsp12 |
| MT020881.1 | 76 | T | A | 0.30951969 | 5UTR |
| MT020881.1 | 15835 | T | C | 0.27817864 | nsp12 |
| MT020881.1 | 9004 | T | C | 0.2725297 | nsp4 |
| MT020881.1 | 27761 | T | C | 0.23979331 | ORF7ab |
| MT020881.1 | 27758 | G | A | 0.23503984 | ORF7ab |
| MT020881.1 | 27760 | T | A | 0.23338718 | ORF7ab |
| MT020881.1 | 23607 | G | A | 0.16164629 | S protein |
| MT020881.1 | 25597 | T | A | 0.11132332 | ORF3a |
| MT020881.1 | 13847 | A | G | 0.10343619 | nsp12 |
| MT020881.1 | 13847 | A | C | 0.10167229 | nsp12 |
| MT020881.1 | 20895 | A | G | 0.10132043 | nsp16 |
| MT020881.1 | 13936 | G | T | 0.09978308 | nsp12 |
| MT020881.1 | 27919 | T | C | 0.03700183 | ORF8 |
| MT020881.1 | 15357 | T | C | 0.03006198 | nsp12 |
| MT020881.1 | 11566 | T | A | 0.02776367 | nsp6 |
| MT020881.1 | 26291 | G | A | 0.02282875 | E protein |
| MT020881.1 | 26380 | A | G | 0.0227346 | E protein |
| MT020881.1 | 3370 | T | C | 0.0207805 | nsp3 |
| MT020881.1 | 1102 | C | T | 0.02036598 | nsp2 |
| MT020881.1 | 26333 | C | T | 0.01990121 | E protein |
| MT020881.1 | 13422 | T | C | 0.01911119 | nsp10 |
| MT020881.1 | 29826 | T | C | 0.01860634 | 3UTR |
| MT020881.1 | 10451 | A | G | 0.01816096 | nsp5 |
| MT020881.1 | 13542 | T | G | 0.01752491 | nsp12 |
| MT020881.1 | 23618 | A | G | 0.01738262 | S protein |
| MT020881.1 | 22678 | A | G | 0.0172715 | S protein |
| MT020881.1 | 2676 | C | T | 0.01715519 | nsp2 |
| MT020881.1 | 20135 | T | C | 0.01681523 | nsp15 |
| MT020881.1 | 14032 | A | G | 0.0164827 | nsp12 |
| MT020881.1 | 8554 | T | C | 0.01602465 | nsp3 |
| MT020881.1 | 15909 | T | C | 0.015841 | nsp12 |
| MT020881.1 | 8821 | A | G | 0.01583194 | nsp4 |
| MT020881.1 | 29254 | G | A | 0.01528393 | N protein |
| MT020881.1 | 2473 | A | G | 0.01513145 | nsp2 |
| MT020881.1 | 13947 | A | G | 0.01500082 | nsp12 |
| MT020881.1 | 19044 | T | A | 0.01447999 | nsp14 |
| MT020881.1 | 1442 | T | G | 0.0142923 | nsp2 |
| MT020881.1 | 11566 | T | G | 0.01424648 | nsp6 |

| | | | | | |
|---|---|---|---|---|---|
| MT020881.1 | 14425 | C | T | 0.01423488 | nsp12 |
| MT020881.1 | 15771 | T | A | 0.01417065 | nsp12 |
| MT020881.1 | 22114 | T | C | 0.01395526 | S protein |
| MT020881.1 | 7562 | T | C | 0.01283226 | nsp3 |
| MT020881.1 | 5389 | T | C | 0.01269604 | nsp3 |
| MT020881.1 | 19896 | T | C | 0.0120499 | nsp15 |
| MT020881.1 | 6775 | T | A | 0.01200835 | nsp3 |
| MT020881.1 | 1963 | T | G | 0.01196429 | nsp2 |
| MT020881.1 | 1963 | T | A | 0.01161129 | nsp2 |
| MT020881.1 | 8090 | C | A | 0.01066531 | nsp3 |
| MT020881.1 | 6035 | A | C | 0.01060674 | nsp3 |
| MT020881.1 | 4291 | A | G | 0.01052026 | nsp3 |
| MT020881.1 | 4507 | T | C | 0.01009978 | nsp3 |
| MT020881.1 | 7527 | T | C | 0.01009317 | nsp3 |

GS-441524 P9 Lineage 3

| Genome | Position | Reference | Variant | Frequency | Gene |
|---|---|---|---|---|---|
| MT020881.1 | 21784 | T | A | 0.99883638 | S protein |
| MT020881.1 | 8498 | G | T | 0.89620293 | nsp3 |
| MT020881.1 | 23606 | C | T | 0.79840019 | S protein |
| MT020881.1 | 15814 | G | A | 0.77884959 | nsp12 |
| MT020881.1 | 27761 | T | C | 0.76967116 | ORF7ab |
| MT020881.1 | 27758 | G | A | 0.7693819 | ORF7ab |
| MT020881.1 | 27760 | T | A | 0.75892075 | ORF7ab |
| MT020881.1 | 10162 | T | G | 0.75278853 | nsp5 |
| MT020881.1 | 27635 | C | T | 0.1406565 | ORF7ab |
| MT020881.1 | 13847 | A | G | 0.13261104 | nsp12 |
| MT020881.1 | 21846 | C | T | 0.12215551 | S protein |
| MT020881.1 | 23586 | A | G | 0.12119686 | S protein |
| MT020881.1 | 23607 | G | T | 0.11151855 | S protein |
| MT020881.1 | 13847 | A | C | 0.0949957 | nsp12 |
| MT020881.1 | 29672 | T | A | 0.068874 | ORF10 |
| MT020881.1 | 29671 | A | T | 0.06855881 | ORF10 |
| MT020881.1 | 8392 | T | A | 0.03394589 | nsp3 |
| MT020881.1 | 15357 | T | C | 0.03287161 | nsp12 |
| MT020881.1 | 13542 | T | G | 0.0217845 | nsp12 |
| MT020881.1 | 15771 | T | C | 0.02027779 | nsp12 |
| MT020881.1 | 22678 | A | G | 0.01958454 | S protein |
| MT020881.1 | 13422 | T | C | 0.01919456 | nsp10 |
| MT020881.1 | 2145 | T | C | 0.01885566 | nsp2 |
| MT020881.1 | 1963 | T | A | 0.01755548 | nsp2 |
| MT020881.1 | 2473 | A | G | 0.01622181 | nsp2 |
| MT020881.1 | 3306 | T | C | 0.01584226 | nsp3 |
| MT020881.1 | 1963 | T | G | 0.01581683 | nsp2 |
| MT020881.1 | 17142 | T | C | 0.01558044 | nsp13 |
| MT020881.1 | 22114 | T | C | 0.0148617 | S protein |
| MT020881.1 | 19044 | T | A | 0.01469209 | nsp14 |
| MT020881.1 | 15771 | T | A | 0.01461833 | nsp12 |
| MT020881.1 | 9979 | C | T | 0.0143228 | nsp4 |
| MT020881.1 | 20135 | T | C | 0.01412883 | nsp15 |
| MT020881.1 | 13947 | A | G | 0.01397221 | nsp12 |
| MT020881.1 | 19896 | T | C | 0.01364777 | nsp15 |
| MT020881.1 | 11566 | T | G | 0.01360788 | nsp6 |
| MT020881.1 | 8554 | T | C | 0.01257373 | nsp3 |
| MT020881.1 | 1442 | T | A | 0.01181468 | nsp2 |
| MT020881.1 | 4501 | A | C | 0.01151436 | nsp3 |
| MT020881.1 | 1820 | G | A | 0.01141568 | nsp2 |
| MT020881.1 | 5687 | T | C | 0.01119171 | nsp3 |
| MT020881.1 | 16209 | T | A | 0.01118711 | nsp12 |
| MT020881.1 | 13949 | A | G | 0.01118128 | nsp12 |

| | | | | | |
|---|---|---|---|---|---|
| MT020881.1 | 7705 | A | G | 0.01102293 | nsp3 |
| MT020881.1 | 7562 | T | C | 0.0107168 | nsp3 |
| MT020881.1 | 1442 | T | G | 0.01063264 | nsp2 |
| MT020881.1 | 13685 | A | G | 0.01032153 | nsp12 |
| MT020881.1 | 25807 | T | A | 0.01027283 | ORF3a |
| MT020881.1 | 17886 | T | C | 0.01006935 | nsp13 |

DMSO P13 Lineage 1

| Genome | Position | Reference | Variant | Frequency | Gene |
|---|---|---|---|---|---|
| MT020881.1 | 21784 | T | A | 0.998872 | S protein |
| MT020881.1 | 8498 | G | T | 0.93435605 | nsp3 |
| MT020881.1 | 23618 | A | G | 0.93110944 | S protein |
| MT020881.1 | 23525 | C | T | 0.81191505 | S protein |
| MT020881.1 | 26387 | A | G | 0.58502477 | E protein |
| MT020881.1 | 76 | T | A | 0.29417782 | 5UTR |
| MT020881.1 | 11448 | A | C | 0.25219176 | nsp6 |
| MT020881.1 | 7764 | C | T | 0.19456812 | nsp3 |
| MT020881.1 | 22101 | A | C | 0.10349097 | S protein |
| MT020881.1 | 1820 | G | A | 0.08920976 | nsp2 |
| MT020881.1 | 1426 | C | T | 0.07470083 | nsp2 |
| MT020881.1 | 11477 | A | G | 0.06676558 | nsp6 |
| MT020881.1 | 11379 | C | T | 0.06371129 | nsp6 |
| MT020881.1 | 27761 | T | C | 0.05006731 | ORF7ab |
| MT020881.1 | 27760 | T | A | 0.04853342 | ORF7ab |
| MT020881.1 | 27758 | G | A | 0.0482662 | ORF7ab |
| MT020881.1 | 5457 | C | T | 0.0391907 | nsp3 |
| MT020881.1 | 10279 | C | T | 0.03840683 | nsp5 |
| MT020881.1 | 10162 | T | G | 0.03662122 | nsp5 |
| MT020881.1 | 28285 | T | C | 0.02847618 | N protein |
| MT020881.1 | 6987 | G | T | 0.02641879 | nsp3 |
| MT020881.1 | 11471 | C | T | 0.02539208 | nsp6 |
| MT020881.1 | 11566 | T | A | 0.02527973 | nsp6 |
| MT020881.1 | 13542 | T | G | 0.0230747 | nsp12 |
| MT020881.1 | 18377 | C | T | 0.02263877 | nsp14 |
| MT020881.1 | 10684 | T | G | 0.02171505 | nsp5 |
| MT020881.1 | 23607 | G | A | 0.02062235 | S protein |
| MT020881.1 | 19144 | T | G | 0.01997713 | nsp14 |
| MT020881.1 | 27307 | A | G | 0.01993015 | ORF6 |
| MT020881.1 | 13422 | T | C | 0.01742483 | nsp10 |
| MT020881.1 | 2083 | T | A | 0.01692923 | nsp2 |
| MT020881.1 | 26754 | G | A | 0.01629928 | M protein |
| MT020881.1 | 8554 | T | C | 0.0162069 | nsp3 |
| MT020881.1 | 23606 | C | T | 0.0153077 | S protein |
| MT020881.1 | 20135 | T | C | 0.01426686 | nsp15 |
| MT020881.1 | 16209 | T | A | 0.01408647 | nsp12 |
| MT020881.1 | 19044 | T | A | 0.01401901 | nsp14 |
| MT020881.1 | 15814 | G | A | 0.01382722 | nsp12 |
| MT020881.1 | 13457 | T | C | 0.01375539 | nsp12 |
| MT020881.1 | 2473 | A | G | 0.0135885 | nsp2 |
| MT020881.1 | 10165 | C | T | 0.01245033 | nsp5 |
| MT020881.1 | 11522 | T | G | 0.01234263 | nsp6 |
| MT020881.1 | 21789 | C | T | 0.01205457 | S protein |

| | | | | | |
|---|---|---|---|---|---|
| MT020881.1 | 22114 | T | C | 0.0119723 | S protein |
| MT020881.1 | 8821 | A | G | 0.01178571 | nsp4 |
| MT020881.1 | 4456 | C | T | 0.01167532 | nsp3 |
| MT020881.1 | 22678 | A | G | 0.01165976 | S protein |
| MT020881.1 | 25807 | T | A | 0.01162301 | ORF3a |
| MT020881.1 | 11223 | T | C | 0.01143009 | nsp6 |
| MT020881.1 | 23607 | G | T | 0.01140439 | S protein |
| MT020881.1 | 306 | T | C | 0.01134267 | nsp1 |
| MT020881.1 | 4291 | A | G | 0.01113096 | nsp3 |
| MT020881.1 | 2624 | G | A | 0.01108734 | nsp2 |
| MT020881.1 | 22673 | T | C | 0.01105964 | S protein |
| MT020881.1 | 13947 | A | G | 0.01103422 | nsp12 |
| MT020881.1 | 1442 | T | A | 0.0109349 | nsp2 |
| MT020881.1 | 5687 | T | C | 0.01074219 | nsp3 |
| MT020881.1 | 21742 | C | T | 0.01063437 | S protein |
| MT020881.1 | 7705 | A | G | 0.01031779 | nsp3 |
| MT020881.1 | 12296 | C | A | 0.01020023 | nsp8 |

DMSO P13 Lineage 2

| Genome | Position | Reference | Variant | Frequency | Gene |
|---|---|---|---|---|---|
| MT020881.1 | 21784 | T | A | 0.9815236 | S protein |
| MT020881.1 | 23525 | C | T | 0.70679713 | S protein |
| MT020881.1 | 8498 | G | T | 0.69205773 | nsp3 |
| MT020881.1 | 27761 | T | C | 0.6503799 | ORF7ab |
| MT020881.1 | 27758 | G | A | 0.65018661 | ORF7ab |
| MT020881.1 | 27760 | T | A | 0.64269198 | ORF7ab |
| MT020881.1 | 23607 | G | T | 0.44538706 | S protein |
| MT020881.1 | 23618 | A | G | 0.35962054 | S protein |
| MT020881.1 | 19547 | C | T | 0.32929351 | nsp14 |
| MT020881.1 | 29627 | C | T | 0.27142211 | ORF10 |
| MT020881.1 | 21800 | G | A | 0.21682658 | S protein |
| MT020881.1 | 76 | T | A | 0.14562166 | 5UTR |
| MT020881.1 | 22579 | T | C | 0.11323931 | S protein |
| MT020881.1 | 28892 | C | T | 0.08230895 | N protein |
| MT020881.1 | 23607 | G | A | 0.07115326 | S protein |
| MT020881.1 | 26183 | A | G | 0.05876883 | ORF3a |
| MT020881.1 | 22482 | C | T | 0.05772487 | S protein |
| MT020881.1 | 21752 | T | C | 0.0547847 | S protein |
| MT020881.1 | 26353 | C | T | 0.05468853 | E protein |
| MT020881.1 | 25671 | C | T | 0.04979287 | ORF3a |
| MT020881.1 | 26309 | C | A | 0.04283226 | E protein |
| MT020881.1 | 23606 | C | T | 0.04128542 | S protein |
| MT020881.1 | 23606 | C | G | 0.03723452 | S protein |
| MT020881.1 | 5031 | C | T | 0.03627304 | nsp3 |
| MT020881.1 | 415 | T | G | 0.03516239 | nsp1 |
| MT020881.1 | 11522 | T | G | 0.03488943 | nsp6 |
| MT020881.1 | 21846 | C | T | 0.0302586 | S protein |
| MT020881.1 | 15357 | T | C | 0.02961882 | nsp12 |
| MT020881.1 | 11559 | A | G | 0.02902944 | nsp6 |
| MT020881.1 | 11566 | T | A | 0.02760382 | nsp6 |
| MT020881.1 | 23689 | T | G | 0.02735581 | S protein |
| MT020881.1 | 21789 | C | T | 0.02434588 | S protein |
| MT020881.1 | 29284 | C | T | 0.02361305 | N protein |
| MT020881.1 | 26542 | C | T | 0.0235004 | M protein |
| MT020881.1 | 15771 | T | C | 0.02204974 | nsp12 |
| MT020881.1 | 8371 | G | A | 0.02087398 | nsp3 |
| MT020881.1 | 11471 | C | T | 0.02087146 | nsp6 |
| MT020881.1 | 13422 | T | C | 0.0204259 | nsp10 |
| MT020881.1 | 22990 | T | C | 0.02018275 | S protein |
| MT020881.1 | 8554 | T | C | 0.01914432 | nsp3 |
| MT020881.1 | 28853 | T | A | 0.01900012 | N protein |
| MT020881.1 | 26260 | T | G | 0.01885829 | E protein |
| MT020881.1 | 17748 | T | A | 0.01767842 | nsp13 |

| | | | | | |
|---|---|---|---|---|---|
| MT020881.1 | 17142 | T | C | 0.01732043 | nsp13 |
| MT020881.1 | 10451 | A | G | 0.017289 | nsp5 |
| MT020881.1 | 11379 | C | T | 0.01695565 | nsp6 |
| MT020881.1 | 15771 | T | A | 0.01684848 | nsp12 |
| MT020881.1 | 8240 | C | T | 0.01681574 | nsp3 |
| MT020881.1 | 23672 | T | C | 0.0168148 | S protein |
| MT020881.1 | 18196 | A | C | 0.01577562 | nsp14 |
| MT020881.1 | 22678 | A | G | 0.01515376 | S protein |
| MT020881.1 | 26895 | C | T | 0.01484288 | M protein |
| MT020881.1 | 13542 | T | G | 0.01481157 | nsp12 |
| MT020881.1 | 17803 | T | A | 0.01458868 | nsp13 |
| MT020881.1 | 14679 | T | A | 0.01447511 | nsp12 |
| MT020881.1 | 2473 | A | G | 0.01422042 | nsp2 |
| MT020881.1 | 7562 | T | C | 0.01391941 | nsp3 |
| MT020881.1 | 15909 | T | C | 0.01267922 | nsp12 |
| MT020881.1 | 19044 | T | A | 0.01237977 | nsp14 |
| MT020881.1 | 5389 | T | C | 0.01225432 | nsp3 |
| MT020881.1 | 13947 | A | G | 0.01178474 | nsp12 |
| MT020881.1 | 4507 | T | C | 0.01166288 | nsp3 |
| MT020881.1 | 11083 | G | T | 0.01155132 | nsp6 |
| MT020881.1 | 9812 | T | A | 0.01129496 | nsp4 |
| MT020881.1 | 1461 | A | C | 0.0112886 | nsp2 |
| MT020881.1 | 29700 | A | G | 0.01121756 | 3UTR |
| MT020881.1 | 1442 | T | G | 0.01114913 | nsp2 |
| MT020881.1 | 19896 | T | C | 0.01094635 | nsp15 |
| MT020881.1 | 1963 | T | A | 0.0108895 | nsp2 |
| MT020881.1 | 1442 | T | A | 0.01071231 | nsp2 |
| MT020881.1 | 1683 | T | C | 0.01047438 | nsp2 |
| MT020881.1 | 29665 | T | C | 0.01016612 | ORF10 |

DMSO P13 Lineage 3

| Genome | Position | Reference | Variant | Frequency | Gene |
|---|---|---|---|---|---|
| MT020881.1 | 21784 | T | A | 0.99913818 | S protein |
| MT020881.1 | 8498 | G | T | 0.99132358 | nsp3 |
| MT020881.1 | 23618 | A | G | 0.9753509 | S protein |
| MT020881.1 | 23525 | C | T | 0.87085401 | S protein |
| MT020881.1 | 27761 | T | C | 0.42961104 | ORF7ab |
| MT020881.1 | 27758 | G | A | 0.42433342 | ORF7ab |
| MT020881.1 | 27760 | T | A | 0.42191323 | ORF7ab |
| MT020881.1 | 76 | T | A | 0.15503022 | 5UTR |
| MT020881.1 | 11471 | C | T | 0.09514758 | nsp6 |
| MT020881.1 | 28096 | A | T | 0.08426392 | ORF8 |
| MT020881.1 | 25526 | G | T | 0.08277504 | ORF3a |
| MT020881.1 | 14679 | T | C | 0.05642687 | nsp12 |
| MT020881.1 | 17142 | T | C | 0.05323539 | nsp13 |
| MT020881.1 | 11522 | T | G | 0.04205989 | nsp6 |
| MT020881.1 | 10162 | T | G | 0.04130696 | nsp5 |
| MT020881.1 | 11379 | C | T | 0.04011052 | nsp6 |
| MT020881.1 | 13099 | A | G | 0.03924284 | nsp10 |
| MT020881.1 | 26234 | A | T | 0.02985769 | NA |
| MT020881.1 | 15357 | T | C | 0.02939662 | nsp12 |
| MT020881.1 | 15771 | T | C | 0.02719304 | nsp12 |
| MT020881.1 | 13542 | T | G | 0.02435639 | nsp12 |
| MT020881.1 | 18556 | A | G | 0.02395543 | nsp14 |
| MT020881.1 | 19182 | A | G | 0.0234245 | nsp14 |
| MT020881.1 | 28411 | C | T | 0.02186692 | N protein |
| MT020881.1 | 2145 | T | C | 0.02020632 | nsp2 |
| MT020881.1 | 10451 | A | G | 0.01975684 | nsp5 |
| MT020881.1 | 13422 | T | C | 0.01906588 | nsp10 |
| MT020881.1 | 22678 | A | G | 0.01849605 | S protein |
| MT020881.1 | 22114 | T | C | 0.01841935 | S protein |
| MT020881.1 | 5281 | C | T | 0.01837991 | nsp3 |
| MT020881.1 | 8554 | T | C | 0.01765766 | nsp3 |
| MT020881.1 | 19044 | T | A | 0.01739046 | nsp14 |
| MT020881.1 | 17156 | C | T | 0.01725518 | nsp13 |
| MT020881.1 | 15771 | T | A | 0.01722214 | nsp12 |
| MT020881.1 | 2473 | A | G | 0.01692242 | nsp2 |
| MT020881.1 | 20135 | T | C | 0.0156463 | nsp15 |
| MT020881.1 | 15909 | T | C | 0.01493313 | nsp12 |
| MT020881.1 | 19896 | T | C | 0.0148382 | nsp15 |
| MT020881.1 | 13409 | A | G | 0.01399786 | nsp10 |
| MT020881.1 | 22990 | T | C | 0.01398495 | S protein |
| MT020881.1 | 13947 | A | G | 0.01354887 | nsp12 |
| MT020881.1 | 6405 | T | C | 0.01342457 | nsp3 |
| MT020881.1 | 16209 | T | A | 0.01322544 | nsp12 |

| | | | | | |
|---|---|---|---|---|---|
| MT020881.1 | 7527 | T | C | 0.01242117 | nsp3 |
| MT020881.1 | 26341 | A | C | 0.01238646 | E protein |
| MT020881.1 | 1442 | T | G | 0.01235824 | nsp2 |
| MT020881.1 | 23606 | C | T | 0.01185182 | S protein |
| MT020881.1 | 20136 | A | G | 0.01163242 | nsp15 |
| MT020881.1 | 13542 | T | A | 0.01150748 | nsp12 |
| MT020881.1 | 22114 | T | A | 0.0114119 | S protein |
| MT020881.1 | 10210 | T | A | 0.01117726 | nsp5 |
| MT020881.1 | 4423 | C | T | 0.01114152 | nsp3 |
| MT020881.1 | 1442 | T | A | 0.01077618 | nsp2 |
| MT020881.1 | 4291 | A | G | 0.0105451 | nsp3 |
| MT020881.1 | 20471 | A | G | 0.0105042 | nsp15 |
| MT020881.1 | 7562 | T | C | 0.01050304 | nsp3 |
| MT020881.1 | 21931 | T | C | 0.01039987 | S protein |
| MT020881.1 | 22673 | T | C | 0.01029919 | S protein |
| MT020881.1 | 4147 | T | C | 0.01013237 | nsp3 |
| MT020881.1 | 1045 | A | G | 0.01010489 | nsp2 |
| MT020881.1 | 1683 | T | C | 0.0100613 | nsp2 |

GS-441524 P13 Lineage 1

| Genome | Position | Reference | Variant | Frequency | Gene |
|---|---|---|---|---|---|
| MT020881.1 | 23607 | G | A | 0.98462009 | S protein |
| MT020881.1 | 14033 | A | G | 0.97574124 | nsp12 |
| MT020881.1 | 13937 | T | C | 0.92317722 | nsp12 |
| MT020881.1 | 19270 | T | G | 0.63242754 | nsp14 |
| MT020881.1 | 15715 | T | G | 0.61722181 | nsp12 |
| MT020881.1 | 26537 | C | T | 0.56413544 | M protein |
| MT020881.1 | 854 | C | T | 0.5460251 | nsp2 |
| MT020881.1 | 23607 | G | T | 0.44473342 | S protein |
| MT020881.1 | 15836 | G | T | 0.38311203 | nsp12 |
| MT020881.1 | 22034 | A | G | 0.28240024 | S protein |
| MT020881.1 | 2219 | A | G | 0.15103025 | nsp2 |
| MT020881.1 | 16746 | T | C | 0.1215436 | nsp13 |
| MT020881.1 | 4775 | A | G | 0.11107922 | nsp3 |
| MT020881.1 | 28310 | C | A | 0.09177023 | N protein |
| MT020881.1 | 26461 | C | G | 0.08051303 | E protein |
| MT020881.1 | 29330 | C | A | 0.07194252 | N protein |
| MT020881.1 | 25876 | A | G | 0.06939007 | ORF3a |
| MT020881.1 | 8332 | T | C | 0.06286415 | nsp3 |
| MT020881.1 | 29397 | A | G | 0.05587388 | N protein |
| MT020881.1 | 14679 | T | C | 0.05316676 | nsp12 |
| MT020881.1 | 23855 | C | T | 0.05305704 | S protein |
| MT020881.1 | 21999 | A | C | 0.05302543 | S protein |
| MT020881.1 | 19667 | A | G | 0.05253612 | nsp15 |
| MT020881.1 | 12296 | C | A | 0.05242859 | nsp8 |
| MT020881.1 | 15222 | C | T | 0.05031391 | nsp12 |
| MT020881.1 | 26151 | C | A | 0.04550005 | ORF3a |
| MT020881.1 | 17568 | T | C | 0.04471268 | nsp13 |
| MT020881.1 | 17030 | A | G | 0.0415901 | nsp13 |
| MT020881.1 | 27383 | A | T | 0.03958908 | ORF6 |
| MT020881.1 | 13847 | A | T | 0.0386585 | nsp12 |
| MT020881.1 | 11522 | T | G | 0.03408173 | nsp6 |
| MT020881.1 | 11566 | T | A | 0.02961276 | nsp6 |
| MT020881.1 | 10451 | A | G | 0.02222583 | nsp5 |
| MT020881.1 | 12896 | C | A | 0.02054232 | nsp9 |
| MT020881.1 | 14679 | T | A | 0.01999699 | nsp12 |
| MT020881.1 | 2145 | T | C | 0.01997337 | nsp2 |
| MT020881.1 | 19018 | C | T | 0.01989334 | nsp14 |
| MT020881.1 | 4273 | T | C | 0.01970504 | nsp3 |
| MT020881.1 | 19135 | T | G | 0.01955849 | nsp14 |
| MT020881.1 | 22678 | A | G | 0.01872421 | S protein |
| MT020881.1 | 8498 | G | T | 0.01854714 | nsp3 |
| MT020881.1 | 15909 | T | C | 0.01779135 | nsp12 |
| MT020881.1 | 29459 | C | T | 0.01778775 | N protein |

| | | | | | |
|---|---|---|---|---|---|
| MT020881.1 | 17142 | T | C | 0.01775292 | nsp13 |
| MT020881.1 | 1963 | T | A | 0.0173913 | nsp2 |
| MT020881.1 | 27760 | T | A | 0.01683252 | ORF7ab |
| MT020881.1 | 20135 | T | C | 0.01676953 | nsp15 |
| MT020881.1 | 27761 | T | C | 0.0166089 | ORF7ab |
| MT020881.1 | 27758 | G | A | 0.01642702 | ORF7ab |
| MT020881.1 | 7009 | T | G | 0.01600328 | nsp3 |
| MT020881.1 | 15771 | T | A | 0.01587302 | nsp12 |
| MT020881.1 | 13542 | T | G | 0.01583226 | nsp12 |
| MT020881.1 | 493 | T | C | 0.01583189 | nsp1 |
| MT020881.1 | 7922 | T | C | 0.01563477 | nsp3 |
| MT020881.1 | 1963 | T | G | 0.01489284 | nsp2 |
| MT020881.1 | 19044 | T | A | 0.01486098 | nsp14 |
| MT020881.1 | 25679 | T | G | 0.01460791 | ORF3a |
| MT020881.1 | 2473 | A | G | 0.01437779 | nsp2 |
| MT020881.1 | 25038 | A | G | 0.01436063 | S protein |
| MT020881.1 | 2895 | A | C | 0.01430634 | nsp3 |
| MT020881.1 | 13947 | A | G | 0.01357642 | nsp12 |
| MT020881.1 | 1442 | T | A | 0.01349497 | nsp2 |
| MT020881.1 | 22114 | T | C | 0.01340849 | S protein |
| MT020881.1 | 10210 | T | A | 0.01324874 | nsp5 |
| MT020881.1 | 8554 | T | C | 0.01299714 | nsp3 |
| MT020881.1 | 28045 | C | T | 0.01196076 | ORF8 |
| MT020881.1 | 16209 | T | A | 0.01180983 | nsp12 |
| MT020881.1 | 25704 | T | A | 0.01169137 | ORF3a |
| MT020881.1 | 25923 | T | C | 0.01151635 | ORF3a |
| MT020881.1 | 4507 | T | C | 0.01151045 | nsp3 |
| MT020881.1 | 1820 | G | A | 0.0114372 | nsp2 |
| MT020881.1 | 22104 | G | A | 0.0111406 | S protein |
| MT020881.1 | 27919 | T | C | 0.01112779 | ORF8 |
| MT020881.1 | 15771 | T | G | 0.01056417 | nsp12 |
| MT020881.1 | 13542 | T | A | 0.01011405 | nsp12 |
| MT020881.1 | 7705 | A | G | 0.01005025 | nsp3 |

GS-441524 P13 Lineage 2

| Genome | Position | Reference | Variant | Frequency | Gene |
|---|---|---|---|---|---|
| MT020881.1 | 21784 | T | A | 0.99880145 | S protein |
| MT020881.1 | 8498 | G | T | 0.9946714 | nsp3 |
| MT020881.1 | 23607 | G | T | 0.97313015 | S protein |
| MT020881.1 | 1820 | G | A | 0.96437313 | nsp2 |
| MT020881.1 | 15835 | T | C | 0.86267299 | nsp12 |
| MT020881.1 | 9004 | T | C | 0.84895379 | nsp4 |
| MT020881.1 | 27761 | T | C | 0.77257954 | ORF7ab |
| MT020881.1 | 27758 | G | A | 0.77048334 | ORF7ab |
| MT020881.1 | 27760 | T | A | 0.76285384 | ORF7ab |
| MT020881.1 | 23525 | C | T | 0.24391154 | S protein |
| MT020881.1 | 11522 | T | G | 0.22255426 | nsp6 |
| MT020881.1 | 23607 | G | A | 0.19299191 | S protein |
| MT020881.1 | 16938 | A | C | 0.13187742 | nsp13 |
| MT020881.1 | 2272 | G | A | 0.11181524 | nsp2 |
| MT020881.1 | 26936 | C | T | 0.07974563 | M protein |
| MT020881.1 | 24923 | T | G | 0.07687604 | S protein |
| MT020881.1 | 13847 | A | C | 0.07452101 | nsp12 |
| MT020881.1 | 16347 | A | G | 0.05985185 | nsp13 |
| MT020881.1 | 20321 | A | C | 0.04708166 | nsp15 |
| MT020881.1 | 22316 | G | A | 0.04478094 | S protein |
| MT020881.1 | 23423 | C | T | 0.04323833 | S protein |
| MT020881.1 | 7590 | A | C | 0.04311903 | nsp3 |
| MT020881.1 | 15822 | G | T | 0.04230328 | nsp12 |
| MT020881.1 | 15357 | T | C | 0.03191291 | nsp12 |
| MT020881.1 | 3951 | A | G | 0.03004499 | nsp3 |
| MT020881.1 | 570 | A | C | 0.02907619 | nsp1 |
| MT020881.1 | 11188 | A | G | 0.02589223 | nsp6 |
| MT020881.1 | 11566 | T | A | 0.0241064 | nsp6 |
| MT020881.1 | 13542 | T | G | 0.02252356 | nsp12 |
| MT020881.1 | 22678 | A | G | 0.02174237 | S protein |
| MT020881.1 | 11462 | A | G | 0.02154725 | nsp6 |
| MT020881.1 | 13422 | T | C | 0.0213199 | nsp10 |
| MT020881.1 | 15154 | C | T | 0.02065069 | nsp12 |
| MT020881.1 | 20483 | C | T | 0.02055698 | nsp15 |
| MT020881.1 | 25807 | T | C | 0.02055158 | ORF3a |
| MT020881.1 | 15810 | C | T | 0.02010859 | nsp12 |
| MT020881.1 | 3625 | A | G | 0.0190977 | nsp3 |
| MT020881.1 | 1963 | T | A | 0.01893552 | nsp2 |
| MT020881.1 | 25494 | G | T | 0.01861668 | ORF3a |
| MT020881.1 | 17142 | T | C | 0.01785216 | nsp13 |
| MT020881.1 | 14679 | T | A | 0.01774955 | nsp12 |
| MT020881.1 | 7815 | C | T | 0.01751505 | nsp3 |
| MT020881.1 | 2606 | A | T | 0.0166769 | nsp2 |

| | | | | | |
|---|---|---|---|---|---|
| MT020881.1 | 22114 | T | C | 0.01647636 | S protein |
| MT020881.1 | 9139 | T | C | 0.01638324 | nsp4 |
| MT020881.1 | 20135 | T | C | 0.01637857 | nsp15 |
| MT020881.1 | 10620 | A | G | 0.01634541 | nsp5 |
| MT020881.1 | 19044 | T | A | 0.01554345 | nsp14 |
| MT020881.1 | 3488 | T | C | 0.01506985 | nsp3 |
| MT020881.1 | 23616 | G | A | 0.01489821 | S protein |
| MT020881.1 | 13180 | T | G | 0.01466964 | nsp10 |
| MT020881.1 | 13947 | A | G | 0.01452311 | nsp12 |
| MT020881.1 | 17886 | T | C | 0.01446979 | nsp13 |
| MT020881.1 | 1963 | T | G | 0.01426919 | nsp2 |
| MT020881.1 | 23997 | C | T | 0.01382896 | S protein |
| MT020881.1 | 8554 | T | C | 0.0136557 | nsp3 |
| MT020881.1 | 19896 | T | C | 0.01299523 | nsp15 |
| MT020881.1 | 4824 | C | A | 0.01259129 | nsp3 |
| MT020881.1 | 4291 | A | G | 0.01258318 | nsp3 |
| MT020881.1 | 163 | A | G | 0.01251371 | 5UTR |
| MT020881.1 | 25704 | T | A | 0.01251176 | ORF3a |
| MT020881.1 | 21307 | G | A | 0.0124141 | nsp16 |
| MT020881.1 | 11750 | C | T | 0.01223634 | nsp6 |
| MT020881.1 | 14318 | C | A | 0.01201861 | nsp12 |
| MT020881.1 | 25923 | T | C | 0.0116545 | ORF3a |
| MT020881.1 | 6405 | T | C | 0.01122931 | nsp3 |
| MT020881.1 | 25201 | A | G | 0.01121186 | S protein |
| MT020881.1 | 22673 | T | C | 0.01099333 | S protein |
| MT020881.1 | 1442 | T | G | 0.01091405 | nsp2 |
| MT020881.1 | 11653 | C | T | 0.01089833 | nsp6 |
| MT020881.1 | 1442 | T | A | 0.0107641 | nsp2 |
| MT020881.1 | 16209 | T | A | 0.01045179 | nsp12 |
| MT020881.1 | 25807 | T | A | 0.01008204 | ORF3a |
| MT020881.1 | 27602 | G | A | 0.01006486 | ORF7ab |
| MT020881.1 | 23606 | C | T | 0.01002036 | S protein |

GS-441524 P13 Lineage 3

| Genome | Position | Reference | Variant | Frequency | Gene |
|---|---|---|---|---|---|
| MT020881.1 | 21784 | T | A | 0.99903577 | S protein |
| MT020881.1 | 8498 | G | T | 0.99462597 | nsp3 |
| MT020881.1 | 15715 | T | G | 0.99267399 | nsp12 |
| MT020881.1 | 10162 | T | G | 0.98864644 | nsp5 |
| MT020881.1 | 15814 | G | A | 0.98817656 | nsp12 |
| MT020881.1 | 23606 | C | T | 0.98408053 | S protein |
| MT020881.1 | 14019 | T | C | 0.97722862 | nsp12 |
| MT020881.1 | 27758 | G | A | 0.9620389 | ORF7ab |
| MT020881.1 | 27761 | T | C | 0.96185128 | ORF7ab |
| MT020881.1 | 27760 | T | A | 0.95415853 | ORF7ab |
| MT020881.1 | 14679 | T | A | 0.0398022 | nsp12 |
| MT020881.1 | 23525 | C | T | 0.03777943 | S protein |
| MT020881.1 | 27383 | A | T | 0.03677112 | ORF6 |
| MT020881.1 | 15357 | T | C | 0.02878457 | nsp12 |
| MT020881.1 | 27635 | C | T | 0.02653687 | ORF7ab |
| MT020881.1 | 10451 | A | G | 0.02397105 | nsp5 |
| MT020881.1 | 11566 | T | A | 0.02188366 | nsp6 |
| MT020881.1 | 8554 | T | C | 0.02180137 | nsp3 |
| MT020881.1 | 13542 | T | G | 0.02058724 | nsp12 |
| MT020881.1 | 13422 | T | C | 0.01933156 | nsp10 |
| MT020881.1 | 11566 | T | G | 0.01862146 | nsp6 |
| MT020881.1 | 28087 | C | T | 0.0182658 | ORF8 |
| MT020881.1 | 2145 | T | C | 0.01755889 | nsp2 |
| MT020881.1 | 22678 | A | G | 0.01502097 | S protein |
| MT020881.1 | 26234 | A | T | 0.01494185 | NA |
| MT020881.1 | 22990 | T | C | 0.0139671 | S protein |
| MT020881.1 | 13947 | A | G | 0.01393862 | nsp12 |
| MT020881.1 | 19044 | T | A | 0.01391148 | nsp14 |
| MT020881.1 | 15909 | T | C | 0.01371115 | nsp12 |
| MT020881.1 | 2473 | A | G | 0.01343274 | nsp2 |
| MT020881.1 | 14679 | T | C | 0.01338456 | nsp12 |
| MT020881.1 | 1442 | T | G | 0.0127143 | nsp2 |
| MT020881.1 | 16209 | T | A | 0.01266707 | nsp12 |
| MT020881.1 | 19896 | T | C | 0.01246982 | nsp15 |
| MT020881.1 | 1963 | T | G | 0.01246537 | nsp2 |
| MT020881.1 | 25704 | T | A | 0.0120219 | ORF3a |
| MT020881.1 | 3817 | C | T | 0.01176753 | nsp3 |
| MT020881.1 | 5687 | T | C | 0.01172507 | nsp3 |
| MT020881.1 | 15771 | T | A | 0.01164469 | nsp12 |
| MT020881.1 | 7527 | T | C | 0.0114486 | nsp3 |
| MT020881.1 | 20136 | A | G | 0.0112598 | nsp15 |
| MT020881.1 | 8383 | T | C | 0.0112426 | nsp3 |
| MT020881.1 | 7562 | T | C | 0.01122309 | nsp3 |

| | | | | | |
|---|---|---|---|---|---|
| MT020881.1 | 23710 | A | G | 0.0110858 | S protein |
| MT020881.1 | 29246 | A | G | 0.01084834 | N protein |
| MT020881.1 | 1442 | T | A | 0.01076683 | nsp2 |
| MT020881.1 | 4147 | T | C | 0.01070933 | nsp3 |
| MT020881.1 | 10620 | A | G | 0.01070431 | nsp5 |
| MT020881.1 | 8821 | A | G | 0.01067687 | nsp4 |
| MT020881.1 | 25201 | A | G | 0.01057748 | S protein |
| MT020881.1 | 22673 | T | C | 0.01042843 | S protein |
| MT020881.1 | 22114 | T | A | 0.01041205 | S protein |
| MT020881.1 | 13048 | T | C | 0.01009271 | nsp10 |
| MT020881.1 | 2719 | T | C | 0.01003841 | nsp2 |
| MT020881.1 | 5389 | T | C | 0.0100349 | nsp3 |

PP nsp12-V792I

| Genome | Position | Reference | Variant | Frequency | Gene |
|---|---|---|---|---|---|
| MT020881.1 | 28281 | A | G | 0.99993964 | N protein |
| MT020881.1 | 15814 | G | A | 0.99963794 | nsp12 |
| MT020881.1 | 28558 | T | A | 0.99922901 | N protein |
| MT020881.1 | 23606 | C | T | 0.99914695 | S protein |
| MT020881.1 | 8498 | G | T | 0.99899051 | nsp3 |
| MT020881.1 | 27758 | G | A | 0.99890668 | ORF7ab |
| MT020881.1 | 21784 | T | A | 0.99881762 | S protein |
| MT020881.1 | 10162 | T | G | 0.99873418 | nsp5 |
| MT020881.1 | 27761 | T | C | 0.99863589 | ORF7ab |
| MT020881.1 | 27760 | T | A | 0.99703821 | ORF7ab |
| MT020881.1 | 26105 | A | C | 0.50499163 | ORF3a |
| MT020881.1 | 27383 | A | T | 0.09900166 | ORF6 |
| MT020881.1 | 12463 | A | T | 0.07813847 | nsp8 |
| MT020881.1 | 3765 | A | C | 0.06765579 | nsp3 |
| MT020881.1 | 3096 | C | T | 0.06300866 | nsp3 |
| MT020881.1 | 13456 | A | G | 0.02473422 | nsp12 |
| MT020881.1 | 7252 | G | T | 0.02408522 | nsp3 |
| MT020881.1 | 8107 | A | G | 0.02377622 | nsp3 |
| MT020881.1 | 21390 | A | G | 0.02304147 | nsp16 |
| MT020881.1 | 8986 | C | A | 0.02194026 | nsp4 |
| MT020881.1 | 26234 | A | T | 0.02075502 | NA |
| MT020881.1 | 26233 | G | C | 0.02022106 | NA |
| MT020881.1 | 26125 | C | A | 0.01923861 | ORF3a |
| MT020881.1 | 28012 | A | C | 0.01919496 | ORF8 |
| MT020881.1 | 18596 | G | T | 0.01793543 | nsp14 |
| MT020881.1 | 21046 | G | T | 0.01786414 | nsp16 |
| MT020881.1 | 10330 | T | C | 0.01629213 | nsp5 |
| MT020881.1 | 25807 | T | G | 0.01619567 | ORF3a |
| MT020881.1 | 26116 | G | T | 0.01608998 | ORF3a |
| MT020881.1 | 11083 | G | T | 0.01601228 | nsp6 |
| MT020881.1 | 2224 | A | G | 0.0149741 | nsp2 |
| MT020881.1 | 18321 | T | C | 0.01343748 | nsp14 |
| MT020881.1 | 28254 | A | C | 0.01196448 | ORF8 |
| MT020881.1 | 26068 | G | T | 0.01187609 | ORF3a |
| MT020881.1 | 18643 | G | T | 0.01173837 | nsp14 |
| MT020881.1 | 21059 | C | A | 0.01114119 | nsp16 |
| MT020881.1 | 2380 | A | G | 0.01107778 | nsp2 |
| MT020881.1 | 11114 | G | T | 0.01103331 | nsp6 |
| MT020881.1 | 1438 | T | A | 0.01087114 | nsp2 |
| MT020881.1 | 14967 | G | T | 0.01086803 | nsp12 |
| MT020881.1 | 27126 | G | T | 0.01050176 | M protein |
| MT020881.1 | 1820 | G | A | 0.01041581 | nsp2 |
| MT020881.1 | 4369 | T | A | 0.01035291 | nsp3 |

| | | | | | |
|---|---|---|---|---|---|
| MT020881.1 | 3932 | G | T | 0.01024924 | nsp3 |
| MT020881.1 | 5594 | G | T | 0.01019872 | nsp3 |
| MT020881.1 | 11029 | G | T | 0.01004016 | nsp6 |
| MT020881.1 | 18615 | G | T | 0.01000803 | nsp14 |
| MT020881.1 | 29560 | G | T | 0.01000196 | ORF10 |

PP nsp12-S759A/V792I

| Genome | Position | Reference | Variant | Frequency | Gene |
|---|---|---|---|---|---|
| MT020881.1 | 8498 | G | T | 0.99983585 | nsp3 |
| MT020881.1 | 15814 | G | A | 0.99973562 | nsp12 |
| MT020881.1 | 14019 | T | C | 0.99927199 | nsp12 |
| MT020881.1 | 10162 | T | G | 0.99909926 | nsp5 |
| MT020881.1 | 27758 | G | A | 0.99905752 | ORF7ab |
| MT020881.1 | 23606 | C | T | 0.99901579 | S protein |
| MT020881.1 | 27761 | T | C | 0.99887709 | ORF7ab |
| MT020881.1 | 21784 | T | A | 0.99877425 | S protein |
| MT020881.1 | 27760 | T | A | 0.99742786 | ORF7ab |
| MT020881.1 | 15715 | T | G | 0.99718246 | nsp12 |
| MT020881.1 | 27383 | A | T | 0.06430849 | ORF6 |
| MT020881.1 | 3319 | A | G | 0.04825123 | nsp3 |
| MT020881.1 | 24240 | C | T | 0.04812956 | S protein |
| MT020881.1 | 7252 | G | T | 0.02410304 | nsp3 |
| MT020881.1 | 21046 | G | T | 0.0200721 | nsp16 |
| MT020881.1 | 7279 | C | A | 0.01847804 | nsp3 |
| MT020881.1 | 18596 | G | T | 0.01552273 | nsp14 |
| MT020881.1 | 13542 | T | G | 0.01444788 | nsp12 |
| MT020881.1 | 7350 | C | A | 0.01427733 | nsp3 |
| MT020881.1 | 21059 | C | A | 0.01292949 | nsp16 |
| MT020881.1 | 25807 | T | G | 0.01189834 | ORF3a |
| MT020881.1 | 3932 | G | T | 0.01171382 | nsp3 |
| MT020881.1 | 18643 | G | T | 0.01165695 | nsp14 |
| MT020881.1 | 26068 | G | T | 0.01142839 | ORF3a |
| MT020881.1 | 13903 | G | T | 0.01045839 | nsp12 |
| MT020881.1 | 18615 | G | T | 0.01006947 | nsp14 |
